# Accelerating Molecular Dynamics with Hydrogen Mass Repartitioning to Prioritize Immunogenic Neoantigens for Cancer Vaccines through Peptide–HLA Binding Free-Energy Calculations

**DOI:** 10.64898/2026.10.11.751335

**Authors:** Diego Amaya, Jorge Flórez-Suárez, Juan Diego Mora, Carlos A. Parra-López

## Abstract

Personalized neoantigen-based vaccines have progressed from concept to clinical evidence for some of the most aggressive human tumors, yet a key challenge remains: determining which candidates should be prioritized for vaccine inclusion. Public neoantigens, shared across tumor subtypes and patient populations, offer potential as off-the-shelf candidates for broad-spectrum immunotherapy. Candidate selection relies on in silico pipelines that estimate binding; however, binding affinity alone is an imperfect marker of immunogenicity. Molecular dynamics (MD) combined with endpoint free-energy methods estimates binding free energy directly from simulated trajectories, and hydrogen mass repartitioning (HMR) mitigates the computational cost of hydrogen-involving bond vibrations. This study assessed whether HMR-accelerated MD can prioritize immunogenic neoantigens through peptide–HLA binding free-energy calculations and quantified the trade-off between accuracy and computational cost. Ten immunogenic public neoantigens (seven paired with their wild-type counterparts and three frameshift neoantigens analyzed as unpaired references) were modeled in complex with their HLA-I alleles, built de novo using AlphaFold3 and propagated with a 4-fs HMR time step. Mean binding free energy was computed across ten replicas per complex, and each SNV pair was scored concordant with experimental data when the neoantigen formed a more favorable complex than its wild type. Free energies were recomputed at 100–500 ns to identify the shortest trajectory preserving reliable ranking. Free-energy calculations reproduced experimental concordance for six of seven SNV pairs and were inconclusive for TP53 R8H; frameshift neoantigens yielded comparable values. Most complexes retained both peptide termini anchored to the HLA groove throughout, and RMSD and RMSF metrics agreed with one another and with the free-energy rankings. The PIK3CA E545K (E11K) pair showed the most distinctive signature: the peptides engaged the groove almost identically over P1–P8, with the entire difference confined to the C-terminal pocket, where loss of the wild-type’s anchor also disrupted its backbone anchor. This pair showed the largest free-energy separation in the panel, and restricting analysis to bound frames confirmed a substantial electrostatic and van der Waals advantage favoring the neoantigen. These findings support HMR-accelerated binding free-energy MD as a practical refinement step for neoantigen prioritization and its application to assessing public neoantigens for cancer vaccine design.

## 1. Introduction

Personalized neoantigen-based vaccines have progressed from concept to clinical evidence for some of the most aggressive human tumors. In resected high-risk melanoma, the individualized mRNA neoantigen therapy mRNA-4157 (V940) combined with pembrolizumab prolonged recurrence-free survival compared with pembrolizumab alone in a randomized phase 2b trial, with an 18-month recurrence-free survival of 79% vs. 62% (**1**). In pancreatic ductal adenocarcinoma (a tumor lethal in the vast majority of patients) an individualized mRNA neoantigen vaccine (autogene cevumeran) induced high-magnitude neoantigen-specific T cells in half of treated patients, and those who mounted a vaccine-expanded T-cell response had substantially longer recurrence-free survival (**2**). The rationale underlying these therapies is that neoantigens arise from tumor-specific somatic mutations and are largely absent from normal tissues, so recognition of these mutation-derived antigens enables the immune system to distinguish cancer from healthy cells and is now regarded as a major determinant of the activity of clinical cancer immunotherapies (**3**). Unlike self-derived tumor-associated antigens, mutation-derived neoantigens are foreign to the host (distinguished from germline-encoded proteins and recognized as non-self) and are therefore, in principle, not constrained by central tolerance, making them attractive targets for high-avidity T cells (**4**).

The central practical problem is no longer whether neoantigens can be therapeutic, but which of the many candidates in a given tumor should be included in a vaccine. Candidate identification relies on a largely in silico pipeline: tumor and normal DNA are sequenced to identify tumor-specific mutations, and the resulting mutant peptides are scored for presentation by the patient’s human leukocyte antigen (HLA) molecules using machine-learning predictors such as NetMHCpan-4.1, which integrate binding affinity and mass-spectrometry eluted-ligand data to estimate peptide–MHC binding (5). These tools are accurate at the binding step, yet only a small fraction of predicted binders is genuinely immunogenic. In a multi-laboratory consortium benchmark (TESLA), a model built to enrich for true epitopes filtered out roughly 98% of non-immunogenic peptides while maintaining a precision above 0.70, illustrating both the magnitude of the false-positive burden in conventional prediction and the room for improvement in candidate prioritization (6).

A principal reason for this gap is that binding affinity alone is an imperfect proxy for immunogenicity. Experimentally, the *stability* of the peptide–MHC class I complex (its kinetic lifetime rather than its equilibrium affinity) is a better correlate of cytotoxic T-cell immunogenicity: across affinity-balanced peptide sets, immunogenic peptides tend to remain bound to MHC-I more stable than non-immunogenic ones, and a substantial fraction of non-immunogenic binders can be explained by unstable interactions, frequently traceable to sub-optimal anchor residues (7). Consistently, pan-specific predictors of complex stability improve epitope identification when combined with binding predictors (8). Stability, however is, a property of the three-dimensional complex and of its dynamics, which sequence-based predictors capture only indirectly.

Physics- and structure-based modeling explicitly addresses this layer. Energetic scoring of high-resolution peptide–HLA structural models has been shown to outperform non-structural approaches in predicting CD8⁺ T-cell responses, providing both improved accuracy and mechanistic insight into the determinants of immunogenicity (9). Among such approaches, molecular dynamics (MD) combined with endpoint free-energy methods (in particular, the molecular-mechanics Poisson–Boltzmann surface-area (MM/PBSA) approach) estimates the binding free energy of a complex directly from simulated trajectories, occupying a useful middle ground between fast empirical scoring and costly rigorous alchemical methods (10). Controlling the large variance of individual trajectories through ensemble averaging over independent replicas is essential for obtaining stable and reproducible MD-based binding free-energy estimates. Building on this rationale, our group previously used molecular docking and MD as a proof of concept for two HLA-A*02:01-restricted melanoma neoantigens and their wild-type counterparts, and found that the immunogenic neoantigen formed the more stable complex, remaining anchored at both peptide termini throughout the trajectory, in agreement with its in vitro immunogenicity (11). That study, however, was qualitative and limited to two neoantigen– wild-type pairs on the HLA-A*02:01 allele, leaving open the question of whether a quantitative, energy-based criterion can distinguish immunogenic from non-immunogenic neoantigens across a broader panel of MHC alleles.

The primary obstacle to applying MD at the scale required by a prioritization pipeline is computational cost, dominated by the short integration time step (typically 1–2 fs) imposed by the fastest bond vibrations involving hydrogen. Hydrogen mass repartitioning (HMR) mitigates this directly: by transferring part of the mass of heavy atoms to the hydrogens bonded to them, the highest-frequency motions are slowed, allowing the time step to be increased by up to a factor of two (to 4 fs) with no significant change in the thermodynamic or kinetic properties of the simulated system (12). In practice, this roughly halves the wall-clock time required for a given trajectory, making otherwise prohibitive ensemble simulations of many peptide–HLA complexes tractable,

Finally, unlike personal neoantigens, shared tumor neoantigens (13), tumor-associated neoepitopes reported to be immunogenic across tumor subtypes and patient populations, exhibiting HLA allele specificity, and representing putative targets for public neoantigen-based cancer immunotherapies have been reported (14). The use of these public neoantigens combined with efficient vaccine platforms would provide non-personalized “off-the-shelf” therapeutic vaccine candidates for broad-spectrum immunotherapy.

Here we test whether HMR-accelerated MD can prioritize immunogenic neoantigens using peptide–HLA binding free-energy calculations, and we quantify the trade-off between accuracy and computational cost. We simulated seven neoantigens, together with their non-immunogenic wild-type counterparts and three frameshifts public neoantigens, each in complex with the HLA class I allele, all with experimentally documented immunogenicity. The peptide–HLA complexes were built de novo with AlphaFold3 (15) (rather than by the template-based docking used in our earlier work) and propagated with a 4-fs HMR time step. For each complex, we computed the mean binding free energy over ten independent replicas (MM-PBSA enthalpy with a quasi-harmonic configurational-entropy term), and scored a pair as concordant with experiment when the immunogenic neoantigen formed the energetically more favorable complex than its wild-type sequence. The panel also included three frameshift-derived neoantigens, which, lacking a one-to-one wild-type counterpart, were simulated as unpaired immunogenic references. To determine the simulation length needed for reliable ranking, we recomputed the binding free energy at 100, 200, 300, 400, and 500 ns and identified the shortest trajectory that preserved agreement with the experimental data. The neoantigen was correctly ranked below its wild-type counterpart in six of the seven pairs, and 300 ns emerged as the best compromise between experimental agreement and computational cost, supporting HMR-accelerated binding free-energy calculation as a practical refinement step for neoantigen prioritization in personalized cancer vaccines.

## 2. Materials and Methods

### 2.1. Neoantigen Panel and HLA Restrictions

We assembled a panel of tumor neoantigens with experimentally documented CD8⁺ T-cell immunogenicity, drawn from the literature, together with the corresponding non-immunogenic wild-type peptide for each single-nucleotide-variant (SNV) neoantigen. The panel comprises two complementary sets. The first set comprises seven SNV neoantigen/wild-type pairs, in which the mutant and wild-type peptides differ by a single residue and are presented by the same HLA class I allele; these pairs constitute the validation set for the pairwise ranking analysis (Section 2.8). The second set comprises three frameshift (FS) neoantigens (in *APC*, *NPM1* and *TGFBRII* genes), which, by construction, generate entirely novel reading frames and therefore have no one-to-one wild-type counterpart; these were simulated as additional, unpaired immunogenic complexes. In total, the study comprises 17 distinct peptide–HLA complexes (7 mutant + 7 wild-type + 3 frameshift). Peptide sequences, genes, mutation types, HLA restrictions and the originating reports are given in **Table 1**. Separately from this panel, the two HLA-A*02:01-restricted neoantigen/wild-type pairs of our earlier proof-of-concept study (**11**)(ASTN1 and AKAP6, four complexes), whose in vitro CD8⁺ T-cell immunogenicity was reported by Strønen et al.,(**16**) were simulated first with the same protocol as a feasibility pilot; they are reported separately in Section 3.1 and are excluded from all panel-level statistics.

**Table 1.**
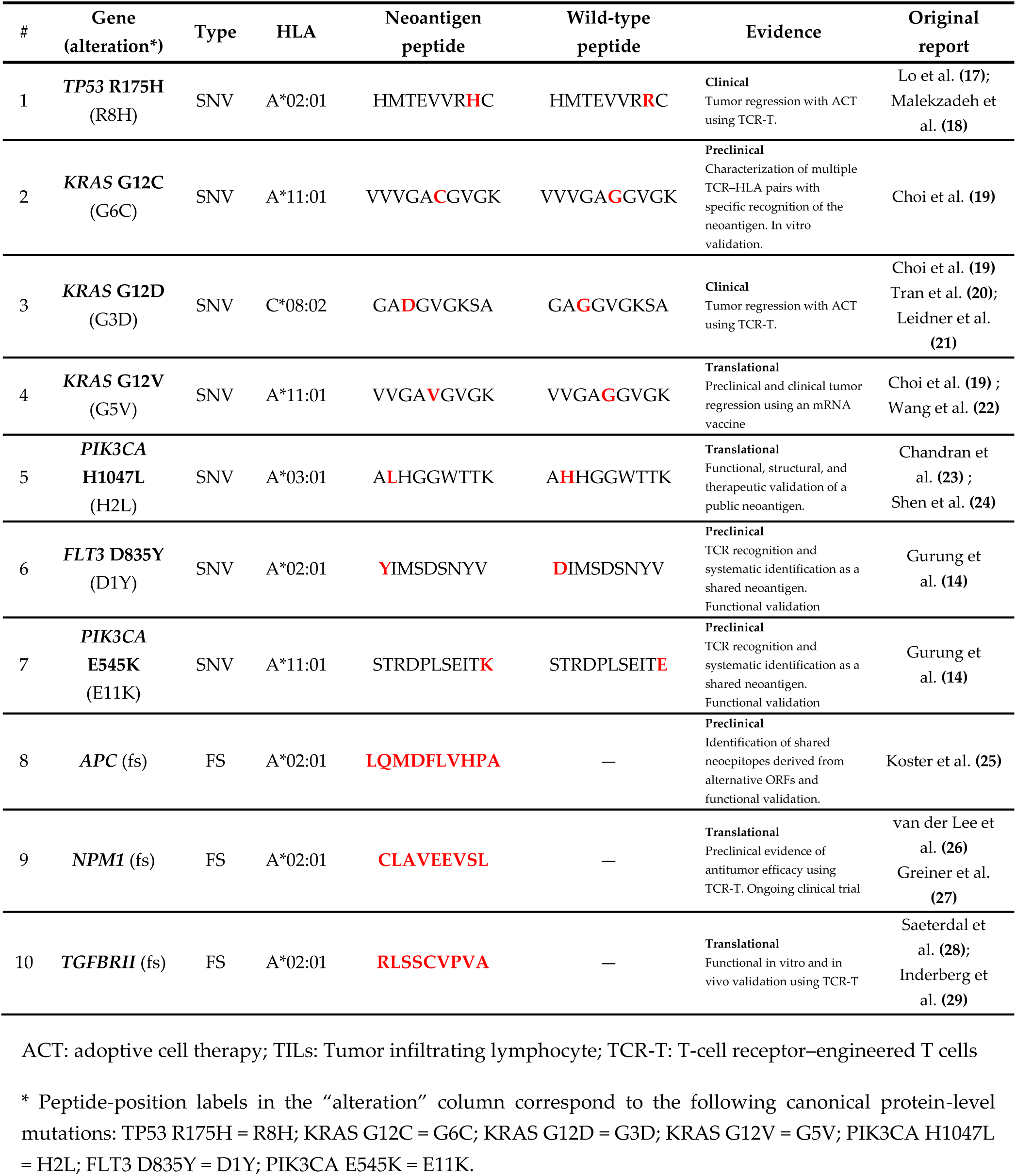
Neoantigen panel simulated in this study and their associated evidence. Mutated residues in the SNV neoantigens are shown in bold relative to the wild-type sequence. FS = frameshift; SNV = single-nucleotide variant.

Each neoantigen was simulated in complex with the HLA class I allele under which its immunogenicity had been reported. Every modeled complex contained the HLA class I heavy chain (α1–α2–α3 domains), β2-microglobulin, and the 9–11-residue peptide.

### 2.2. Modeling of the Peptide–HLA Complexes

Three-dimensional models of each peptide–HLA complex were generated *de novo* with AlphaFold3 (**15**) via the AlphaFold Server (https://alphafoldserver.com) using the server’s default setting. The HLA class I heavy-chain sequence, the β2-microglobulin sequence, and the neoantigen (or wild-type) peptide sequence were provided as a single multi-chain prediction. The top-ranked predicted model (by the server’s default ranking score) was retained for each complex and used as the starting structure for molecular dynamics (MD) simulations.

### 2.3. System Preparation and Hydrogen Mass Repartitioning

System preparation was automated using VMD/psfgen (**30**). For each predicted complex, histidine residues were assigned the neutral δ-tautomer (HSD). Topologies and coordinates were generated per chain using the CHARMM36m all-atom protein force field (**31**).

The defining methodological feature of this work was introduced at this stage: hydrogen mass repartitioning (HMR) (**12**) was applied to the solute using the psfgen hmassrepart operation, which increases each hydrogen atom’s mass to 3.024 amu and subtracts the corresponding mass from its bonded heavy atom, leaving the total molecular mass unchanged. This slows the highest-frequency X–H stretching motions, permitting a doubled integration time step (4 fs instead of the conventional 2 fs) without loss of stability, which roughly halves the wall-clock cost of each MD simulation (**12**).

The HMR-repartitioned complex was then solvated in a rectangular box of modified TIP3P water (**32,33**) with a minimum solute (wall padding of 20 Å). The system was neutralized by adding Na⁺/Cl⁻ counter-ions.

### 2.4. Molecular Dynamics Protocol

All simulations were run with GPU-resident NAMD3 (**34**). Non-bonded interactions used a 12 Å real-space cutoff, with a switching function applied from 10 Å and a 14 Å pair-list distance. Long-range electrostatics were treated with the particle-mesh Ewald (PME) method (**35**) (1.0 Å grid spacing). Electrostatics were evaluated every two steps, and short-range non-bonded forces every step. CHARMM 1–4 electrostatic scaling was used. The protocol consisted of three phases:

1. **Energy minimization:** The solvated, ionized system was energy minimized for 20,000 conjugate-gradient steps (1 fs reference time step, flexible covalent bonds, no thermostat or barostat).
2. **Restrained equilibration (4 fs, HMR):** Equilibration was performed at 310 K and 1 atm in the NPT ensemble. Temperature was maintained with a Langevin thermostat (damping coefficient 1 ps⁻¹), and pressure with a Langevin piston barostat (target 1 atm, period 400 fs, decay 200 fs) (**36**). Positional harmonic restraints were applied to the solute and released in two successive stages: first, all protein atoms were restrained (force constant 1.0 kcal mol⁻¹ Å⁻²) for 5 ns; then restraints were confined to the protein backbone and progressively relaxed over five sub-stages (force constant scaled 1.0 → 0.8 → 0.6 → 0.4 → 0.2 kcal mol⁻¹ Å⁻², 5 × 4 ns), for 20 ns of backbone-restrained equilibration.
3. **Production (4 fs, HMR):** Unrestrained production dynamics were then run under the same NPT, 4 fs HMR conditions for 500 ns per replica.

### 2.5. Replicas

To average the sensitivity of endpoint free-energy estimates to initial conditions, each of the 17 complexes was simulated in 10 independent replicas, yielding 170 production trajectories in total. Each replica was equilibrated independently from the same AlphaFold3 model of the complex, with initial atomic velocities assigned randomly from the Maxwell–Boltzmann distribution at 310 K, yielding ten statistically independent trajectories per complex. Ensemble averaging across replicas was used to control the large trajectory-to-trajectory variance of single-trajectory MM-PBSA estimates.

### 2.6. Binding Free-Energy Calculation

Peptide–HLA binding free energies were computed using the molecular-mechanics Poisson–Boltzmann surface-area (MM-PBSA) method (**10**), with a single-trajectory protocol, in which the complex, the receptor (HLA heavy chain + β2-microglobulin), and the ligand (peptide) are all extracted from the same trajectory.

#### 2.6.1. Topology conversion

Because the trajectories were generated using a CHARMM force field, the NAMD PSF topologies and CHARMM36m parameters were converted to AMBER ChamberParm topologies using ParmEd (AmberTools (**37**)), preserving the CMAP, Urey–Bradley, and NBFIX terms. Separate solvated, dry-complex, receptor, and ligand topologies were generated for each system.

#### 2.6.2. Trajectory preparation

For each replica and each analysis window, the first 50 ns of each production trajectory was discarded as equilibration. Water and ions were stripped with CPPTRAJ (**38**), and snapshots were extracted at regular intervals, yielding approximately 400 frames per replica for the MM-PBSA enthalpy calculation.

#### 2.6.3. Poisson–Boltzmann settings

MM-PBSA was performed with MMPBSA.py (**39**), using sander for energy evaluation and the settings required for CHARMM-derived (ChamberParm) topologies: prmtop-assigned radii (radiopt = 0) with the linear non-polar solvation model (inp = 1); solute interior dielectric ε_in_ = 4 and solvent dielectric ε_out_ = 80; ionic strength 0.15 M; grid scale 2.0 (≈0.5 Å spacing); solvent probe radius 1.4 Å; and fill ratio 4.0. The MM-PBSA binding enthalpy ΔH was decomposed into van der Waals (VDWAALS), electrostatic (EEL), polar solvation (EPB), and non-polar solvation (ENPOLAR) contributions.

#### 2.6.4. Configurational entropy

The binding free energy was computed as ΔG_bind_ = ΔH_MM-PBSA_ − TΔS_conf_, where the solute configurational entropy change on binding, ΔS_conf_, was estimated using the quasi-harmonic (QH) approximation (**40**) from the mass-weighted covariance matrix of the Cα coordinates of the complex, receptor and ligand, computed independently for each replica. The resulting estimate, denoted ΔG_bind_, is the binding free energy reported throughout the Results.

#### 2.6.5. Ensemble averaging

For each complex, ΔG_bind_ was computed per replica, and the ensemble estimate was reported as the mean of the ten per-replica values ± the standard error of the mean (SEM) across replicas. The difference between a neoantigen and its wild-type counterpart was quantified as ΔΔG = ΔG(neoantigen) − ΔG(wild-type), with its SEM propagated from the two ensembles.

### 2.7. Determination of the Optimal Simulation Time

To identify the shortest trajectory that still yields reliable rankings, ΔG_bind_ was recomputed for every complex using progressively longer portions of the production run (100, 200, 300, 400, and 500 ns) by truncating the analysis window and repeating the full MM-PBSA + quasi-harmonic-entropy procedure of Section 2.6 for each window. Convergence was assessed per complex by examining the plateau of the ensemble ΔG_bind_ across the longest windows: a complex was considered converged when its ensemble ΔG_bind_ over the last three windows (300, 400, and 500 ns) agreed to within 2 SEM (i.e., the spread across these three windows did not exceed twice the mean per-window SEM). The optimal simulation length was defined as the shortest window on this common plateau that also preserved the experimental agreement of the pairwise rankings (Section 2.8) across the panel; this trade-off between accuracy and computational cost identified 300 ns as the working simulation time.

### 2.8. Ranking Criterion against Experimental Immunogenicity

For each of the seven SNV pairs, the experimental ground truth is that the neoantigen is immunogenic whereas its wild-type counterpart is not. A pair was scored as in agreement with experiment when the neoantigen formed the energetically more favorable complex, i.e., ΔG_bind_(neoantigen) < ΔG_bind_(wild-type) (equivalently, ΔΔG < 0). The three frameshift neoantigens, which have no wild-type comparator, were not included in the pairwise agreement metric; their ΔG_bind_ values are reported as additional immunogenic reference points.

### 2.9. Peptide–HLA Stability Analysis

As a structural, dynamics-based complement to the energetic ranking, the persistence of peptide anchoring within the binding groove was monitored across the ten replicas of each complex. Each peptide terminus was represented by its terminal anchor region (the terminal residue together with its adjacent neighbor residue), and two quantities were computed per frame relative to the HLA binding groove: the minimum heavy-atom distance between the anchor region and the groove, and the number of anchor–groove heavy-atom contacts (atom pairs within 4.5 Å), normalized to a reference value averaged over the first 5 ns. A terminus was scored as transiently unbound when its minimum distance exceeded 5.0 Å or its contact fraction fell below 0.30 of the reference, sustained for ≥2 ns and ignoring the first 50 ns. From these events, we derived, per complex, the fraction of replicas showing an unbinding event, the first-unbinding times, the fraction of analyzed time spent unbound, and a Kaplan–Meier survival curve (with 95% Greenwood confidence intervals) describing the retention of each terminus over the ensemble.

### 2.10. Peptide Conformational Analysis: RMSD Distribution and Per-Residue Fluctuation

The peptide conformational analyses were run on all 17 complexes (170 replicas). Every quantity reported in Section 3.6 is computed on the peptide alone; the HLA heavy chain and β2-microglobulin are excluded from all reported values and enter only through the trajectory that produced them.

#### 2.10.1. Analysis window and superposition convention

The analysis window was frames 418-4166 of each production trajectory, i.e. 50–500 ns (3749 frames per replica, 37,490 frames per complex), the first 50 ns being discarded as equilibration in accordance with the MM-PBSA convention of Section 2.6.2. Note that the conformational analyses therefore use the full 50-500 ns trajectory, whereas the binding free energies reported in Sections 3.1–3.3 use the 300 ns working window; the two are complementary readouts of the same simulations rather than the same calculation at two lengths.

Each frame was superposed on the peptide’s own Cα atoms before any measurement. This choice is deliberate and determines what the analysis can see: with the peptide superposed on itself, rigid-body motion of the peptide inside the groove (sliding, rocking, and in particular a terminus lifting out of the pocket) is removed by the fit, and the resulting RMSD (Root Mean Square Deviation), RMSF (Root Mean Square Fluctuation) and distributions report the peptide’s internal conformational change only. Displacement of the peptide relative to the groove is measured separately, and by a different criterion, in the terminal-stability analysis of Section 2.9.

#### 2.10.2. RMSD distribution

RMSD was computed for each frame against frame 1 of that replica’s own production trajectory (t ≈ 0) using the peptide backbone atoms N, Cα, C and O.

The RMSD distribution of each complex was estimated by pooling all 37,490 frames (10 replicas × 3749 frames) and rendering them as a normalized histogram (70 bins over a common range) overlaid with a Gaussian kernel-density estimate. All panels of a given figure share one common x-scale and one common y-scale so that they can be compared directly.

Because frames within a replica are strongly autocorrelated, mutant/wild-type inference was carried out at the replica level: a two-sided Mann–Whitney U test on the ten mutants against the ten wild-type per-replica mean RMSD values. The Kolmogorov–Smirnov distance D between the two pooled distributions is additionally reported as a purely descriptive measure of how different the two full distributions are; its p-value would be meaningless at the frame level and is not given.

#### 2.10.3. Per-residue RMSF

Per-residue RMSF was measured about the ensemble mean structure, following the CPPTRAJ’S recipe: superpose every frame on the first frame, compute the average structure, re-superpose every frame on that average, and only then measure the fluctuation of each residue about its mean position. RMSF was computed independently within each replica and then averaged over the ten replicas, with error bars given as the standard error of that mean across replicas.

Because the panel contains 9-, 10- and 11-residue peptides, results are displayed on two length-independent coordinate systems. The terminal plot uses eight positions, the first four (P1–P4) and the last four (PΩ−3, PΩ−2, PΩ−1, PΩ), so that peptides of different length align on their termini, which are the anchor regions. The core plot uses five central slots M1–M5, taken as positions 3–7 for a 9-mer and 4–8 for an 11-mer; a 10-mer has no single central residue, so the four residues symmetric about the 5.5 midpoint (positions 4–7) are assigned to slots M2–M5 and slot M1 is left empty.

### 2.11. Peptide–HLA Interaction-Network Analysis

To resolve the origin of the free-energy difference in the pair that behaved most distinctively (PIK3CA E545K, STRDPLSEIT**K**/STRDPLSEIT**E** on HLA-A*11:01; Section 3.7), the complete residue-level interaction network between peptide and HLA was reconstructed from the 20 trajectories of that pair (2 peptides × 10 replicas × 4166 frames).

#### 2.11.1. Scope and interaction types

The receptor scope was restricted to HLA heavy-chain residues 1–180, the α1+α2 platform that forms the peptide-binding groove.

For every frame, each peptide residue was scored against each of the 180 groove residues for five interaction types. Distances use heavy atoms only; hydrogens enter only through the hydrogen-bond angle.

Contacts reported in Section 3.7 are defined following this geometric criterion: any heavy atom of the peptide residue within 4.5 Å of any heavy atom of the HLA residue.

Gas-phase molecular-mechanics interaction energies were decomposed over the same residue pairs. The screened electrostatic (electrostatics interaction into a solvent) is reported, using a distance-dependent dielectric ε(r) = 4r together with a Debye–Hückel factor exp (−r/7.86 Å) at the pipeline’s 0.15 M ionic strength. The screened value is a qualitative descriptor used for ranking which electrostatic interaction pairs survive in water, not a thermodynamic quantity.

#### 2.11.2. Conditioning windows

The wild-type peptide of this pair spends roughly 57% of its trajectory with its C-terminus detached (Section 3.5). Any contact difference measured over all frames would therefore be partly a restatement of the unbinding result rather than an explanation of it. Every comparison was consequently computed in four conditioning windows: onset (first 2 ns, before any unbinding event in either system), early (first 25 ns, matched initial-state window), bound (50–500 ns, restricted to frames in which the peptide C-terminal region satisfies the bound criterion of Section 2.9), and all (50–500 ns, every frame, i.e. the ensemble the MM-PBSA calculation used). The conclusion is identical in all four. Unless stated otherwise, the values reported in Section 3.7 are from the bound window, which is the conservative choice because it removes the detached frames and therefore asks whether the wild-type interface is weaker even while it is engaged.

### 2.12. Software and Statistics

Structure prediction used AlphaFold3 (**15**); system preparation used VMD/psfgen (VMD 1.9.4) (**30**); simulations used NAMD3 (**34**) with the CHARMM36m force field (**31**); free-energy calculations used AmberTools 25 (**37**) (ParmEd, CPPTRAJ (**38**), sander, MMPBSA.py (**39**)). Trajectory processing used CPPTRAJ (**38**); entropy estimation, convergence, unbinding, conformational and interaction-network analyses, and figure generation were performed with in-house Python scripts. Uncertainties are reported as SEM across the ten replicas unless otherwise stated.

Throughout the study the replica, not the frame, is the unit of statistics. Group comparisons between a neoantigen and its wild-type counterpart use the two-sided Mann–Whitney U test on the ten per-replica values of each system; where many interactions are tested against the same replicas, p-values are corrected by the Benjamini–Hochberg procedure and reported as false-discovery rates (q). The Kolmogorov–Smirnov distance D is used only as a descriptive measure of the separation between two full distributions and is reported without a p-value.

## 3. Results

We simulated the full panel of 17 peptide–HLA complexes (seven SNV neoantigen/wild-type pairs and three unpaired FS neoantigens, see **Table 1**), each in ten independent replicas propagated for 500 ns with a 4-fs hydrogen-mass-repartitioned (HMR) time step, for a total of 170 production trajectories and ≈85 µs of aggregate sampling. All 170 trajectories ran to completion without numerical instability and were analyzed using the single-trajectory MM-PBSA binding-free-energy protocol of Section 2.6; the modeled peptides remained accommodated in the binding groove for most of each trajectory in most complexes (Section 3.6), indicating that doubling the integration time step did not destabilize the peptide–HLA complexes.

### 3.1. A pilot test on the proof-of-concept complexes establishes the feasibility of the protocol

Before committing to the full panel, we applied the entire procedure to the four peptide–HLA complexes of our earlier proof-of-concept study(**11**), which were the first systems simulated with the HMR protocol and served as a pilot test. These are two HLA-A*02:01-restricted neoantigen/wild-type pairs whose in vitro CD8⁺ T-cell readout is known and is external to the present work: ASTN1 (wild-type KPYGLDWAEL, neoantigen KLYGLDWAEL, a proline-to-leucine substitution at P2), whose neoantigen is immunogenic, and AKAP6 (wild-type WLIDMESLV, neoantigen WLIDMKSLV, a glutamate-to-lysine substitution at P6), whose neoantigen is not (**11**). The two pairs therefore probe the calculation in opposite directions: it should resolve a free-energy preference for the ASTN1 pair and should not resolve one for the AKAP6 pair. The four complexes were prepared, simulated and analyzed exactly as described in Section 2 (HMR with a 4-fs time step, ten independent 500 ns replicas per complex, MM-PBSA with a quasi-harmonic entropy term, and the same 100, 200, 300, 400 and 500 ns analysis windows), so their values are directly comparable with those of the panel. Ensemble binding free energies over the five windows are given in **Table 2**.

**Table 2.**
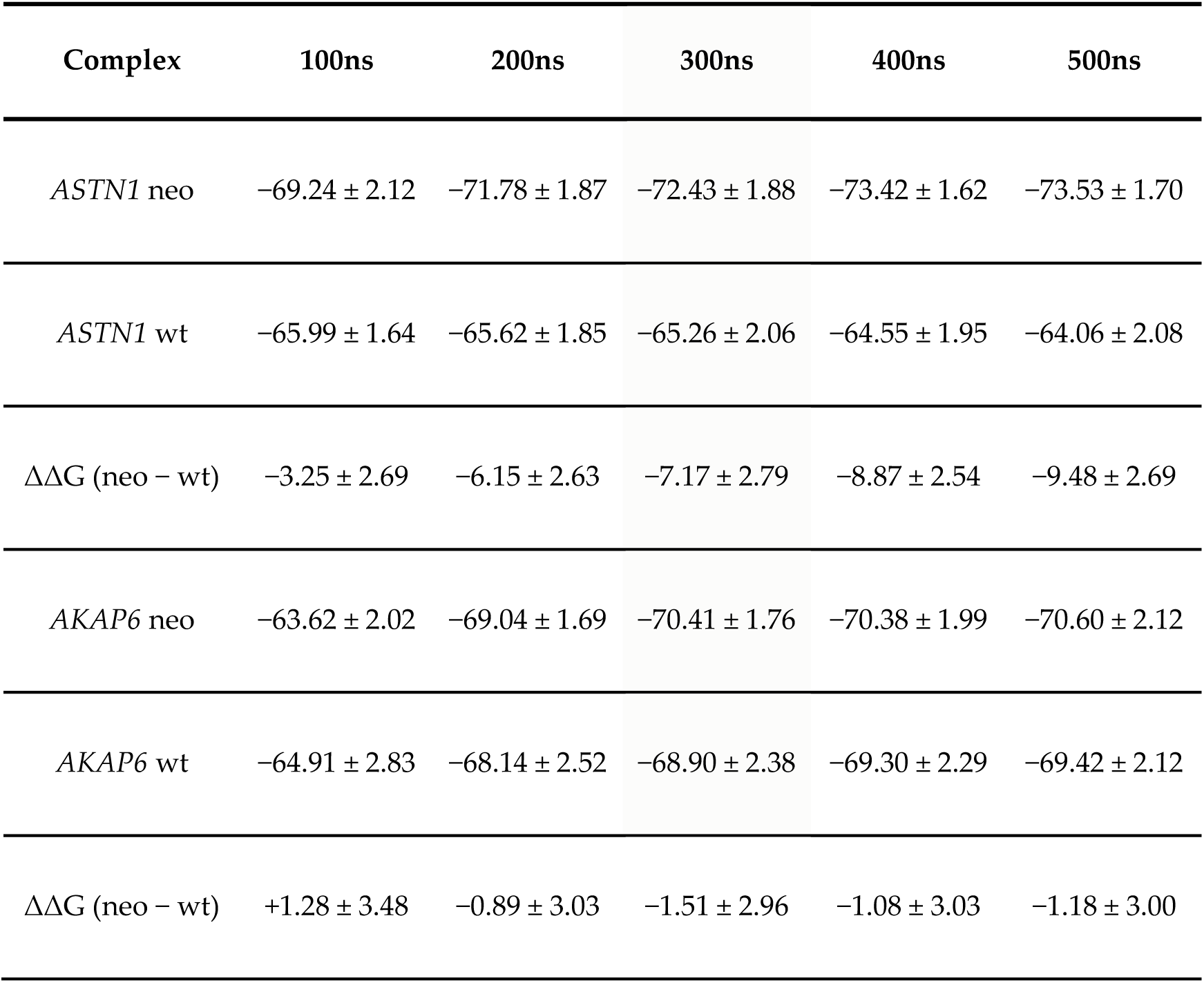
Ensemble binding free energy ΔG_bind_ of the four pilot complexes over the five analysis windows (mean ± SEM across ten replicas), and the neoantigen−wild-type difference ΔΔG of each pair. All values are in kcal/mol. The two pairs are the HLA-A*02:01-restricted complexes of our earlier proof-of-concept study: ASTN1 (wild-type KPYGLDWAEL, neoantigen KLYGLDWAEL; P2L), whose neoantigen is immunogenic for CD8⁺ T cells in vitro, and AKAP6 (wild-type WLIDMESLV, neoantigen WLIDMKSLV; E6K), whose neoantigen is not (**11**).

The pilot reproduced the experimental distinction, and did so with increasing clarity as the analysis window was lengthened. For the immunogenic ASTN1 pair, ΔΔG is negative at every window and grows steadily in magnitude, from −3.25 ± 2.69 kcal/mol at 100 ns (1.2 SEM, i.e. unresolved) to −7.17 ± 2.79 kcal/mol at 300 ns (2.6 SEM) and −9.48 ± 2.69 kcal/mol at 500 ns (3.5 SEM). For the non-immunogenic AKAP6 pair, no free-energy preference is resolved at any window: ΔΔG is +1.28 ± 3.48 kcal/mol at 100 ns and lies between −0.89 and −1.51 kcal/mol from 200 ns onward, in every case smaller in magnitude than its own SEM (≤ 0.5 SEM). The calculation therefore resolves a difference only for the pair for which the in vitro assay resolves one.

Three conclusions were drawn from this pilot, and they determined how the main study was run. First, the HMR protocol is numerically viable for this class of system: all forty replicas (four complexes × ten replicas × 500 ns, ≈20 µs of aggregate sampling) ran to completion at a 4-fs time step without numerical instability. Second, a 100 ns analysis window is too short: it is the only window at which the ASTN1 separation falls below twice its SEM, and the only one at which the sign of the (unresolved) AKAP6 difference is reversed with respect to every longer window, whereas from 300 ns onward both pairs are stable in sign and in magnitude and all four complexes satisfy the plateau criterion of Section 2.7 over the 300, 400 and 500 ns windows (largest spread 1.2 kcal/mol, against a tolerance of at least 3.5 kcal/mol). Third, the absolute binding free energies are of the expected order (ASTN1 neoantigen, −72.43 ± 1.88 kcal/mol at 300 ns), within the range later found for the neoantigens of the panel (−61.6 to −75.9 kcal/mol; Table 3). On this basis the protocol was applied unchanged to the 17 complexes of the panel.

**Table 3.**
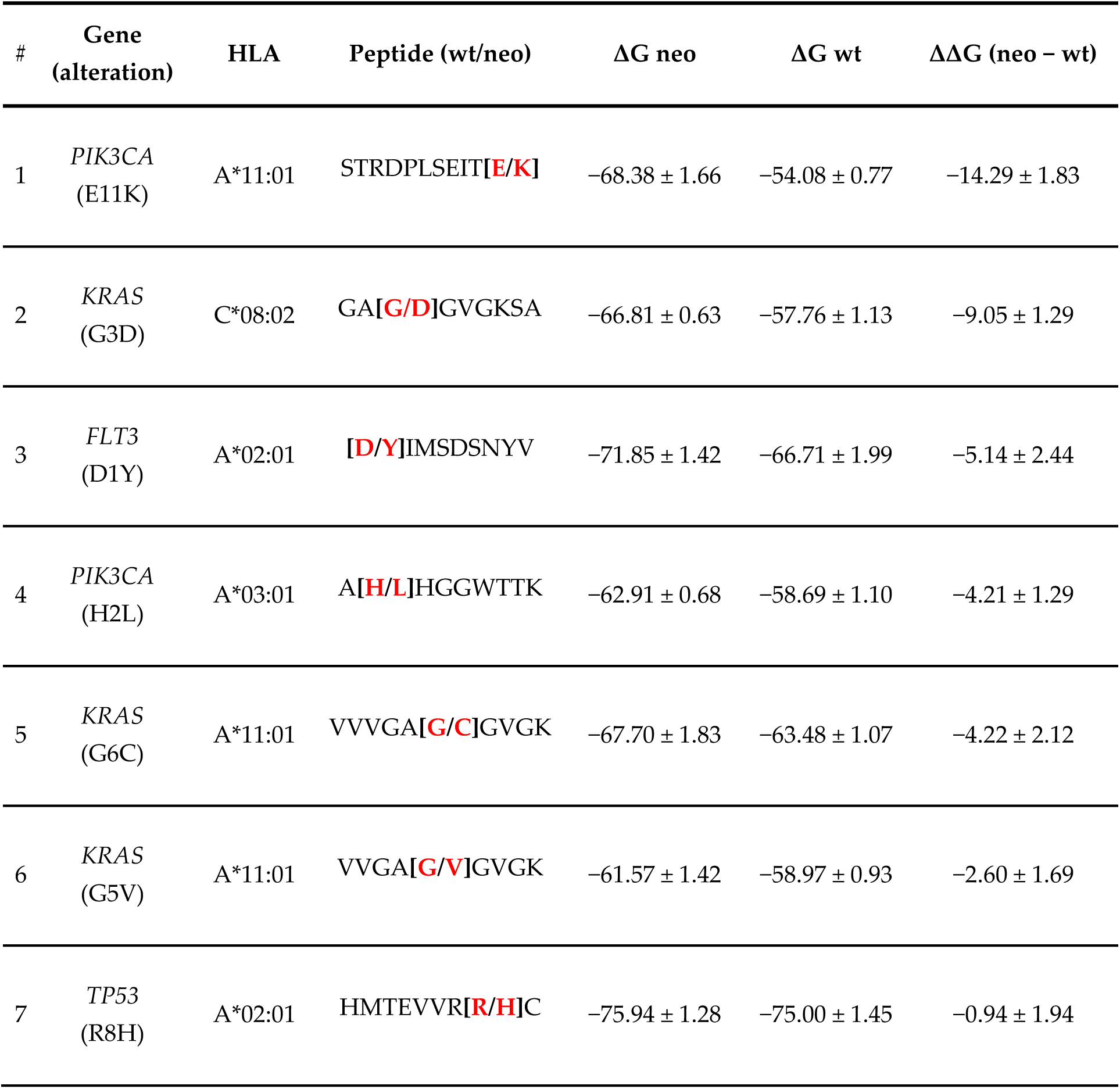
Ensemble binding free energy ΔG_bind_ over the 300 ns window (mean ± SEM across ten replicas), and the neoantigen−wild-type difference ΔΔG for each SNV pair. All values are in kcal/mol. More negative ΔG indicates more favorable binding; ΔΔG < 0 indicates that the neoantigen binds more favorably than its wild-type counterpart (concordant with experimental data).

### 3.2. Binding free energy discriminates neoantigens from their wild-type counterparts

For each pair of single-nucleotide variants (SNVs), the experimental ground truth is that the mutant peptide is immunogenic, whereas its wild-type counterpart is not (see **Table 1**). We therefore investigated whether the calculated binding free energy is more favorable (i.e., more negative) for the neoantigen than for the wild-type peptide. This is expressed as ΔΔG = ΔG(neoantigen) − ΔG(wild-type) < 0 (see Section 2.8). Ensemble binding free energies over the 300 ns window are summarized in **Table 3**.

In all seven pairs, the neoantigen was assigned the more favorable (more negative) binding free energy, so the sign criterion ΔΔG < 0 is satisfied throughout. The seven pairs separate cleanly into two groups by the magnitude of the discrimination. For six pairs, the neoantigen is favored by a substantial margin, with ΔΔG ranging from −2.6 kcal/mol (KRAS G5V) to −14.3 kcal/mol (PIK3CA E11K). Three of these are resolved well beyond statistical noise: PIK3CA E11K (−14.3, ≈7.8 SEM), KRAS G3D (−9.0, ≈7.0 SEM) and PIK3CA H2L (−4.2, ≈3.3 SEM); FLT3 D1Y (−5.1) and KRAS G6C (−4.2, ≈2.0 SEM) are resolved at the ≈2-SEM level; and KRAS G5V (−2.6, ≈1.5 SEM) points in the concordant direction, with margins just below twice its SEM. The remaining pair, TP53 R8H, is the single exception: its ΔΔG of −0.9 ± 1.9 kcal/mol is statistically indistinguishable from zero (it is the only pair whose |ΔΔG| is smaller than its own SEM), so the calculation does not resolve a free-energy preference between the mutant HMTEVVR**H**C and wild-type HMTEVVR**R**C peptides. We therefore conclude that the HMR-accelerated free-energy calculation reproduces the experimental neoantigen/wild-type ranking for six of the seven pairs and is inconclusive for the TP53 R8H pair. This 6/7 outcome extends, on a larger and more diverse panel and using a quantitative energetic criterion, the qualitative discrimination reported for two HLA-A*02:01 melanoma neoantigens in our earlier proof-of-concept study (**11**).

It is worth noting that the TP53 R8H complexes are also the two with the most favorable absolute binding free energies in the whole panel (ΔG ≈ −75 to −76 kcal/mol for both mutant and wild-type).

### 3.3. Frameshift neoantigens bind with favorable free energy

The three frameshift neoantigens lack one-to-one wild-type comparators and were therefore excluded from the pairwise metric; their binding free energies are reported as additional immunogenic reference points (see **Table 4**). At the 300 ns window, the APC (LQMDFLVHPA) and NPM1 (CLAVEEVSL) frameshift peptides bind HLA-A*02:01 with ΔG_bind_ of −67.05 ± 1.56 and −66.22 ± 1.17 kcal/mol, respectively, and the TGFBRII (RLSSCVPVA) peptide binds with −60.54 ± 1.71 kcal/mol. These values fall within the range spanned by the immunogenic SNV neoantigens (−61.6 to −75.9 kcal/mol; Table 2): the APC and NPM1 peptides fall squarely within it and the TGFBRII peptide is marginally below the least-favorable SNV neoantigen (KRAS G5V, −61.6), placing the three experimentally validated frameshift epitopes among the more favorably ranked complexes in the panel.

**Table 4.** Ensemble binding free energy ΔG_bind_ for the three unpaired frameshift neoantigens over the 300 ns window (mean ± SEM across ten replicas), kcal/mol.

| Gene | HLA | Peptide | $\Delta G_{\text{bind}}$ |
| --- | --- | --- | --- |
| APC | A*02:01 | LQMDFLVHPA | $-67.05 \pm 1.56$ |
| NPM1 | A*02:01 | CLAVEEVSL | $-66.22 \pm 1.17$ |
| TGFBR2 | A*02:01 | RLSSCPVA | $-60.54 \pm 1.71$ |

### 3.4. Effect of simulation length and choice of a 300 ns working window

To determine the shortest trajectory that still yields reliable rankings, we recomputed ΔG_bind_ for each complex using progressively longer segments of the production run (100, 200, 300, 400, and 500 ns; Section 2.7). The neoantigen−wild-type differences are summarized in **Table 5**.

**Table 5.** Neoantigen−wild-type ΔΔG (mean ± SEM, kcal/mol) for each SNV pair as a function of analysis-window length. Negative throughout indicates that the experimental ranking is preserved across all windows.

| Pair (alteration) | 100ns | 200ns | 300ns | 400ns | 500ns |
| --- | --- | --- | --- | --- | --- |
| *PIK3CA* (E11K) | $-14.58 \pm 1.43$ | $-13.90 \pm 1.51$ | $-14.29 \pm 1.83$ | $-13.39 \pm 2.05$ | $-12.89 \pm 2.19$ |
| *KRAS* (G3D) | $-8.55 \pm 0.96$ | $-8.74 \pm 1.13$ | $-9.05 \pm 1.29$ | $-9.25 \pm 1.34$ | $-9.20 \pm 1.29$ |
| *FLT3* (D1Y) | $-1.46 \pm 1.92$ | $-3.73 \pm 1.99$ | $-5.14 \pm 2.44$ | $-4.91 \pm 2.48$ | $-5.16 \pm 2.53$ |
| *PIK3CA* (H2L) | $-2.08 \pm 1.69$ | $-2.92 \pm 1.20$ | $-4.21 \pm 1.29$ | $-4.96 \pm 1.65$ | $-5.48 \pm 1.82$ |
| *KRAS* (G6C) | $-4.78 \pm 2.21$ | $-4.07 \pm 2.24$ | $-4.22 \pm 2.12$ | $-4.37 \pm 2.00$ | $-4.49 \pm 1.92$ |
| *KRAS* (G5V) | $-2.10 \pm 1.49$ | $-2.31 \pm 1.55$ | $-2.60 \pm 1.69$ | $-3.17 \pm 1.89$ | $-3.73 \pm 1.81$ |
| *TP53* (R8H) | $-1.25 \pm 2.37$ | $-1.54 \pm 1.87$ | $-0.94 \pm 1.94$ | $-1.01 \pm 2.04$ | $-1.17 \pm 1.81$ |

Two observations follow. First, the qualitative outcome is remarkably stable with simulation length: every ΔΔG is negative in every window, so the 6/7 pairwise agreement (and the single TP53 R8H exception) is already established at 100 ns and does not change as the trajectories are extended. The ranking is therefore not the limiting factor in choosing a simulation length. Second, the magnitudes converge unevenly across the panel: most pairs reach a stable ΔΔG by ∼200–300 ns (e.g., PIK3CA E11K, KRAS G6C, KRAS G3D, FLT3 D1Y, TP53 R8H), whereas two pairs (KRAS G5V and PIK3CA H2L) continue to drift toward more negative ΔΔG out to 500 ns, indicating that the absolute separation for those two is not fully converged even at the longest window.

The working simulation time was therefore set not by the rankings (which are window-independent) but by the convergence of the per-complex ensemble free energy. All 17 complexes satisfy the plateau criterion of Section 2.7: the ensemble ΔG_bind_ values over the last three windows (300, 400, and 500 ns) agree to within 2 SEM, with the largest spread across those three windows being 1.6 kcal/mol (for the APC frameshift) and smaller for every other complex. In other words, 300 ns is the shortest window on the converged plateau common to the whole panel. Moving from 300 ns to 500 ns changes neither any pairwise ranking nor the ensemble means by more than ∼1.6 kcal/mol, whereas extending each replica from 300 to 500 ns costs 67% more simulation time. Therefore, 300 ns represent the best trade-off between agreement with the experimental data and computational cost, and was adopted as the working window for the rest of this work.

### 3.5. Replica-to-replica dispersion of the free-energy estimates

For each complex, the per-replica binding free energies and their evolution with simulation length are summarized in the **Supplementary Fig. S1 panels**. Each panel shows, on the left, the ensemble mean ΔG_bind_ ± SEM versus simulation time, and, on the right, the ten individual per-replica trajectories alongside the ensemble mean. These panels are not intended as a formal convergence analysis but to convey the spread of the free-energy estimate across the ten replicas, i.e., the replica-to-replica reproducibility of a single-trajectory MM-PBSA value.

The dispersion is substantial and varies markedly across complexes. It is quantified as the standard deviation of the ten per-replica ΔG_bind_ values (averaged over the five windows). The replica spread ranges from ≈1.9 to ≈5.8 kcal/mol, and the full per-replica range within a single complex reaches up to ≈25 kcal/mol (still ≈21 kcal/mol even at the 300ns working window). Grouped by magnitude (see **Table 6**), the complexes fall into four bands: a very-high-spread band (SD ≥ 5.5 kcal/mol: KRAS G6C neoantigen, FLT3 wild-type, TGFBRII frameshift), a high band (≈4.5–5.5: APC frameshift, PIK3CA E11K neoantigen, TP53 R8H neoantigen), a medium band (≈3.0–4.5: KRAS G5V neoantigen, TP53 R8H wild-type, FLT3 neoantigen, PIK3CA H2L wild-type, NPM1 frameshift, KRAS G3D wild-type, KRAS G6C wild-type), and a low band (≤ 3.0: KRAS G5V wild-type, PIK3CA H2L neoantigen, PIK3CA E11K wild-type, KRAS G3D neoantigen). The last of which is the most reproducible complex in the panel (SD ≈ 1.9 kcal/mol).

**Table 6.** Replica dispersion of the binding free energy, expressed as the standard deviation (SD, kcal/mol) of the ten per-replica ΔG_bind_ values averaged across the five analysis windows. Complexes are grouped into four spread bands. Per-complex panels are shown in Supplementary Fig. S1.

| Spread band<br>(SD range) | Complex | Replica SD |
| --- | --- | --- |
| Very high<br>($\geq 5.5$) | *KRAS* (G6C) neo - A*11:01 | 5.8 |
|  | *FLT3* (D1Y) wt - A*02:01 | 5.7 |
|  | *TGFBR2* (fs) - A*02:01 | 5.6 |
| High<br>(4.5–5.5) | *APC* (fs) - A*02:01 | 5.2 |
|  | *PIK3CA* (E11K) neo - A*11:01 | 5.1 |
|  | *TP53* (R8H) neo - A*02:01 | 4.6 |
| Medium<br>(3.0–4.5) | *KRAS* (G5V) neo - A*11:01 | 4.4 |
|  | *TP53* (R8H) wt - A*02:01 | 4.3 |
|  | *FLT3* (D1Y) neo - A*02:01 | 4.3 |
|  | *PIK3CA* (H2L) wt - A*03:01 | 3.8 |
|  | *NPM1* (fs) - A*02:01 | 3.8 |
|  | *KRAS* (G3D) wt - C*08:02 | 3.2 |
|  | *KRAS* (G6C) wt - A*11:01 | 3.1 |
| Low<br>($\leq 3.0$) | *KRAS* (G5V) wt - A*11:01 | 3.0 |
|  | *PIK3CA* (H2L) neo - A*03:01 | 2.9 |
|  | *PIK3CA* (E11K) wt - A*11:01 | 2.5 |
|  | *KRAS* (G3D) neo - C*08:02 | 1.9 |

Two patterns are discernible. The clearest is allele- and mutation-linked: for all three HLA-A*11:01-restricted KRAS/PIK3CA pairs the neoantigen is more dispersed than its wild-type counterpart (KRAS G6C, 5.8 vs 3.1; PIK3CA E11K, 5.1 vs 2.5; KRAS G5V, 4.4 vs 3.0), suggesting that the introduced mutation increases the conformational heterogeneity sampled by these HLA-A*11:01 binders; the opposite or no trend is seen for the HLA-A*02:01, HLA-A*03:01, and HLA-C*08:02 pairs, so this observation should be treated as suggestive rather than general, given only three HLA-A*11:01 pairs. The second pattern links dispersion to peptide mobility: the two frameshift peptides with the highest spread, TGFBRII (5.6) and APC (5.2), are also the two complexes with the most frequent terminal unbinding (Section 3.5), consistent with weaker peptide anchoring broadening the conformational ensemble and thereby inflating the between-replica variance of the end-point free-energy estimate. The overarching message of **Figure S1**, however, is method-level: because a single replica can deviate from the ensemble mean by several kcal/mol (and individual replicas span up to ∼25 kcal/mol within one complex), a single-trajectory MM-PBSA value is not a reliable estimator on its own, and the ten-replica ensemble averaging adopted here is essential for stable ranking.

### 3.6. Peptide–terminal stability in the binding groove

As a structural complement to the energetic ranking, we monitored the persistence of peptide anchoring by tracking transient unbinding of each peptide terminus from the HLA groove across the ten replicas of each complex (Section 2.9). Ensemble statistics are summarized in **Table 7**; per-complex distance and contact time series along with Kaplan–Meier retention curves are provided as **Supplementary Fig. S2**.

**Table 7.**
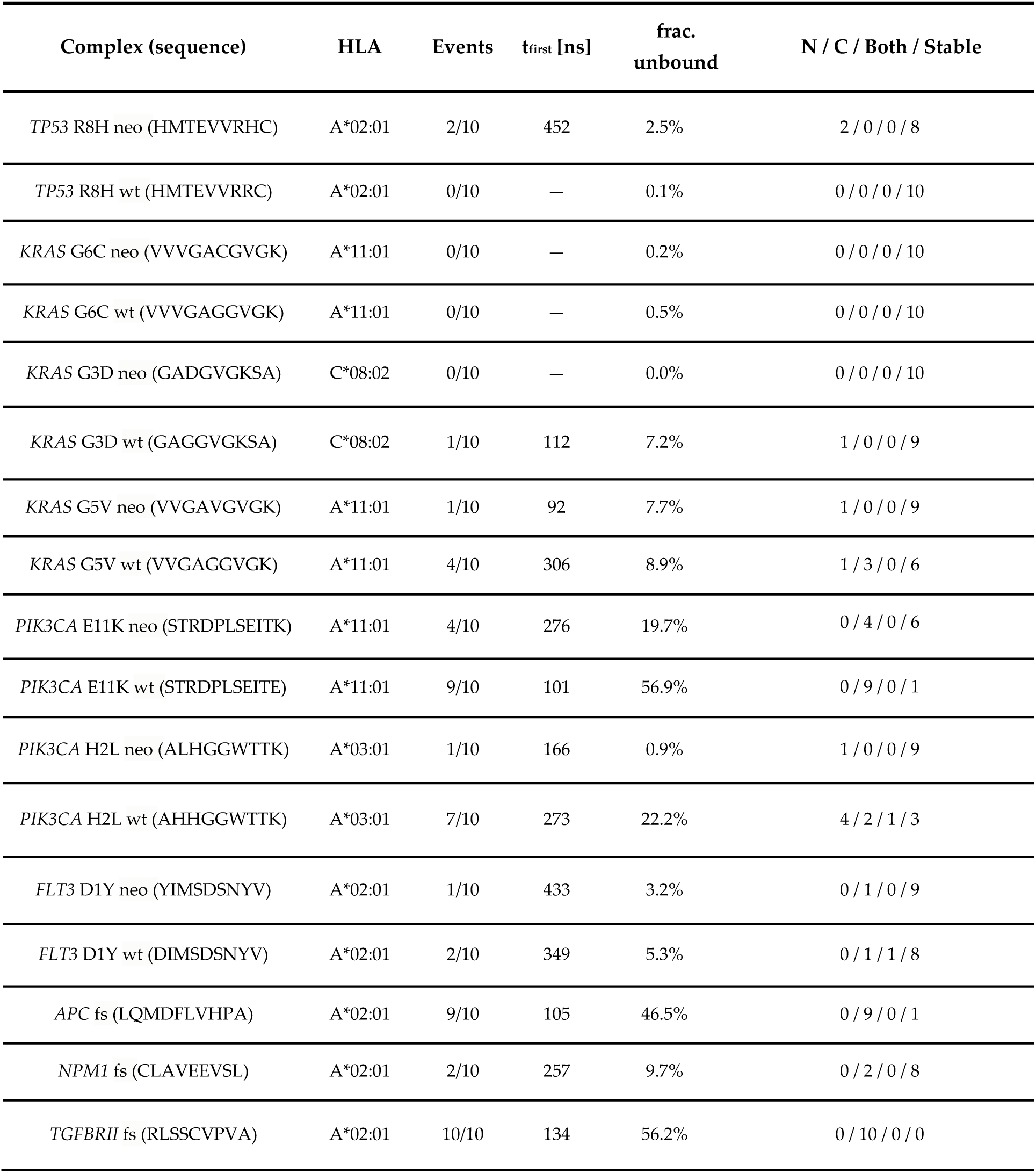
Peptide–terminal unbinding statistics across the ten replicas of each complex (criterion: minimum anchor–groove distance and normalized contact count; scope: terminal residue + adjacent neighbor; Section 2.9). "Events" = replicas with at least one sustained unbinding event; "frac. unbound" = mean fraction of analyzed time spent unbound; breakdown = replicas with N-terminal-only / C-terminal-only / both / neither (stable) events. Per-complex panels are shown in **Supplementary Fig. S2**.

Most complexes kept both peptide termini anchored for essentially the entire analyzed trajectory: eleven of the seventeen complexes showed sustained unbinding in ≤2 of ten replicas and spent <10% of the analyzed time unbound. Four of those (KRAS G6C neoantigen and wild-type, KRAS G3D neoantigen, TP53 R8H wild-type) were fully stable, with no unbinding events in any replica. When unbinding did occur, it overwhelmingly involved the C-terminus rather than the N-terminus. At the other extreme, four complexes were markedly unstable: the TGFBRII frameshift peptide (events in 10/10 replicas, 56% of analyzed time unbound), the PIK3CA E11K wild-type (9/10, 57%), the APC frameshift peptide (9/10, 47%), and the PIK3CA H2L wild-type (7/10, 22%). In these, excursions were confined to the C-terminus for the TGFBRII and APC frameshift peptides and the E11K wild-type, but occurred predominantly at the N-terminus for the H2L wild-type (4 N-only / 2 C-only / 1 both; **Table 7**).

For the SNV pairs, terminal stability tracks the energetic ranking in the expected direction for several pairs: the PIK3CA E11K and PIK3CA H2L wild-type peptides are much more prone to unbinding than their neoantigen counterparts (E11K, 9/10 vs 4/10; H2L, 7/10 vs 1/10), mirroring the more favorable ΔG_bind_ of the neoantigens. The two frameshift peptides with the highest unbinding frequency are also the highest-dispersion complexes in Section 3.5, reinforcing the link between conformational lability and free-energy noise. We note that this structural stability measure is a qualitative, complementary readout: it is concordant with, but not a substitute for, the energetic ranking, and it does not by itself resolve the TP53 R8H pair (the R8H neoantigen shows rare N-terminal excursions in 2/10 replicas, whereas its wild-type is fully stable), consistent with the small energetic separation found for that pair in Section 3.2.

### 3.7 The neoantigen is generally the more conformationally constrained partner

The free-energy ranking in Sections 3.3–3.4 indicates which complex is more favorable but does not explain the underlying reasons for those results. We therefore characterized the conformational behavior of each peptide directly from the same trajectories, using two complementary readouts (Section 2.10): the RMSD distribution, which shows how many distinct conformations a peptide visits and how often, and the per-residue RMSF, which shows where along the sequence the peptide is pinned down and where it is mobile. Both are computed on the peptide alone, over the 50–500 ns window, with each frame superposed on the peptide’s own Cα atoms. That fitting convention is important for interpretation: it removes rigid-body motion of the peptide inside the groove, so these measures describe the peptide’s internal shape change and by construction, are not a measure of unbinding (which is quantified separately in Section 3.6).

#### 3.7.1. RMSD distribution

**Figure 1** shows the RMSD distribution of all 17 peptides, each panel pooling 37,490 frames, on a common pair of axes so that panels can be compared directly. The corresponding statistics are given in **Table 8**.

**Figure 1.**
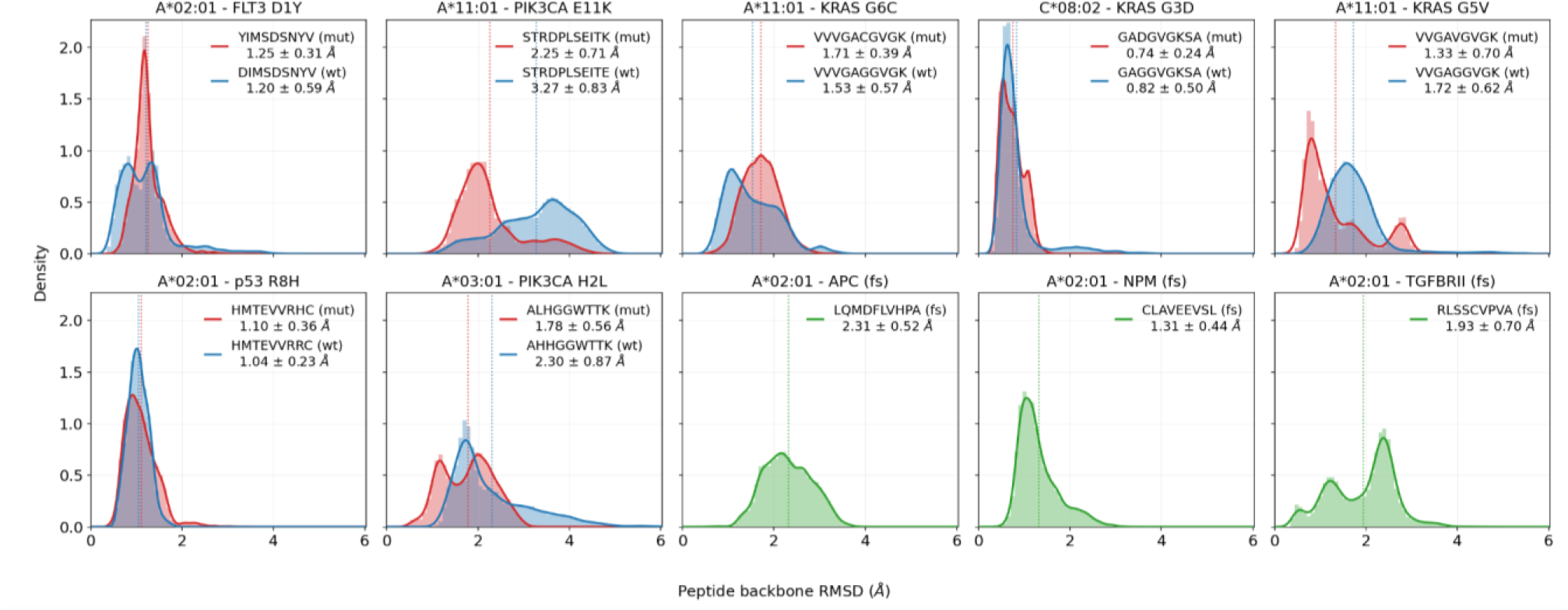
Peptide RMSD distribution of the 17 peptide–HLA complexes. Each panel plots the probability density of the peptide backbone (N, Cα, C, O) RMSD sampled over 50–500 ns, pooling all ten replicas (37,490 frames per curve); the shaded histogram is the raw frequency distribution and the solid line its kernel-density estimate. RMSD is measured against frame 1 of each replica’s own production run after superposition on the peptide’s own Cα atoms, so the abscissa reports internal conformational change. Panels 1–7 (top row and first two of the bottom row) show the seven SNV pairs, with the neoantigen in red and the wild-type peptide in blue; panels 8-10 show the three unpaired frameshift neoantigens in green. Dotted vertical lines mark the mean of each distribution, and the inset legends give mean ± SD in Å. All panels share one common x-scale and one common y-scale, so broad, right-shifted distributions (for example the PIK3CA H2L neoantigen, which is clearly bimodal) stand out directly against sharp, low-RMSD ones (for example KRAS G3D).

**Table 8.** Peptide backbone RMSD over the 50–500 ns window for the seven SNV pairs (mean ± SD over the 37,490 pooled frames of each complex). Δ is the neoantigen−wild-type difference; a negative Δ means the neoantigen is internally more constrained. p is a two-sided Mann–Whitney U test on the seven mutants against the seven wild-type per-replica mean RMSD values (the replica being the unit of statistics, Section 2.10.2). D is the Kolmogorov–Smirnov distance between the two pooled distributions, reported as a descriptive measure of their separation only. Alterations are given in peptide-position notation, as in Table 1. Bold p values are below 0.05.

| Pair<br>(allele – alteration) | Neoantigen<br>RMSD [Å] | Wild-type<br>RMSD [Å] | $\Delta$ (neo – wt)<br>RMSD | Mann-Whitney<br>$p$ | KS $D$ |
| --- | --- | --- | --- | --- | --- |
| A*11:01 – *PIK3CA* E11K | 2.25 ± 0.71 | 3.27 ± 0.83 | –1.02 | **0.002** | 0.569 |
| A*03:01 – *PIK3CA* H2L | 1.78 ± 0.56 | 2.30 ± 0.87 | –0.52 | 0.076 | 0.252 |
| A*11:01 – *KRAS* G5V | 1.33 ± 0.70 | 1.72 ± 0.62 | –0.39 | **0.038** | 0.452 |
| C*08:02 – *KRAS* G3D | 0.74 ± 0.24 | 0.82 ± 0.50 | –0.08 | 0.623 | 0.080 |
| A*02:01 – *FLT3* D1Y | 1.25 ± 0.31 | 1.20 ± 0.59 | +0.05 | 0.571 | 0.254 |
| A*02:01 – *TP53* R8H | 1.10 ± 0.36 | 1.04 ± 0.23 | +0.06 | 0.571 | 0.125 |
| A*11:01 – *KRAS* G6C | 1.71 ± 0.39 | 1.53 ± 0.57 | +0.18 | 0.241 | 0.269 |

The neoantigen has the lower mean RMSD in four of the seven pairs. Only two pairs reach significance at the replica level, and both of them favors the neoantigen: PIK3CA E11K (Δ = −1.02 Å, p = 0.002) and KRAS G5V (Δ = −0.39 Å, p = 0.038). PIK3CA H2L shows the same direction with a larger effect but does not reach the threshold (Δ = −0.52 Å, p = 0.076). The RMSD distribution therefore supports the same ordering as the free energy, but on its own it has much weaker discriminative power.

The exception is PIK3CA E11K, where the difference is not one of degree but of kind. The wild-type STRDPLSEITE shows a broad multimodal-like distribution, indicating that it alternates among several distinct conformational states, whereas the neoantigen STRDPLSEITK remains in a single, narrower state; the Kolmogorov–Smirnov distance of 0.569 is the largest in the panel. This is also the pair with the largest ΔΔG (Section 3.2) and the largest asymmetry in terminal stability (Section 3.6), and it is the pair we dissect in Section 3.7. Among the frameshift neoantigens, NPM1 CLAVEEVSL is narrow and low (1.31 ± 0.44 Å), while APC LQMDFLVHPA (2.31 ± 0.52 Å) and TGFBRII RLSSCVPVA (1.93 ± 0.70 Å, visibly multimodal) are broad, consistent with the terminal-stability ranking in **Table 7**.

#### 3.7.2. Per-residue fluctuation

The per-residue picture is more informative than the whole-peptide one. **Figure 2** shows the Cα RMSF for the first four and last four residues of each peptide, aligned at the termini so that peptides of different lengths are comparable at their anchor positions; **Figure 3** shows the central residues on the M1–M5 coordinate.

**Figure 2.**
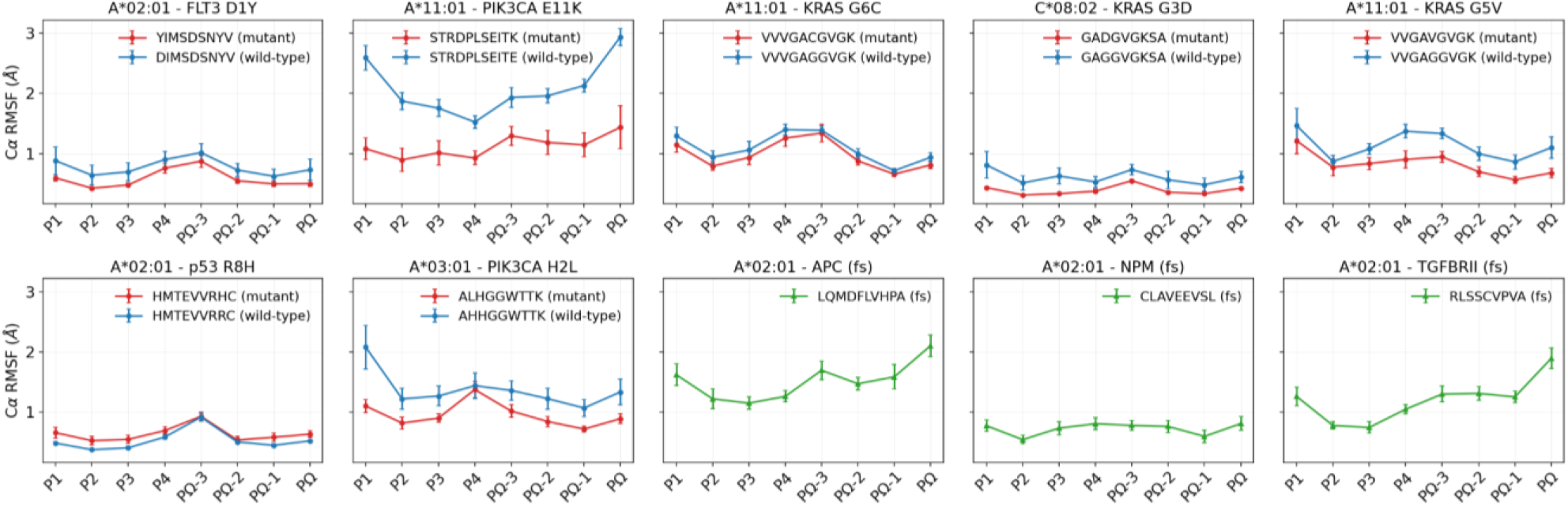
Cα RMSF of the four N-terminal (P1–P4) and four C-terminal (PΩ−3, PΩ−2, PΩ−1, PΩ) residues across all 17 peptides, with one panel per SNV pair or unpaired frameshift neoantigen. Because peptides of 9, 10, and 11 residues are aligned at their termini, the leftmost and rightmost points of every panel correspond to the same structural positions, namely the anchor regions of the groove. Neoantigens are shown in red, wild-type peptides in blue, and frameshift neoantigens in green. Each point is the mean over the ten replicas of the per-replica RMSF, measured relative to that replica’s ensemble-mean structure over 50–500 ns; error bars are the SEM across replicas. All panels share a common y-axis. Higher values indicate greater residue mobility.

**Figure 3.**
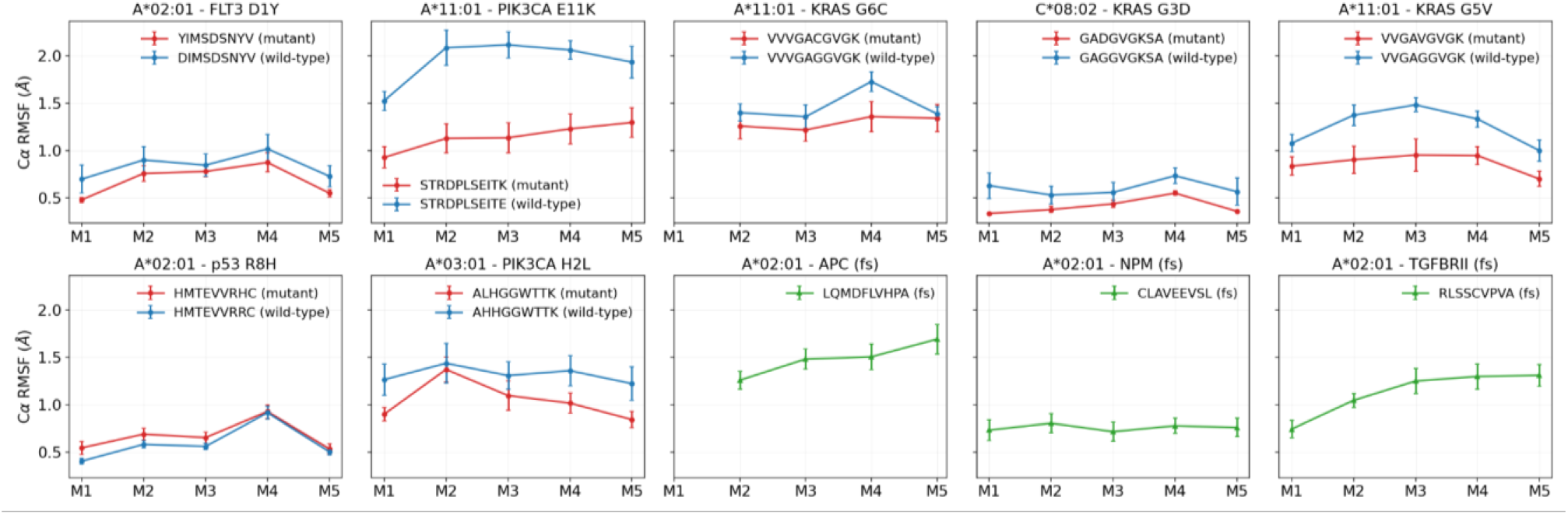
Cα RMSF of the central residues of all 17 peptides, plotted across five length-independent slots M1– M5 (positions 3–7 for a 9-mer, 4–8 for an 11-mer, and 4–7 mapped onto M2–M5 for a 10-mer, whose even length leaves M1 empty; Section 2.10.3). Colors, error bars, averaging, and the shared y-axis are as in Figure 2. These positions point away from the groove floor and toward the T-cell receptor, so this panel describes the mobility of the surface that a T cell would read.

**Figures 2 and 3** read together show that the difference between a neoantigen and its wild-type counterpart is not confined to a single region of the peptide. In six of the seven pairs, the neoantigen curve lies below the wild-type curve at every plotted position: all eight terminal slots of **Figure 2**, from P1 to PΩ, and all five core slots M1–M5 of **Figure 3**. Since the two panels together cover every residue of every peptide, the statement can be made per residue: the neoantigen has the lower Cα RMSF in 57 of the 66 residue-level comparisons in the seven pairs, and all nine exceptions belong to the single TP53 R8H pair, in which the reversal is likewise uniform across the sequence but negligible in size (+0.01 to +0.17 Å). Neighboring residues of one peptide are strongly correlated, so these 66 comparisons are not 66 independent observations, and the pair-level count of six out of seven remains the meaningful one; but the fact that the sign is uniform within each pair, rather than averaging out across a mixture of positions, is what the two figures add to the whole-peptide numbers. The magnitude of the effect varies over an order of magnitude across pairs and is small in several of them: KRAS G6C, the weakest concordant pair, separates by only 0.05–0.37 Å depending on position, and FLT3 D1Y by 0.07–0.29 Å, differences that are visible in the panels but comparable to the replica-to-replica error bars. PIK3CA E11K is in a different regime altogether. Its wild-type STRDPLSEITE is between 1.5 and 2.4 times as mobile as the neoantigen at every one of the eleven residues, and the smallest per-residue difference in that pair (0.60 Å, at P4) exceeds in magnitude the largest per-residue difference found anywhere in five of the other six pairs, PIK3CA H2L (0.98 Å at P1) being the only one to come close.

Beyond the direction of the effect, the two panels indicate where along the peptide the effect acts. First, the fluctuation profile has a characteristic shape that recurs across the panel. It is clearest in the twelve 9-mers, for which the terminal and core slots refer to non-overlapping residues: averaged over those twelve peptides, the mean Cα RMSF runs 0.98, 0.65, 0.72, 0.90, 0.89, 0.98, 0.76, 0.67, and 0.84 Å from P1 to P9. There are therefore two rigidity minima, at P2 and P8, each one residue from an end, and the fluctuation rises again in three places: at the two chain ends and in the middle of the peptide. Eleven of the twelve 9-mers have their single most rigid residue at P2 or P8, and across the full panel of 17, the most rigid of the eight terminal slots is one of these two positions in 14 cases. This means that the peptides are, in other words, held near both extremities and comparatively free in between, which is how a class I ligand is bound. The N-terminal minimum has a direct structural reading: P2 is one of the two canonical anchor positions, and its side chain is buried in the groove’s B pocket (**13,41**). The C-terminal minimum needs a more careful interpretation, because it falls at P8 rather than at PΩ, the anchor residue itself. The last residue is in fact more mobile than its neighbor (0.81 against 0.66 Å, even when the three peptides that release a terminus are excluded) which is what one expects of a chain terminus: it is held by its own anchoring side chain and by its C-terminal carboxylate, but unlike an internal residue it has a backbone neighbor on one side only. Second, in **Figure 3**, the largest core-window difference falls exactly on the mutated residue in all three KRAS pairs, which are the only ones whose substitution lies within that window: M1 = P3 for G3D (−0.29 Å), M3 = P5 for G5V (−0.53 Å) and M4 = P6 for G6C (−0.37 Å). The same holds at the other end of the peptide for FLT3 D1Y in **Figure 2**, where the substitution at P1 produces that pair’s largest difference (−0.29 Å). Replacing a glycine, which has no side chain and confers maximal backbone freedom, with a residue that has one rigidifies that residue more than any other. PIK3CA E11K is, again, an interesting case. Its substitution is at P11, yet the two largest differences in the pair are at P1 (−1.51 Å) and at P11 (−1.50 Å), in equal measure, and P1 lies ten residues from the mutation. However, surprisingly, no unbinding event manifests at the N-terminal of the wild-type peptide (**Table 7**).

Across all residues of the peptide, the neoantigen is less flexible than its wild-type counterpart in six of the seven pairs: FLT3 D1Y 0.61 vs. 0.78 Å, PIK3CA E11K 1.13 vs. 2.09 Å, KRAS G6C 1.04 vs. 1.18 Å, KRAS G3D 0.39 vs. 0.60 Å, KRAS G5V 0.84 vs. 1.17 Å, and PIK3CA H2L 0.97 vs. 1.37 Å. The single exception is TP53 R8H, where the neoantigen is marginally more flexible than the wild type (0.64 vs. 0.53 Å). This is the same pair for which the free-energy calculation failed to resolve (Section 3.2) and whose RMSD distributions are almost indistinguishable (**Figure 1**). Thus, the three results (Free energy, RMSD distribution, and RMSF) agree with one another.

### 3.8. Residue-level interaction network of the PIK3CA E545K pair: how a single anchor substitution reorganizes the entire C-terminal interface

The PIK3CA E545K pair (neoantigen STRDPLSEITK, wild-type STRDPLSEITE, both restricted by HLA-A*11:01) is the most distinctive in the panel across all readouts reported so far: it has the largest free-energy separation (ΔΔG = −14.29 ± 1.83 kcal/mol over the 300 ns window; **Table 3**), the largest asymmetry in terminal stability (the wild-type releases its C-terminus in 9/10 replicas and spends 56.9% of the analyzed time detached, against 4/10 and 19.7% for the neoantigen; **Table 7**, **Supplementary Figure S2**), the largest RMS-distribution separation (**Table 8**), and the largest fluctuation difference (**Figures 2 and 3**). We therefore reconstructed, frame by frame, the complete interaction network between each peptide residue and each of the 180 residues of the HLA groove, for all 20 trajectories of this pair (Section 2.11), to identify which specific interactions account for that behavior.

Because the wild-type is detached for more than half of its trajectory, any contact difference measured across all frames would partly restate the unbinding result rather than explain it. All values below are therefore taken from the bound conditioning window, that is, the subset of 50–500 ns frames in which the wild-type C-terminal region satisfies the bound criterion. The comparison thus asks whether the wild-type interface is weaker even while it is engaged; the same conclusion is obtained in all four conditioning windows of Section 2.11.2.

#### 3.8.1. The difference is limited to the last three peptide positions

The difference map in **Figure 4** makes the result immediately visible: over positions P1 to P8, the two peptides engage the groove almost identically, and the entire difference is a compact block at the last three positions facing the F pocket, the pocket at the C-terminal end of the groove that receives the PΩ side chain. **Table 9** presents the same information quantitatively, interaction type by interaction type and position by position.

**Figure 4.**
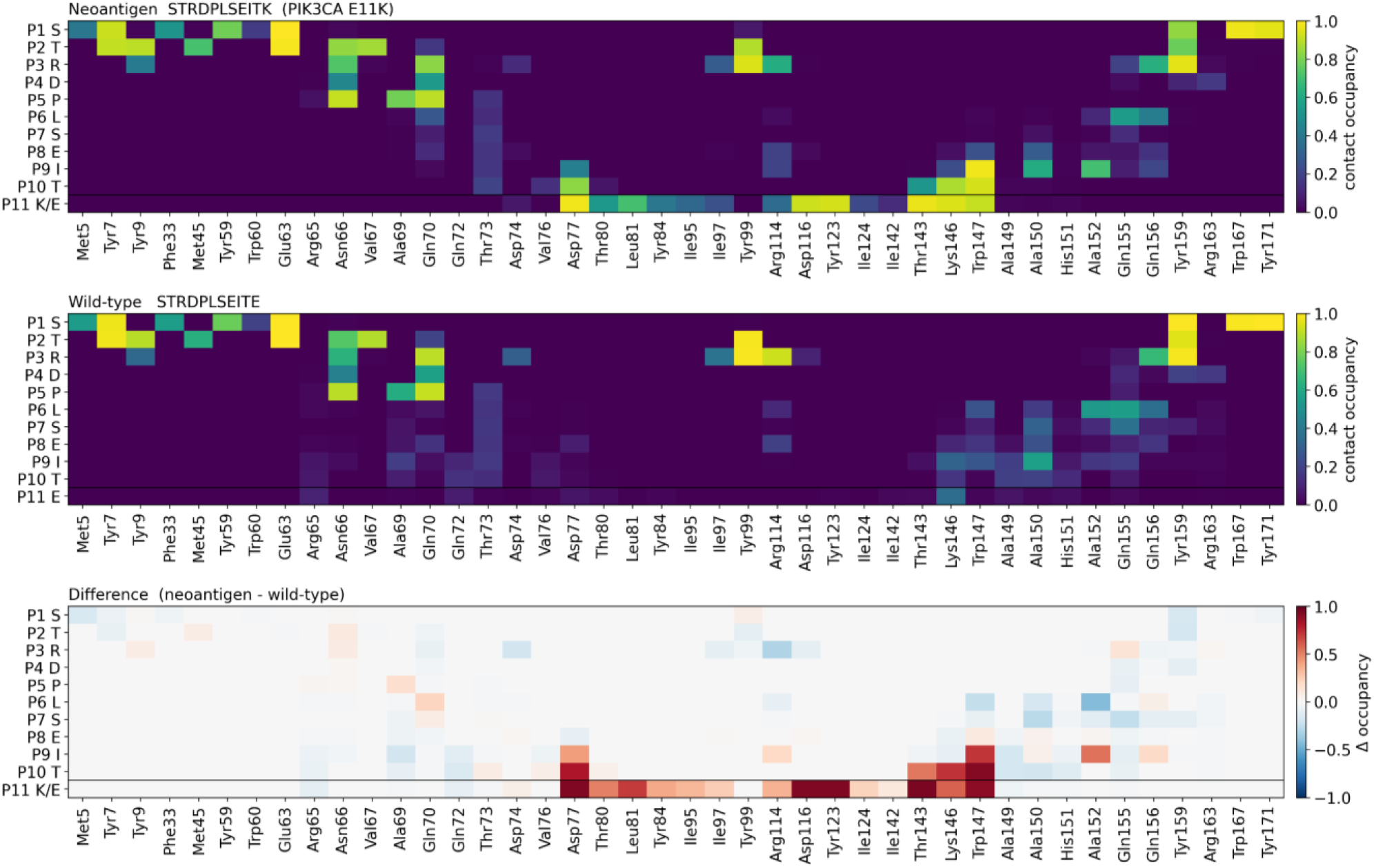
Contact occupancy for each peptide position against each HLA groove residue, computed only from bound frames (50–500 ns) and averaged across the ten replicas. A contact is scored when any heavy atom of the peptide residue lies within 4.5 Å of any heavy atom of the HLA residue, and occupancy is the fraction of frames in which that contact is present. Rows are peptide positions P1–P11, columns are the HLA groove residues that make any contact; horizontal black lines mark the mutated position P11. Top: neoantigen STRDPLSEITK. Middle: wild-type STRDPLSEITE. Bottom: the difference map (neoantigen − wild-type), with red indicating contacts made preferentially by the neoantigen and blue those made preferentially by the wild-type. The upper two thirds of the difference map is essentially blank: positions P1–P8 are indistinguishable between the two peptides. The entire difference is concentrated in a block at P9/P10/P11 against the F-pocket residues Asp77, Thr80, Leu81, Tyr84, Ile95, Asp116, Tyr123, Thr143, Lys146 and Trp147.

**Table 9.** For each interaction type, per peptide position and in bound frames, values are given as neoantigen / wild-type (mean over ten replicas). The two peptides are identical except at P11. Contacts are summed occupancies over all 180 groove residues (so they can exceed 1 and indicate how much of the groove that position engages); hydrogen bonds, salt bridges, hydrophobic contacts and cation–π interactions are mean counts per frame; vdW and “Screened elec.” interactions are given in kcal/mol. “Screened elec.” applies a distance-dependent dielectric with Debye damping and is a qualitative descriptor; it is not a real free energy value (Section 2.11.1).

| Pos. | Res.<br>neo / wt | Contacts | H-bonds | Salt<br>bridges | Hydrophobic | Cation-π | vdW | Screened<br>elec. |
| --- | --- | --- | --- | --- | --- | --- | --- | --- |
| P1 | S | 6.72 / 7.07 | 4.07 / 4.58 | 0.0 / 0.0 | 0.0 / 0.0 | 0.0 / 0.0 | -3.8 / -3.6 | -13.5 / -14.2 |
| P2 | T | 7.05 / 7.27 | 2.61 / 2.55 | 0.0 / 0.0 | 4.2 / 4.6 | 0.0 / 0.0 | -8.1 / -8.5 | -6.3 / -6.3 |
| P3 | R | 5.84 / 6.36 | 1.52 / 1.91 | 0.09 / 0.16 | 5.9 / 6.8 | 0.77 / 0.64 | -14.9 / -16.8 | -2.0 / -2.7 |
| P4 | D | 1.32 / 1.51 | 0.39 / 0.29 | 0.15 / 0.14 | 0.0 / 0.1 | 0.0 / 0.0 | -2.8 / -3.4 | -1.7 / -1.5 |
| P5 | P | 2.80 / 2.70 | 0.02 / 0.02 | 0.0 / 0.0 | 2.1 / 2.0 | 0.0 / 0.0 | -6.0 / -5.8 | -0.2 / -0.1 |
| P6 | L | 1.57 / 2.37 | 0.12 / 0.05 | 0.0 / 0.0 | 1.5 / 2.8 | 0.0 / 0.0 | -3.2 / -4.9 | 0.0 / 0.0 |
| P7 | S | 0.47 / 1.45 | 0.05 / 0.26 | 0.0 / 0.0 | 0.0 / 0.0 | 0.0 / 0.0 | -1.2 / -2.3 | -0.1 / -0.4 |
| P8 | E | 1.39 / 1.57 | 0.49 / 0.68 | 0.20 / 0.29 | 0.5 / 0.3 | 0.0 / 0.0 | -2.1 / -1.8 | -1.7 / -2.4 |
| P9 | I | 3.73 / 2.62 | 0.19 / 0.08 | 0.0 / 0.0 | 5.0 / 3.7 | 0.0 / 0.0 | -7.4 / -5.2 | 0.3 / -0.1 |
| P10 | T | 3.60 / 1.51 | 1.45 / 0.27 | 0.0 / 0.0 | 0.3 / 0.5 | 0.0 / 0.0 | -4.0 / -2.5 | -4.5 / -0.5 |
| P11 | K / E | 8.78 / 1.24 | 4.81 / 0.90 | 2.19 / 0.60 | 8.7 / 0.4 | 0.20 / 0.00 | -8.4 / -0.8 | -20.9 / -4.9 |
| **Total** |  | **43.28 / 35.67** | **15.72 / 11.58** | **2.63 / 1.19** | **28.3 / 21.2** | **0.96 / 0.64** | **-61.8 / -55.6** | **-50.6 / -33.2** |

Every interaction type moves in the same direction and each separates the two sets of replicas **(Table 10).** Yet in each case, the entire difference comes from P9–P11: over P1–P8, the wild-type is level with, or slightly ahead of the neoantigen on every measure (**Table 9**). The wild-type is therefore not making a weaker version of the same interface. It is making the same N-terminal interface and essentially no C-terminal one, with its P11 contributing 1.24 units of contact occupancy against 8.78 for the neoantigen’s P11.

**Table 10.** Interface totals with replica-level statistics (bound frames, mean ± SEM across ten replicas). p is the result of a two-sided Mann–Whitney U test comparing the ten neoantigen and the ten wild-type replica values; p ≈ 1.8 × 10⁻⁴ is the floor of the test at this sample size and indicates that the two groups of replicas do not overlap at all (Section 2.11.3).

| Interaction | Neoantigen | Wild-type | $\Delta$ (neo – wt) | $p$ |
| --- | --- | --- | --- | --- |
| Contacts (summed occupancy) | $43.28 \pm 0.60$ | $35.67 \pm 0.38$ | +7.61 | $1.8 \times 10^{-4}$ |
| Hydrogen bonds | $15.72 \pm 0.39$ | $11.58 \pm 0.25$ | +4.14 | $1.8 \times 10^{-4}$ |
| Salt bridges | $2.63 \pm 0.20$ | $1.19 \pm 0.14$ | +1.44 | $2.5 \times 10^{-4}$ |
| Hydrophobic contacts | $28.27 \pm 1.18$ | $21.21 \pm 0.84$ | +7.06 | 0.003 |
| Cation–π interactions | $0.96 \pm 0.07$ | $0.64 \pm 0.12$ | +0.33 | 0.031 |
| vdW energy (kcal/mol) | $-61.80 \pm 1.08$ | $-55.56 \pm 0.77$ | -6.24 | 0.002 |
| Screened electrostatics (kcal/mol) | $-50.64 \pm 1.37$ | $-33.21 \pm 1.25$ | -17.44 | $1.8 \times 10^{-4}$ |

Two chemistry-specific columns in **Table 9** merit separate mention because a contact count alone would obscure them. Salt bridges more than double at the interface (2.63 vs. 1.19 per frame), and 2.19 of the neoantigen’s 2.63 salt bridges belong to P11 alone. Cation–π interactions are numerically small but categorical: the neoantigen’s Lys11 contacts the Tyr123 ring in 94% of bound frames (**Table 11**) and satisfies the cation–π geometry in 20% of them (0.20 per frame; **Table 9**), whereas the wild-type’s Glu11 does neither and, being an anion, cannot form a cation–π interaction at all. The remaining cation–π signal in both peptides comes from Arg3 (0.77 vs. 0.64 per frame) and is shared.

**Table 11.** The 14 largest per-contact differences between the two peptides, ranked by Δ occupancy (bound frames).

| Peptide position | HLA residue | Occupancy neo | Occupancy wt | $\Delta$ |
| --- | --- | --- | --- | --- |
| P11 K/E | Thr143 | 0.971 | 0.027 | +0.944 |
| P11 K/E | Tyr123 | 0.935 | 0.009 | +0.926 |
| P11 K/E | Asp116 | 0.923 | 0.001 | +0.923 |
| P11 K/E | Asp77 | 0.979 | 0.057 | +0.922 |
| P10 T | Trp147 | 0.937 | 0.017 | +0.920 |
| P11 K/E | Trp147 | 0.913 | 0.017 | +0.896 |
| P10 T | Asp77 | 0.830 | 0.022 | +0.808 |
| P10 T | Lys146 | 0.878 | 0.155 | +0.723 |
| P9 I | Trp147 | 0.979 | 0.267 | +0.712 |
| P11 K/E | Leu81 | 0.703 | 0.005 | +0.698 |
| P11 K/E | Lys146 | 0.948 | 0.358 | +0.591 |
| P9 I | Ala152 | 0.703 | 0.159 | +0.544 |
| P10 T | Thr143 | 0.525 | 0.001 | +0.524 |
| P11 K/E | Thr80 | 0.534 | 0.033 | +0.501 |

Note also that positions P9 (Ile) and P10 (Thr) carry identical residues in the two peptides yet differ markedly: P10 loses 2.1 of its 3.6 units of contact occupancy and more than three quarters of its hydrogen bonds in the wild-type, and P9 loses about a third of its contacts and a quarter of its hydrophobic packing. Their weakening cannot be a local chemical effect of the substitution, because there is no local chemical change at those positions; it is a consequence of the P11 anchor failing.

Finally, the two quantities in **Tables 9 and 10** that are expressed in energy units can be compared directly, and they are not affected to the same degree. The vdW term differs between the two peptides by −6.24 ± 1.33 kcal/mol, whereas the screened electrostatic term differs by −17.44 ± 1.86 kcal/mol, a factor of 2.8 larger. In relative terms, the neoantigen’s electrostatic interaction with the groove is 53% stronger than the wild-type’s, against 11% for vdW. The most likely origin of that asymmetry is the inversion of formal charge at the mutated position. Lys11 carries a formal +1, and Glu11 a formal −1. The A*11:01 F pocket that receives the PΩ side chain is lined by two carboxylates, Asp77 and Asp116. The pocket that attracts the neoantigen’s side chain therefore repels the wild-type’s, so the interaction at this position does not merely weaken; it changes sign. The per-position decomposition in **Table 9** supports this directly: of the −17.44 kcal/mol total, −16.04 kcal/mol (92% of the whole electrostatic difference) is contributed by P11 alone, with P10 adding a further −4.06 kcal/mol and small positive contributions at P1, P3, and P8 offsetting part of the sum. The vdW term behaves differently: P11 alone accounts for −7.6 kcal/mol, more than the interface total of −6.2, because several other positions offset it. One caveat applies to this comparison. The screened electrostatic term describes the peptide–HLA interaction only and carries no desolvation penalty, whereas burying a charged side chain in a pocket incurs exactly such a penalty. It therefore locates where the interaction differs, not how much of the binding free energy that difference contributes; the Poisson–Boltzmann treatment discussed in Section 3.8.4 is what accounts for the compensation.

#### 3.8.2. A single salt bridge accounts for the largest edge differences

Ranking all 120 tested residue-pair edges by the magnitude of their contact-occupancy difference (**Table 11**) identifies which specific pairwise interactions are lost. Every one of the fourteen strongest differential edges involves P9, P10, or P11 paired with a residue in the F pocket, and the underlying chemistry is straightforward. The A*11:01 F pocket is built around Asp116, and HLA-A*11:01 is an allele whose C-terminal anchor preference is for a basic residue: crystallographic analysis of A*11:01 in complex with immunodominant epitopes confirms primary anchors at P2 (Ile/Val) and at the C-terminal Lys (**42**). The neoantigen’s Lys11 forms a salt bridge with Asp116 that is present in 88% of bound frames, at a mean charged-group distance of 3.26 Å. The wild-type’s Glu11 carries the same negative charge as Asp116; it essentially never forms that interaction (mean occupancy 0.0001, and exactly zero in nine of the ten replicas) and sits a mean of 19.2 Å away from it.

The consequence is greater than the loss of a single contact. Every class I peptide, regardless of sequence, engages its C-terminal carboxylate in the F pocket via the conserved Thr143/Lys146 pair; both peptides here possess that carboxylate, but only the neoantigen retains it (P11 carboxylate–Lys146 occupancy 0.94 vs. 0.11, and mean distance 3.2 Å vs. 13.4 Å). In other words, losing the side-chain anchor also costs the wild-type the sequence-independent backbone anchor that every class I peptide relies on, which is why the damage propagates back to P10 and P9.

#### 3.8.3. The wild-type makes more worse contacts

Rendering the same data as a network (**Figure 5**) reveals a feature that neither the contact map nor the totals convey. The wild-type forms more distinct residue-pair edges than the neoantigen (95 against 86) while carrying less total occupancy weight (34.0 against 42.5). Its C-terminal third is not immobilized in a single pocket; it samples many alternative positions, each held only briefly. The neoantigen’s P11K is the single largest hub in its network, whereas the wild-type’s P11E is the smallest node of its own. This is the network signature of the RMSD distribution and the elevated RMSF reported for this peptide in Section 3.6: an anchor that is present holds the peptide in one state, and an anchor that is absent lets it explore many.

**Figure 5.**
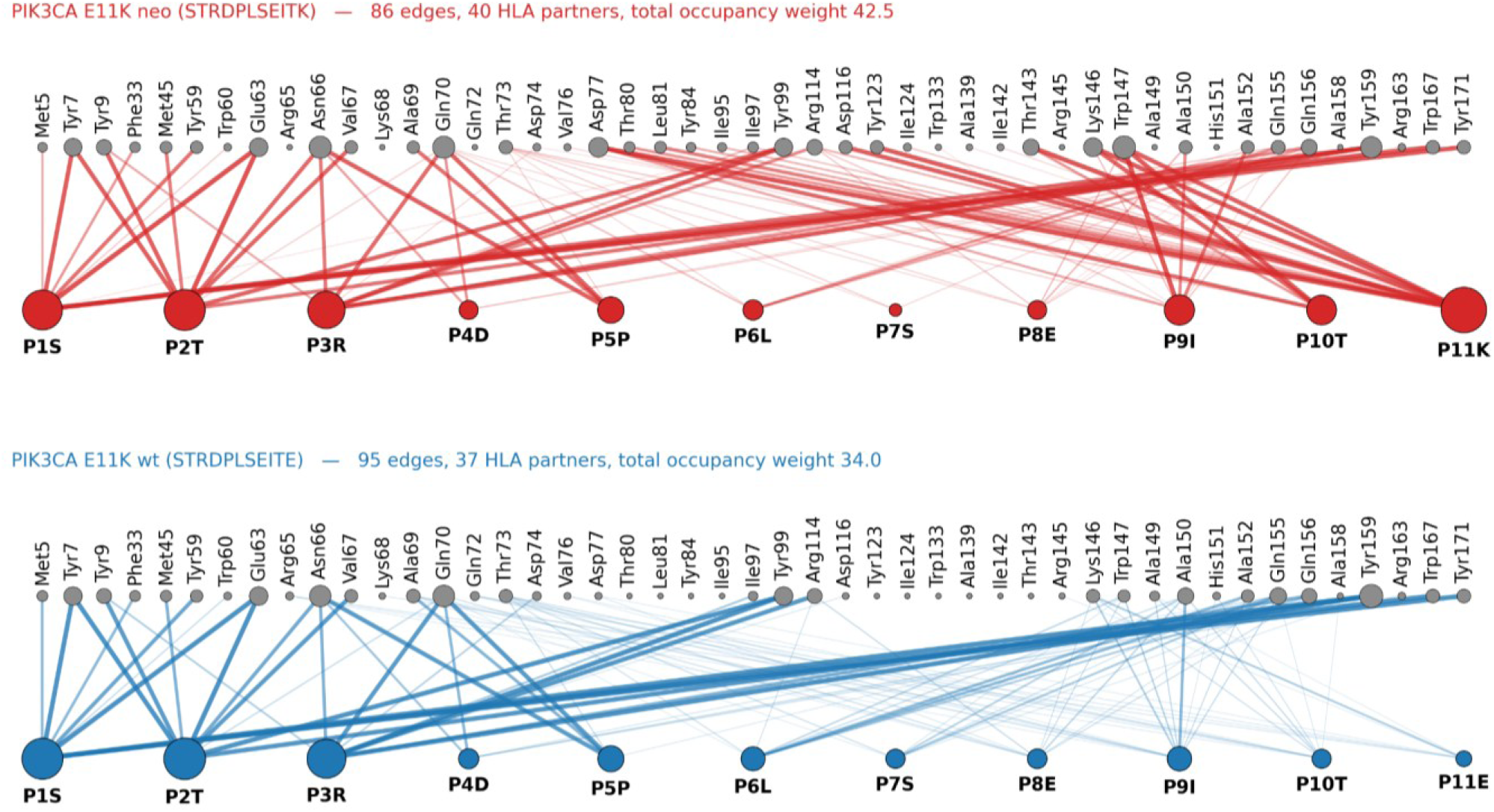
The peptide–HLA interface as a bipartite interaction network, computed over bound frames (50–500 ns). Lower nodes represent the eleven peptide positions, upper nodes represent the HLA groove residues, and an edge is drawn for each residue pair with contact occupancy ≥ 0.05. Edge width is proportional to contact occupancy and node size to weighted degree, i.e., the sum of occupancies, of all edges of that node. Top (red): neoantigen STRDPLSEITK, 86 edges to 40 HLA partners, total occupancy weight 42.5. Bottom (blue): wild-type STRDPLSEITE, 95 edges to 37 HLA partners, total occupancy weight 34. P11K is the largest hub of the neoantigen network, whereas P11E is the smallest node in its own network. Note that the wild-type has more edges but less total weight: having lost its anchor, it makes many weak and transient contacts instead of a few persistent ones.

#### 3.8.4. Implications for the MM-PBSA ranking of this pair

This pair has by far the largest free-energy separation in the panel, raising the legitimate concern that the value is inflated because the wild-type spends more than half of its trajectory detached, so the ensemble entering the MM-PBSA calculation is not that of a bound complex. The interaction energies indicate that the effect is real rather than an artifact of the detached frames: restricting the analysis to bound frames (the most conservative choice available) still leaves a screened electrostatic difference of −17.4 ± 1.9 kcal/mol and a vdW difference of −6.2 kcal/mol in favor of the neoantigen (**Table 10**). The screened electrostatic term is the one that should be compared with a solvated binding enthalpy, and its magnitude is of the right order for the MM-PBSA ΔΔH of −11.75 ± 2.05 kcal/mol computed over the same 50–500 ns window (against ΔΔG = −12.89 ± 2.19 kcal/mol; at the 300 ns working window used throughout Sections 3.2–3.4 the corresponding values are ΔΔH = −12.98 ± 1.66 and ΔΔG = −14.29 ± 1.83 kcal/mol). By contrast, the raw vacuum Coulomb difference for this pair is more than an order of magnitude larger, because no desolvation penalty is subtracted from it; this is precisely the term that the Poisson–Boltzmann treatment exists to correct.

One caveat should nevertheless be attached to the reported ΔΔG for this pair. The wild-type MM-PBSA ensemble genuinely is a mixture of bound and partially released states. The ΔΔG fairly describes f that ensemble, and it is the ensemble that a peptide of this sequence actually populates, but it is not the free-energy difference between two properly bound complexes and should not be read as one.

## 4. Discussion

Tumor neoantigens closely resemble autoantigens; however, the T cells that recognize them are not subject to central or peripheral tolerance and are expected to trigger a highly specific antitumor response. Although the use of neoantigens as personalized vaccines is promising, selecting neoantigens for effective clinical application in cancer immunotherapy remains a formidable challenge. Only a very small percentage of private neoepitopes encoded by tumor mutations bind to MHC molecules, a necessary condition for stimulating tumor-infectious T cell receptors (TCRs). Unlike personal neoantigens, public neoantigens are shared among tumor subtypes or patient populations (**14**), and several recent clinical studies demonstrate that their immediate availability as therapeutic vaccine candidates for broad-spectrum immunotherapy could be possible (**43–47**).

The ability of a neoantigen to elicit an efficient antitumor T-cell response is multifactorial. It depends on its natural processing by tumor cells, its ability to form stable pMHC complexes, and TCR recognition of the structure and dynamics of the pMHC-TCR complex. Unlike infectious-agent epitopes, neoepitopes are altered self-peptides derived from host-encoded proteins that must overcome immune tolerance to their corresponding wild-type (WT) self-peptides. Several groups (**48–52**) have suggested that, in addition to the neoantigen’s binding affinity and groove conformation, other physical properties, such as solvent-exposed surface area or the complex’s flexibility, may be important in determining neoantigen activity. Such differences, acting alone or in combination, could promote productive TCR binding and enable a neoepitope to overcome T cell tolerance to the WT peptide. Therefore, it is reasonable to explore whether molecular dynamics can provide additional information to existing neoantigen selection and prioritization algorithms.

We simulated seven public neoantigens with published experimental evidence of immunogenicity in preclinical and clinical studies, together with their non-immunogenic wild-type counterparts, each in complex with the HLA class I allele, and added three frameshift-derived neoantigens as unpaired immunogenic references for which immunogenicity had been reported. Most of the public neoantigen-MHC complexes studied showed greater neoantigen stability than the WT epitope. Most of the neoantigens relied on modifications of anchored residues. The TP53 R175H neoantigen exhibited the most pronounced trajectory over 300 ns.

The central result of the present study is that an ensemble binding free energy computed from HMR-accelerated trajectories orders the pairs in agreement with the experimental data. ΔΔG is negative in all seven pairs, and in six of them the separation exceeds its own uncertainty; the TP53 R175H pair is not resolved (ΔΔG = −0.94 ± 1.94 kcal/mol). This extends our earlier two-peptide proof of concept (**11**) to a larger, considerably more diverse panel (four HLA alleles, peptide lengths ranging from 9 to 11 residues, and substitutions distributed across the peptide) and, more importantly, replaces a qualitative assessment of trajectory stability with a quantitative criterion. The 4 fs HMR time step made this feasible: none of the 170 trajectories showed numerical instability, peptides remained accommodated in the groove for most of each trajectory in most complexes, and the doubled time step halved the wall-clock cost of what would otherwise have been ≈85 µs of conventional sampling.

Three independent readouts of the same trajectories agree with one another. Energetically, the neoantigen forms the more favorable complex (**Table 3**). Structurally, the neoantigen is the more constrained peptide: it has the lower mean Cα RMSF in six of seven pairs and the lower mean internal RMSD in four of seven pairs (**Table 8**, **Figures 1–3**). That three measures constructed from different atoms, reference frames and criteria converge on the same ordering is the strongest internal evidence we can offer that the ranking reflects a physical property of the complexes rather than an artifact of any single estimator.

Some studies on shared neoepitopes and tumor antigens (**48,53,54**) suggest that greater stiffness would favor T cell recognition by reducing the entropic cost of TCR binding. It has been proposed that characteristics such as the solvent exposure of the peptide-MHC interaction, the degree of solvent exposure of the neoantigen, and the hydrophobicity of the surface are especially important when the mutation includes amino acids in the center of the peptide that are part of the linking group with long or bulky side-chain anchors, as would be the case in our study for KRAS (G6C) and KRAS (G5V).

In our study, anchor and non-anchor substitutions behave differently, and most of our panel consists of non-anchor. It is worth stating explicitly, because it bears on how far these results generalize, that only two of the seven substitutions fall at a canonical class I anchor position. Class I ligands are held mainly by two side chains, one at P2 and one at the C-terminal residue PΩ(**55**); by that definition, PIK3CA E11K sits exactly at PΩ and PIK3CA H2L exactly at P2, whereas FLT3 D1Y (P1), KRAS G3D (P3), KRAS G5V (P5), KRAS G6C (P6 of a 10-mer) and TP53 R8H (P8 of a 9-mer) do not. These two locations correspond to two mechanisms by which a somatic mutation can create a neoepitope, a distinction that can be formalized as group I neoantigens, in which the mutation is buried and interacts mainly with the MHC binding pocket, versus group II, in which the mutated side chain is solvent-facing and can contact the T-cell receptor directly (**56**). The difference bears on what a binding calculation can detect. In the group I case, the substitution generates or improves an anchor residue, so the mutant peptide fits the groove better than its germline counterpart and is presented more efficiently at the tumor-cell surface: the novelty is one of presentation. In the group II case, the substitution occurs at a non-anchor position, typically a central one whose side chain points towards the T-cell-receptor contact sites (**13**). The anchor residues are then identical in the two peptides, so both would be expected to be presented comparably well, and the novelty is instead one of recognition: a neoantigen-specific receptor discriminates a single non-anchor substitution from a germline sequence against which the repertoire has largely been rendered tolerant by negative selection. Which of the two routes yields the more strongly immunogenic peptides remains debated (**13**). Our data show that the two are distinguishable in simulation: the two anchor substitutions produce the two largest fluctuation differences in the panel, and in each case the destabilization is localized to the terminus that the mutated anchor holds: C-terminal for E11K, N-terminal for H2L (**Figure 2**, **Table 7**). The five non-anchor substitutions produce smaller and more diffuse fluctuation differences, and the three narrowest free-energy separations in the panel are all non-anchor (KRAS G5V, −2.60 ± 1.69; TP53 R8H, −0.94 ± 1.94; KRAS G6C, −4.22 ± 2.12 kcal/mol). The tendency is not a rule, however: KRAS G3D is a non-anchor substitution at P3 and nonetheless gives the second-largest separation in the panel (−9.05 ± 1.29 kcal/mol, ≈7 SEM), larger than either anchor pair except E11K. With only two anchor substitutions among seven pairs, anchor position is at most a suggestive predictor of how sharply the calculation will discriminate.

The TP53 R8H pair is an interesting case, and it fails in an informative way. The two **HMTEVVR[R/H]C** complexes are the most favorably bound in the entire panel in absolute terms (ΔG ≈ −75 to −76 kcal/mol for both peptides), they are indistinguishable in their RMSD distributions (1.10 ± 0.36 vs. 1.04 ± 0.23 Å, p = 0.571), their central-residue fluctuation profiles are superimposed within error (**Figure 3**), and both are essentially stable in the groove (**Table 7**). This is the one pair in which the neoantigen is marginally the more flexible peptide. In other words, the calculation does not fail by returning noise; it returns a consistent answer, namely that these two peptides bind HLA-A*02:01 almost equally well and behave almost identically once bound. The substitution is at P8 of a 9-mer, not at an anchor, and R→H is a conservative replacement of one basic side chain by a smaller, weakly basic one at a position that points towards the T-cell receptor rather than into the groove. The most parsimonious interpretation is that the immunogenicity of this neoantigen is not encoded in its MHC binding at all, but at the TCR interface, which our simulations do not model. A binding-energy criterion is therefore necessarily a filter on one of several requirements for immunogenicity, and TP53 R8H is a clean illustration of a peptide that passes the T-cell discrimination test without passing the binding one.

Concerning simulation length and replica ensembles, two method-level findings have direct practical value for anyone deploying this approach. First, the pairwise rankings are established early: every ΔΔG is already negative at the 100 ns window and none changes sign out to 500 ns (**Table 5**). What is not established at 100 ns is the magnitude of the per-complex free energy, and it is convergence of the ensemble mean, not of the ranking, that sets the working simulation time. All 17 complexes reach a common plateau at 300 ns (agreement within 2 SEM across the 300, 400 and 500 ns windows), while extending each replica from 300 to 500 ns costs 67% more simulation time for no change in ranking and no change in ensemble mean beyond ≈1.6 kcal/mol. Second, and more importantly, single trajectories are not usable as estimators: per-replica values span up to ≈25 kcal/mol within one complex, and the standard deviation across replicas reaches 5.8 kcal/mol (**Table 6, Supplementary Figure S1**). The dispersion is itself informative, since the two frameshift peptides with the highest spread (TGFBRII, 5.6; APC, 5.2) are also the two complexes with the most frequent terminal unbinding, consistent with a flexible peptide broadening the conformational ensemble and thereby inflating the between-replica variance. Ensemble averaging over ten replicas is therefore not a refinement of this protocol; it is a precondition for it. Combining HMR with replica averaging is what makes the combination affordable: the factor of two recovered from the time step is spent on independent trajectories rather than on longer ones.

The PIK3CA E11K pair let us ask not only whether the calculation ranks correctly but why. The complete interaction network (Section 3.8) shows that the entire difference between **STRDPLSEITK** and **STRDPLSEITE** is confined to the last three peptide positions and to the F pocket. HLA-A*11:01 prefers a basic residue at PΩ (**42**), so the wild-type Glu11 is not merely a poorer anchor but an electrostatically repelled one: it carries the same charge as the aspartate that lines the pocket. The consequence extends beyond the mutated position. Losing the side-chain anchor also costs the wild-type the conserved, sequence-independent C-terminal carboxylate anchor to Lys146 (occupancy 0.94 vs. 0.11), and the damage propagates back to P10 and P9, positions whose residues are chemically identical in the two peptides and which nonetheless lose most of their hydrogen bonds and a third of their contacts. This is a concrete instance of the “delocalized” effect we proposed on much sparser evidence in our earlier work, in which a mutated residue was seen to modulate the interactions of distant peptide residues with the HLA molecule (**11**); here it is resolved edge by edge, with replica-level statistics, and its structural cause is identified. The network view **(Figure 5)** adds a detail that no contact count conveys: the wild-type forms more distinct residue-pair edges than the neoantigen (95 vs. 86) but carries less total occupancy weight (34.0 vs. 42.5). Having lost its anchor, it does not sit still and bind weakly; it explores many alternative poses, each held only briefly. That is the interaction-network signature of the RMSD distribution and the elevated RMSF measured for the same peptide in Section 3.7, and it illustrates why occupancy, rather than the mere existence of a contact in a static model, is the quantity that discriminates.

The three KRAS pairs are instructive cases because they are the panel’s clearest non-anchor cases, and because glycine is the residue whose replacement has the largest expected effect on backbone flexibility. In KRAS G6C (P6 of the 10-mer VVVGACGVGK) and KRAS G5V (P5 of the 9-mer VVGAVGVGK), a glycine that carries no side chain and confers maximal backbone freedom is replaced by a residue that carries one; in both, the neoantigen is measurably the less flexible peptide (mean Cα RMSF 1.04 vs. 1.18 Å and 0.84 vs. 1.17 Å respectively), which is what that chemistry predicts, and both ΔΔG values are concordant though modest (−4.22 ± 2.12 and −2.60 ± 1.69 kcal/mol). Central positions in an HLA-A*11:01 ligand are not necessarily purely TCR-facing, since secondary anchor positions in the middle of the peptide have been described for this allele (**42**), so a substitution there can plausibly act on both interfaces at once. Independent structural work on KRAS G12 variants (in the canonical notation) presented by HLA-A*11:01 reaches a compatible conclusion by different means, showing that single substitutions at position 12 remodel the peptide–MHC architecture and that peptide conformation and length, rather than binding alone, shape immune visibility (**41**). Our KRAS G12D result (presented by HLA-C*08:02 rather than HLA-A*11:01) is the second-largest separation in the panel (−9.05 ± 1.29 kcal/mol) and is accompanied by the most reproducible free energy of any complex (replica SD 1.9 kcal/mol) and the lowest peptide flexibility in the whole panel (mean Cα RMSF 0.39 Å, against 0.60 Å for its wild-type).

Several constraints should temper the interpretation. (i) MM-PBSA is an endpoint method: it is not expected to reproduce absolute binding free energies, and the values reported here should be read as a ranking scale, not as measurements. The quasi-harmonic entropy term compounds this, since it assumes a single harmonic basin and will be least reliable. (ii) The starting structures are AlphaFold3 predictions rather than experimental coordinates; although the ten replicas of each complex diverge substantially from one another, they all begin from the same model, so a systematic error in the predicted pose would not be detected by our replica statistics. (iii No T-cell receptor is present in any of the simulations, so nothing here addresses the recognition step directly. (iv) The experimental ground truth is binary and drawn from heterogeneous assays across different laboratories, which sets a ceiling on how finely any computational criterion can be validated against it.

In conclusion, these findings support HMR-accelerated binding free-energy MD as a practical refinement step for neoantigen prioritization and its application to assessing public neoantigens for cancer vaccine design. By combining a reduced computational cost with replica-based ensemble averaging, this approach reliably reproduces experimental immunogenicity rankings while also resolving the structural basis of peptide–HLA discrimination at atomic resolution. Taken together, these results position HMR-accelerated MD as a scalable computational filter that can complement existing in silico prediction tools, narrowing candidate lists before experimental validation and supporting the rational selection of public neoantigens for broad-spectrum cancer vaccines.

## Data and code availability

All codes and datasets supporting the findings of this study are openly available on Zenodo:

- DOI Code: https://doi.org/10.5281/zenodo.22548326
- DOI dataset: https://doi.org/10.5281/zenodo.22226388

## Acknowledges

We gratefully acknowledge the Ministry of Science, Technology, and Innovation of Colombia (MinCiencias) for funding this project. We also thank the Universidad Nacional de Colombia for its support of this research and for its support of Dr. Carlos Alberto Parra López during his sabbatical year.

## Authorship

All authors meet the ICMJE criteria for authorship. Each author made substantial contributions to the conception or design of the work, or the acquisition, analysis, or interpretation of data; drafted the work or revised it critically for important intellectual content; approved the final version to be published; and agrees to be accountable for all aspects of the work in ensuring that questions related to the accuracy or integrity of any part of the work are appropriately investigated and resolved. DA, CP contributed to the conception and design of the study; DA performed data acquisition, analysis, and interpretation; DA, JDM, JBFS, CP drafted the manuscript; DA, JDM, JBFS, CP critically revised it for important intellectual content. All authors approved the final version of the manuscript and agree to be accountable for all aspects of the work.

## References

1. Weber JS, Carlino MS, Khattak A, Meniawy T, Ansstas G, Taylor MH, et al. Individualised neoantigen therapy mRNA-4157 (V940) plus pembrolizumab versus pembrolizumab monotherapy in resected melanoma (KEYNOTE-942): a randomised, phase 2b study. Lancet. 2024 Feb 17;403(10427):632–44. doi:10.1016/S0140-6736(23)02268-7 PubMed PMID: 38246194.

2. Rojas LA, Sethna Z, Soares KC, Olcese C, Pang N, Patterson E, et al. Personalized RNA neoantigen vaccines stimulate T cells in pancreatic cancer. Nature. 2023 Jun;618(7963):144–50. doi:10.1038/s41586-023-06063-y PubMed PMID: 37165196.

3. Schumacher TN, Schreiber RD. Neoantigens in cancer immunotherapy. Science (1979). 2015 Apr 3;348(6230):69–74. doi:10.1126/science.aaa4971

4. Jiang T, Shi T, Zhang H, Hu J, Song Y, Wei J, et al. Tumor neoantigens: from basic research to clinical applications. J Hematol Oncol. 2019 Dec 6;12(1):93. doi:10.1186/s13045-019-0787-5

5. Reynisson B, Alvarez B, Paul S, Peters B, Nielsen M. NetMHCpan-4.1 and NetMHCIIpan-4.0: improved predictions of MHC antigen presentation by concurrent motif deconvolution and integration of MS MHC eluted ligand data. Nucleic Acids Res. 2020 Jul 2;48(W1):W449–54. doi:10.1093/nar/gkaa379

6. Wells DK, van Buuren MM, Dang KK, Hubbard-Lucey VM, Sheehan KCF, Campbell KM, et al. Key Parameters of Tumor Epitope Immunogenicity Revealed Through a Consortium Approach Improve Neoantigen Prediction. Cell. 2020 Oct;183(3):818–834.e13. doi:10.1016/j.cell.2020.09.015

7. Harndahl M, Rasmussen M, Roder G, Dalgaard Pedersen I, Sørensen M, Nielsen M, et al. Peptide-MHC class I stability is a better predictor than peptide affinity of CTL immunogenicity. Eur J Immunol. 2012 Jun 8;42(6):1405–16. doi:10.1002/eji.201141774

8. Rasmussen M, Fenoy E, Harndahl M, Kristensen AB, Nielsen IK, Nielsen M, et al. Pan-Specific Prediction of Peptide–MHC Class I Complex Stability, a Correlate of T Cell Immunogenicity. The Journal of Immunology. 2016 Aug 15;197(4):1517–24. doi:10.4049/jimmunol.1600582

9. Riley TP, Keller GLJ, Smith AR, Davancaze LM, Arbuiso AG, Devlin JR, et al. Structure Based Prediction of Neoantigen Immunogenicity. Front Immunol. 2019 Aug 28;10. doi:10.3389/fimmu.2019.02047

10. Genheden S, Ryde U. The MM/PBSA and MM/GBSA methods to estimate ligand-binding affinities. Expert Opin Drug Discov. 2015 May 4;10(5):449–61. doi:10.1517/17460441.2015.1032936

11. Amaya-Ramirez D, Martinez-Enriquez LC, Parra-López C. Usefulness of Docking and Molecular Dynamics in Selecting Tumor Neoantigens to Design Personalized Cancer Vaccines: A Proof of Concept. Vaccines (Basel). 2023 Jun 29;11(7):1174. doi:10.3390/vaccines11071174

12. Hopkins CW, Le Grand S, Walker RC, Roitberg AE. Long-Time-Step Molecular Dynamics through Hydrogen Mass Repartitioning. J Chem Theory Comput. 2015 Apr 14;11(4):1864–74. doi:10.1021/ct5010406

13. Goloudina A, Le Chevalier F, Authié P, Charneau P, Majlessi L. Shared neoantigens for cancer immunotherapy. Molecular Therapy Oncology. 2025 Jun;33(2):200978. doi:10.1016/j.omton.2025.200978

14. Gurung HR, Heidersbach AJ, Darwish M, Chan PPF, Li J, Beresini M, et al. Systematic discovery of neoepitope–HLA pairs for neoantigens shared among patients and tumor types. Nat Biotechnol. 2024 Jul 19;42(7):1107–17. doi:10.1038/s41587-023-01945-y

15. Abramson J, Adler J, Dunger J, Evans R, Green T, Pritzel A, et al. Accurate structure prediction of biomolecular interactions with AlphaFold 3. Nature. 2024 Jun 13;630(8016):493–500. doi:10.1038/s41586-024-07487-w

16. Strønen E, Toebes M, Kelderman S, van Buuren MM, Yang W, van Rooij N, et al. Targeting of cancer neoantigens with donor-derived T cell receptor repertoires. Science (1979). 2016 Jun 10;352(6291):1337–41. doi:10.1126/science.aaf2288

17. Lo W, Parkhurst M, Robbins PF, Tran E, Lu YC, Jia L, et al. Immunologic Recognition of a Shared p53 Mutated Neoantigen in a Patient with Metastatic Colorectal Cancer. Cancer Immunol Res. 2019 Apr 1;7(4):534–43. doi:10.1158/2326-6066.CIR-18-0686

18. Malekzadeh P, Pasetto A, Robbins PF, Parkhurst MR, Paria BC, Jia L, et al. Neoantigen screening identifies broad TP53 mutant immunogenicity in patients with epithelial cancers. Journal of Clinical Investigation. 2019 Feb 4;129(3):1109–14. doi:10.1172/JCI123791

19. Choi J, Goulding SP, Conn BP, McGann CD, Dietze JL, Kohler J, et al. Systematic discovery and validation of T cell targets directed against oncogenic KRAS mutations. Cell Reports Methods. 2021 Sep;1(5):100084. doi:10.1016/j.crmeth.2021.100084

20. Tran E, Robbins PF, Lu YC, Prickett TD, Gartner JJ, Jia L, et al. T-Cell Transfer Therapy Targeting Mutant KRAS in Cancer. N Engl J Med. 2016 Dec 8;375(23):2255–62. doi:10.1056/NEJMoa1609279 PubMed PMID: 27959684.

21. Leidner R, Sanjuan Silva N, Huang H, Sprott D, Zheng C, Shih YP, et al. Neoantigen T-Cell Receptor Gene Therapy in Pancreatic Cancer. New England Journal of Medicine. 2022 Jun 2;386(22):2112–9. doi:10.1056/NEJMoa2119662

22. Wang X, Wang W, Zou S, Xu Z, Cao D, Zhang S, et al. Combination therapy of KRAS G12V mRNA vaccine and pembrolizumab: clinical benefit in patients with advanced solid tumors. Cell Res. 2024 Jun 24;34(9):661–4. doi:10.1038/s41422-024-00990-9

23. Chandran SS, Ma J, Klatt MG, Dündar F, Bandlamudi C, Razavi P, et al. Immunogenicity and therapeutic targeting of a public neoantigen derived from mutated PIK3CA. Nat Med. 2022 May 28;28(5):946–57. doi:10.1038/s41591-022-01786-3

24. Shen M, Chen S, Han X, Hao Y, Wang J, Li L, et al. Identification of an HLA-A*11:01-restricted neoepitope of mutant PIK3CA and its specific T cell receptors for cancer immunotherapy targeting hotspot driver mutations. Cancer Immunology, Immunotherapy. 2024 Jun 4;73(8):150. doi:10.1007/s00262-024-03729-y

25. Koster J, Plasterk RHA. A library of Neo Open Reading Frame peptides (NOPs) as a sustainable resource of common neoantigens in up to 50% of cancer patients. Sci Rep. 2019 Apr 29;9(1):6577. doi:10.1038/s41598-019-42729-2

26. van der Lee DI, Reijmers RM, Honders MW, Hagedoorn RS, de Jong RCM, Kester MGD, et al. Mutated nucleophosmin 1 as immunotherapy target in acute myeloid leukemia. Journal of Clinical Investigation. 2019 Jan 14;129(2):774–85. doi:10.1172/JCI97482

27. Greiner J, Goetz M, Schuler PJ, Bulach C, Hofmann S, Schrezenmeier H, et al. Enhanced stimulation of antigen-specific immune responses against nucleophosmin 1 mutated acute myeloid leukaemia by an anti-programmed death 1 antibody. Br J Haematol. 2022 Sep;198(5):866–74. doi:10.1111/bjh.18326 PubMed PMID: 35799423.

28. Sæterdal I, Gjertsen M, Straten P, Eriksen J, Gaudernack G. A TGFβRII frameshift-mutation-derived CTL epitope recognised by HLA-A2-restricted CD8 + T cells. Cancer Immunology, Immunotherapy. 2001 Nov 1;50(9):469–76. doi:10.1007/s002620100222

29. Inderberg EM, Wälchli S, Myhre MR, Trachsel S, Almåsbak H, Kvalheim G, et al. T cell therapy targeting a public neoantigen in microsatellite instable colon cancer reduces *in vivo* tumor growth. Oncoimmunology. 2017 Apr 3;6(4):e1302631. doi:10.1080/2162402X.2017.1302631

30. Humphrey W, Dalke A, Schulten K. VMD: Visual molecular dynamics. J Mol Graph. 1996 Feb;14(1):33–8. doi:10.1016/0263-7855(96)00018-5

31. Huang J, Rauscher S, Nawrocki G, Ran T, Feig M, de Groot BL, et al. CHARMM36m: an improved force field for folded and intrinsically disordered proteins. Nat Methods. 2017 Jan 7;14(1):71–3. doi:10.1038/nmeth.4067

32. Jorgensen WL, Chandrasekhar J, Madura JD, Impey RW, Klein ML. Comparison of simple potential functions for simulating liquid water. J Chem Phys. 1983 Jul 15;79(2):926–35. doi:10.1063/1.445869

33. MacKerell AD, Bashford D, Bellott M, Dunbrack RL, Evanseck JD, Field MJ, et al. All-Atom Empirical Potential for Molecular Modeling and Dynamics Studies of Proteins. J Phys Chem B. 1998 Apr 1;102(18):3586–616. doi:10.1021/jp973084f

34. Phillips JC, Hardy DJ, Maia JDC, Stone JE, Ribeiro J V., Bernardi RC, et al. Scalable molecular dynamics on CPU and GPU architectures with NAMD. J Chem Phys. 2020 Jul 28;153(4). doi:10.1063/5.0014475

35. Essmann U, Perera L, Berkowitz ML, Darden T, Lee H, Pedersen LG. A smooth particle mesh Ewald method. J Chem Phys. 1995 Nov 15;103(19):8577–93. doi:10.1063/1.470117

36. Feller SE, Zhang Y, Pastor RW, Brooks BR. Constant pressure molecular dynamics simulation: The Langevin piston method. J Chem Phys. 1995 Sep 15;103(11):4613–21. doi:10.1063/1.470648

37. Case DA, Aktulga HM, Belfon K, Cerutti DS, Cisneros GA, Cruzeiro VWD, et al. AmberTools. J Chem Inf Model. 2023 Oct 23;63(20):6183–91. doi:10.1021/acs.jcim.3c01153

38. Roe DR, Cheatham TE. PTRAJ and CPPTRAJ: Software for Processing and Analysis of Molecular Dynamics Trajectory Data. J Chem Theory Comput. 2013 Jul 9;9(7):3084–95. doi:10.1021/ct400341p

39. Miller BR, McGee TD, Swails JM, Homeyer N, Gohlke H, Roitberg AE. MMPBSA.py: An Efficient Program for End-State Free Energy Calculations. J Chem Theory Comput. 2012 Sep 11;8(9):3314–21. doi:10.1021/ct300418h

40. Andricioaei I, Karplus M. On the calculation of entropy from covariance matrices of the atomic fluctuations. J Chem Phys. 2001 Oct 8;115(14):6289–92. doi:10.1063/1.1401821

41. Zhu J, Chen Z, Xu X, Wang Y, Liu P, Wen M, et al. Structure guided analysis of KRAS G12 mutants in HLA-A*11:01 reveals a length encoded immunogenic advantage in G12D. Commun Biol. 2025 Dec 3;9(1):26. doi:10.1038/s42003-025-09285-0

42. Li L, Bouvier M. Structures of HLA-A*1101 Complexed with Immunodominant Nonamer and Decamer HIV-1 Epitopes Clearly Reveal the Presence of a Middle, Secondary Anchor Residue. The Journal of Immunology. 2004 May 15;172(10):6175–84. doi:10.4049/jimmunol.172.10.6175

43. Haldar SD, Huff AL, Wang H Henry, Zhu Z, Berg M, Lu J, et al. First-in-human testing of a mutant KRAS vaccine for pancreatic cancer interception in high-risk cohorts. Cancer Discov. 2026 Jul 16. doi:10.1158/2159-8290.CD-25-2245

44. Pant S, Wainberg ZA, Weekes CD, Furqan M, Kasi PM, Devoe CE, et al. ar-specific amphiphile vaccine in pancreatic and colorectal cancer: the phase 1 AMPLIFY-201 trial. Nat Med. 2024 Feb;30(2):531–42. doi:10.1038/s41591-023-02760-3 PubMed PMID: 38195752.

45. Rappaport AR, Kyi C, Lane M, Hart MG, Johnson ML, Henick BS, et al. A shared neoantigen vaccine combined with immune checkpoint blockade for advanced metastatic solid tumors: phase 1 trial interim results. Nat Med. 2024 Apr;30(4):1013–22. doi:10.1038/s41591-024-02851-9 PubMed PMID: 38538867.

46. Sei S, Ahadova A, Keskin DB, Bohaumilitzky L, Gebert J, von Knebel Doeberitz M, et al. Lynch syndrome cancer vaccines: A roadmap for the development of precision immunoprevention strategies. Front Oncol. 2023;13:1147590. doi:10.3389/fonc.2023.1147590 PubMed PMID: 37035178.

47. D’Alise AM, Willis J, Duzagac F, Hall MJ, Cruz-Correa M, Idos GE, et al. Nous-209 neoantigen vaccine for cancer prevention in Lynch syndrome carriers: a phase 1b/2 trial. Nat Med. 2026 Mar;32(3):1002–11. doi:10.1038/s41591-025-04182-9 PubMed PMID: 41545594.

48. Duan F, Duitama J, Al Seesi S, Ayres CM, Corcelli SA, Pawashe AP, et al. Genomic and bioinformatic profiling of mutational neoepitopes reveals new rules to predict anticancer immunogenicity. J Exp Med. 2014 Oct 20;211(11):2231–48. doi:10.1084/jem.20141308 PubMed PMID: 25245761.

49. Ebrahimi-Nik H, Moussa M, Englander RP, Singhaviranon S, Michaux J, Pak H, et al. Reversion analysis reveals the in vivo immunogenicity of a poorly MHC I-binding cancer neoepitope. Nat Commun. 2021 Nov 5;12(1):6423. doi:10.1038/s41467-021-26646-5 PubMed PMID: 34741035.

50. Brennick CA, George MM, Moussa MM, Hagymasi AT, Seesi S Al, Shcheglova T V, et al. An unbiased approach to defining bona fide cancer neoepitopes that elicit immune-mediated cancer rejection. J Clin Invest. 2021 Feb 1;131(3). doi:10.1172/JCI142823 PubMed PMID: 33320837.

51. Antunes DA, Rigo MM, Freitas M V., Mendes MFA, Sinigaglia M, Lizée G, et al. Interpreting T-Cell Cross-reactivity through Structure: Implications for TCR-Based Cancer Immunotherapy. Front Immunol. 2017 Oct 4;8. doi:10.3389/fimmu.2017.01210

52. Devlin JR, Alonso JA, Ayres CM, Keller GLJ, Bobisse S, Vander Kooi CW, et al. Structural dissimilarity from self drives neoepitope escape from immune tolerance. Nat Chem Biol. 2020 Nov 17;16(11):1269–76. doi:10.1038/s41589-020-0610-1

53. Insaidoo FK, Borbulevych OY, Hossain M, Santhanagopolan SM, Baxter TK, Baker BM. Loss of T Cell Antigen Recognition Arising from Changes in Peptide and Major Histocompatibility Complex Protein Flexibility. Journal of Biological Chemistry. 2011 Nov;286(46):40163–73. doi:10.1074/jbc.M111.283564

54. Borbulevych OY, Insaidoo FK, Baxter TK, Powell DJ, Johnson LA, Restifo NP, et al. Structures of MART-126/27–35 Peptide/HLA-A2 Complexes Reveal a Remarkable Disconnect between Antigen Structural Homology and T Cell Recognition. J Mol Biol. 2007 Oct;372(5):1123–36. doi:10.1016/j.jmb.2007.07.025

55. Kubo RT, Sette A, Grey HM, Appella E, Sakaguchi K, Zhu NZ, et al. Definition of specific peptide motifs for four major HLA-A alleles. J Immunol. 1994 Apr 15;152(8):3913–24. PubMed PMID: 8144960.

56. Finnigan JP, Newman JH, Patskovsky Y, Patskovska L, Ishizuka AS, Lynn GM, et al. Structural basis for self-discrimination by neoantigen-specific TCRs. Nat Commun. 2024 Mar 8;15(1):2140. doi:10.1038/s41467-024-46367-9

