## Supplementary material for "Accelerating Molecular Dynamics with Hydrogen Mass Repartitioning to Prioritize Immunogenic Neoantigens for Cancer Vaccines through Peptide–HLA Binding Free-Energy Calculations"

**Supplementary materials**

**Figure S1.** Convergence and replica-to-replica dispersion of the binding free energy, one pair of panels per complex. For each complex, the left panel shows the ensemble binding free energy ΔGbind as a function of the length of the analysis window (100, 200, 300, 400 and 500 ns), plotted as the mean over the ten replicas with error bars and a shaded band giving ± 1 SEM across replicas. The right panel shows the same five windows resolved into the ten individual replicas (thin colored lines, one color per replica) together with the ensemble mean (bold black line with square markers). All axes are in kcal/mol; more negative values indicate more favorable binding. Complexes are grouped by gene and alteration, with the neoantigen shown first and its wild-type counterpart, where one exists, immediately below.

a) **FLT3 (D1Y)**

Neoantigen:

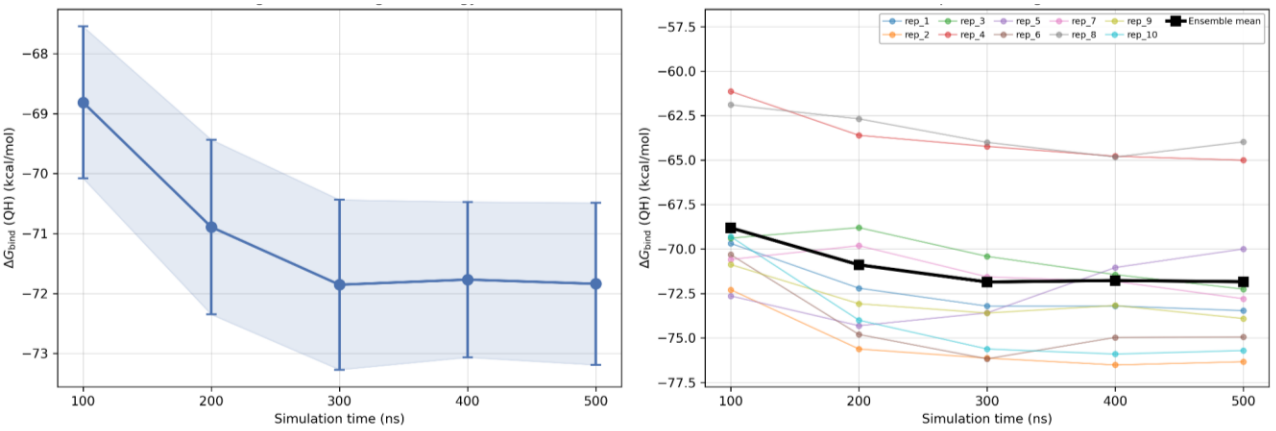


Wild-type:

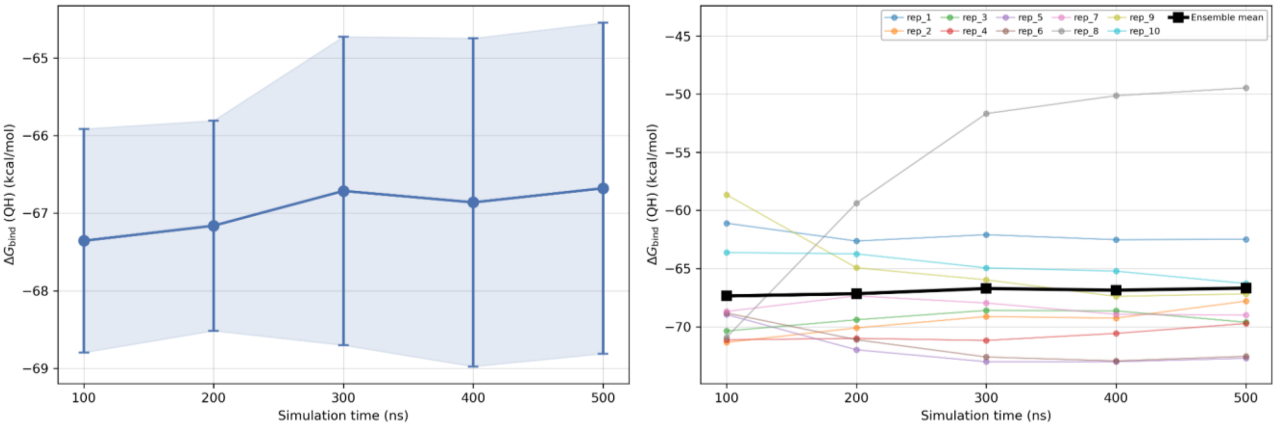

### b) APC (fs)

Neoantigen:

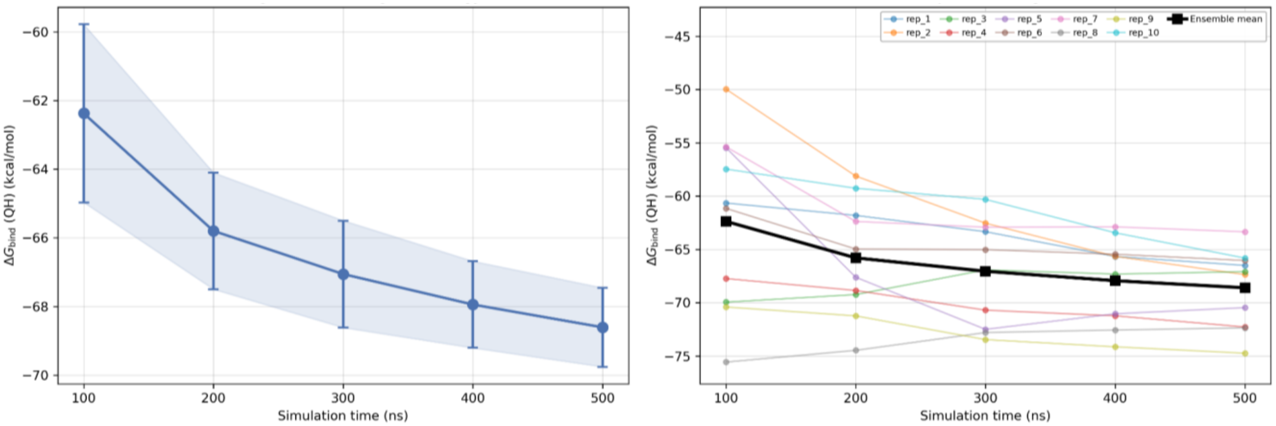


### c) PIK3CA (E11K)

Neoantigen:

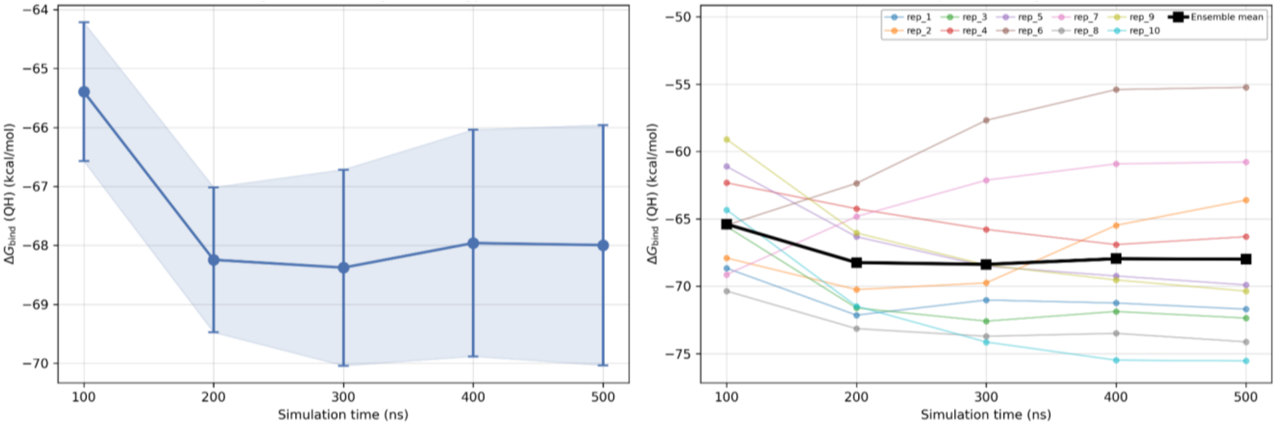


Wild-type:

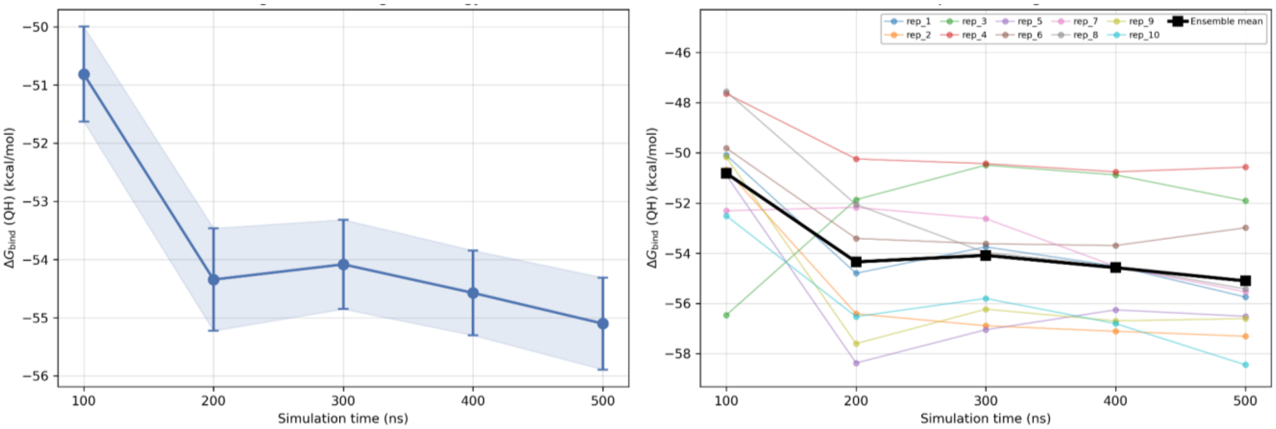

**d) NPM1 (fs)**

Neoantigen:

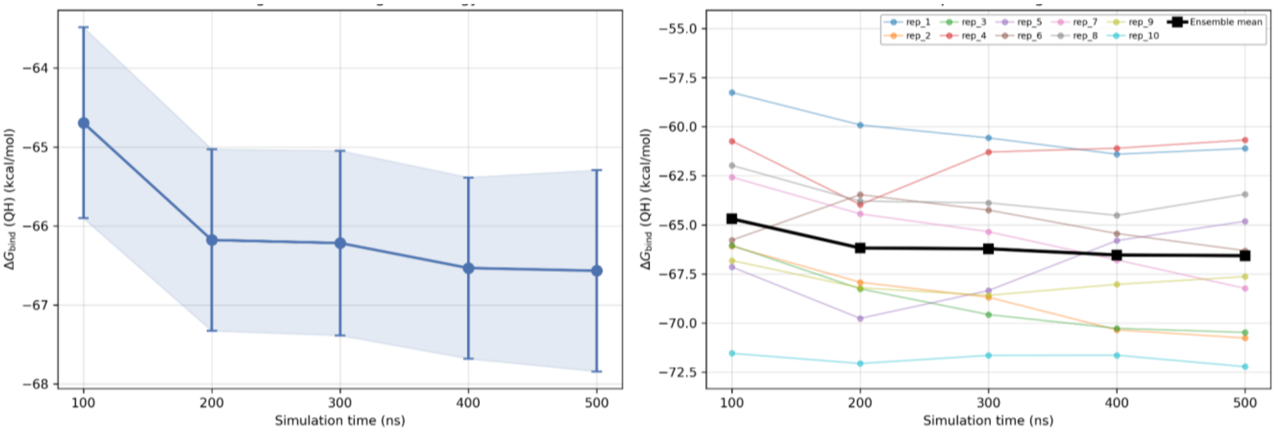


**e) KRAS (G6C)**

Neoantigen:

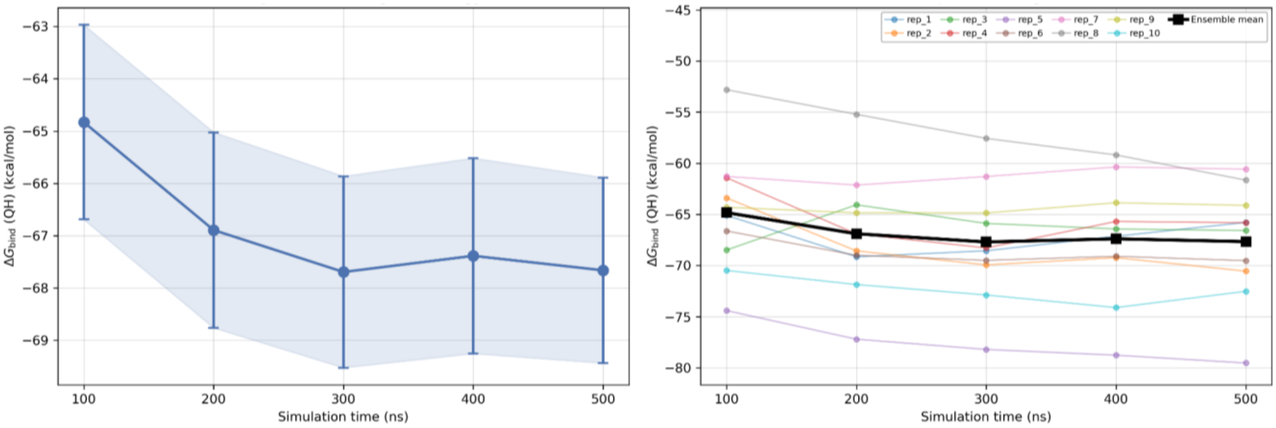


Wild-type:

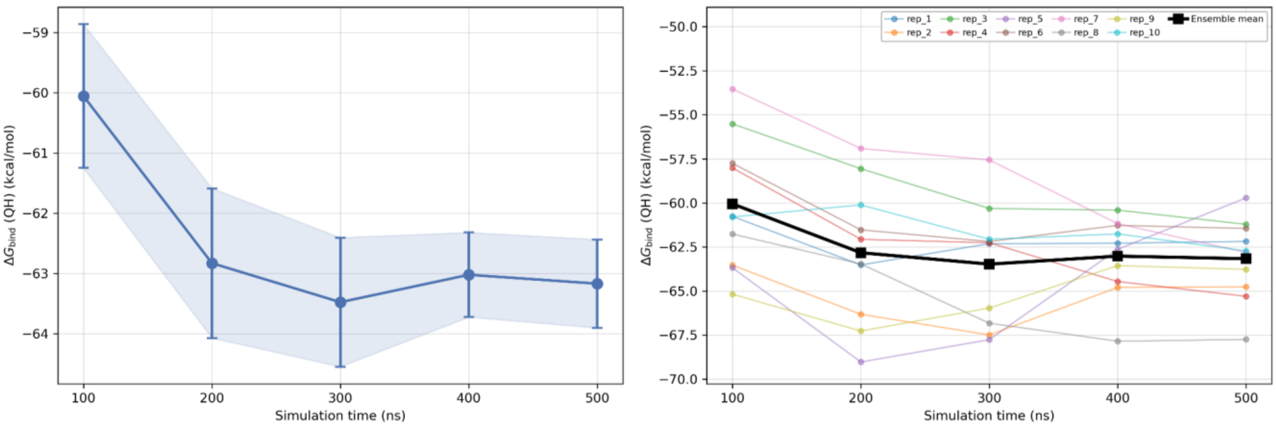

### f) TGFBRII (fs)

Neoantigen:

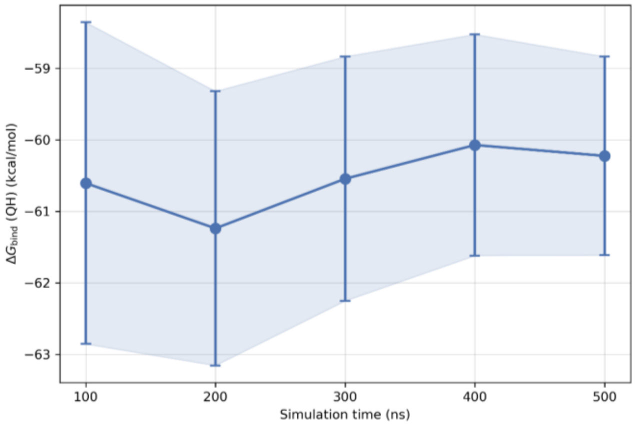


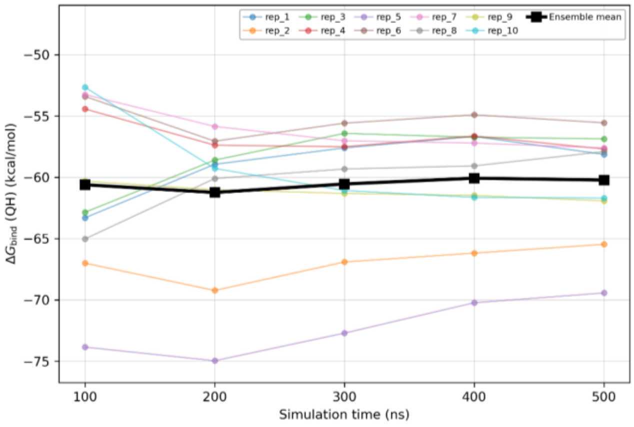


### g) KRAS (G3D)

Neoantigen:

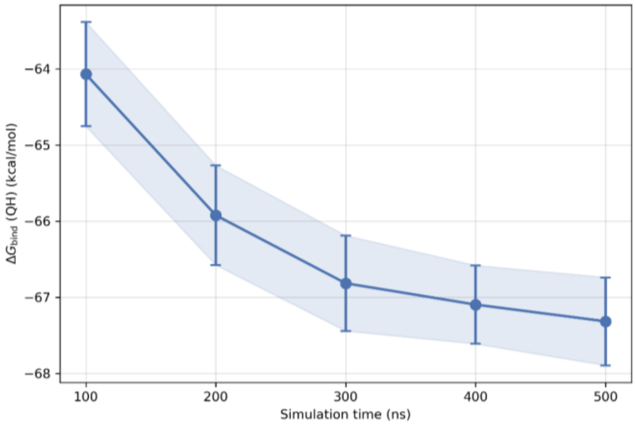


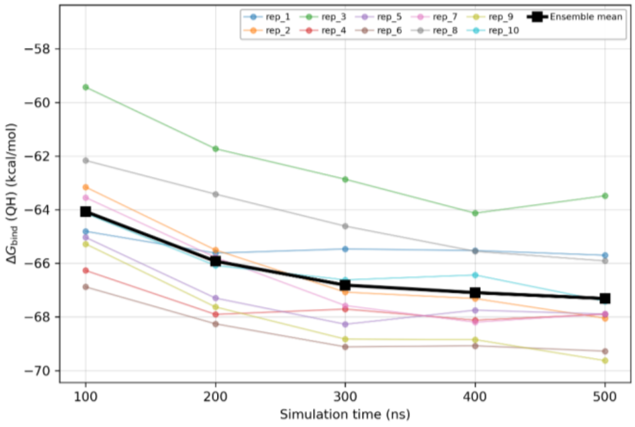


Wild-type:

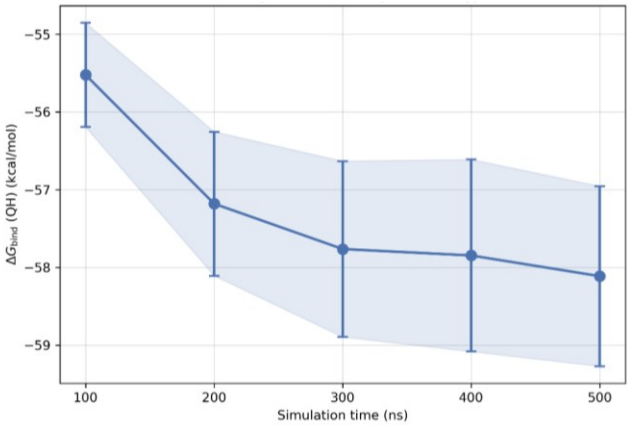


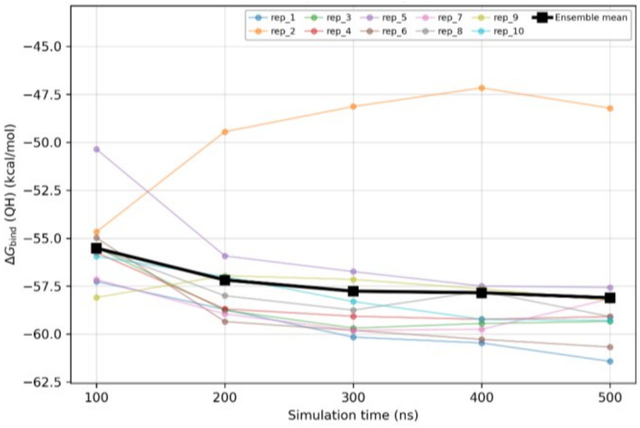

### h) KRAS (G5V)

Neoantigen:

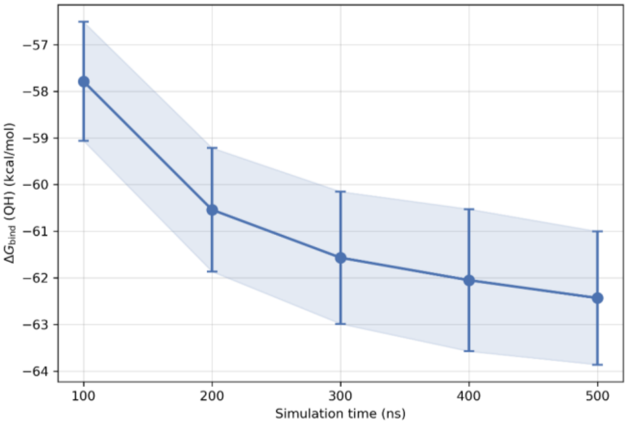


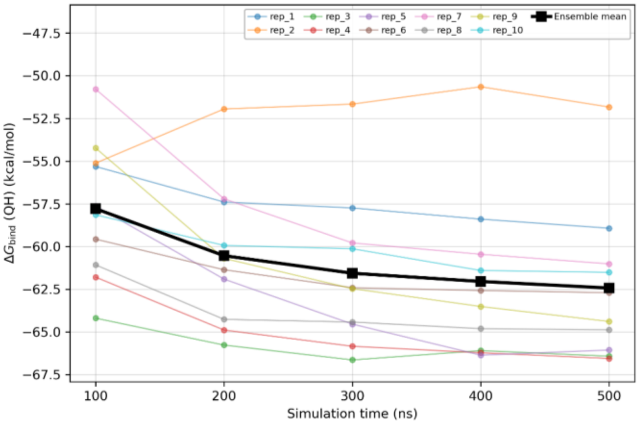


Wild-type:

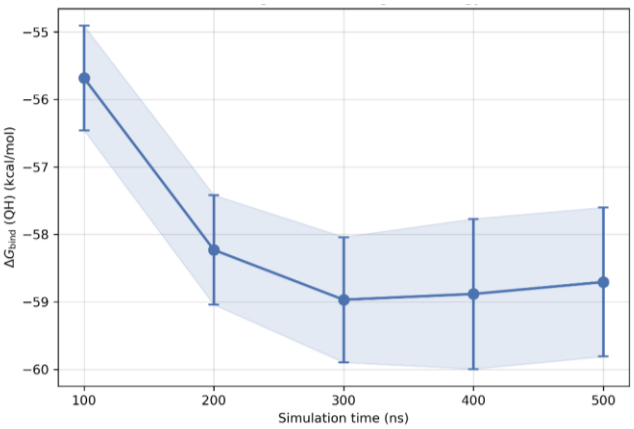


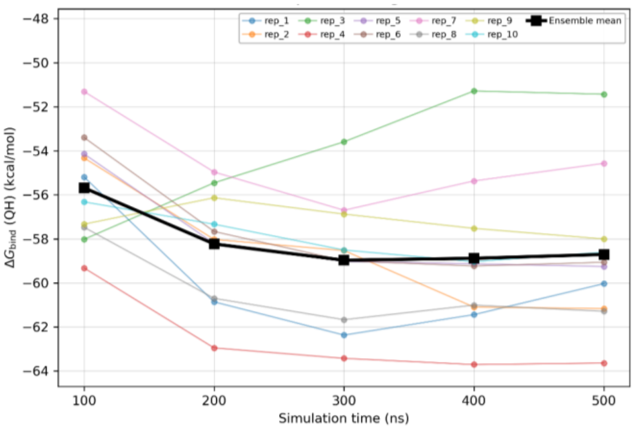


## i) TP53 (R8H)

Neoantigen:

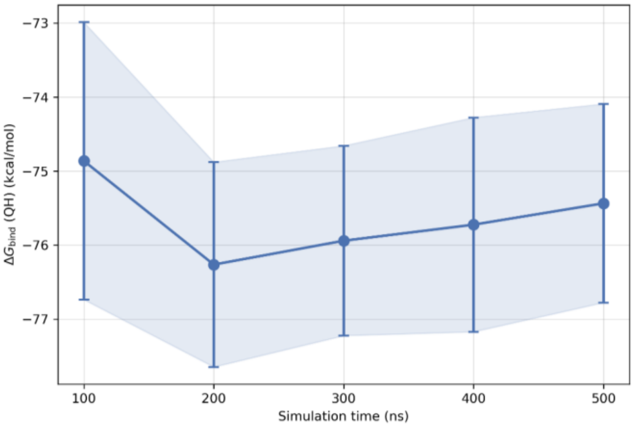


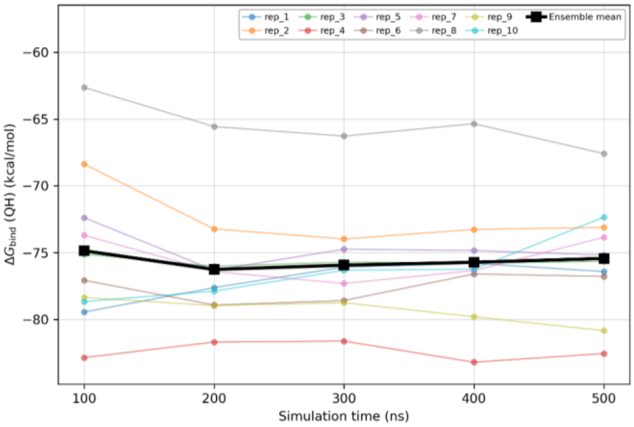

Wild-type:

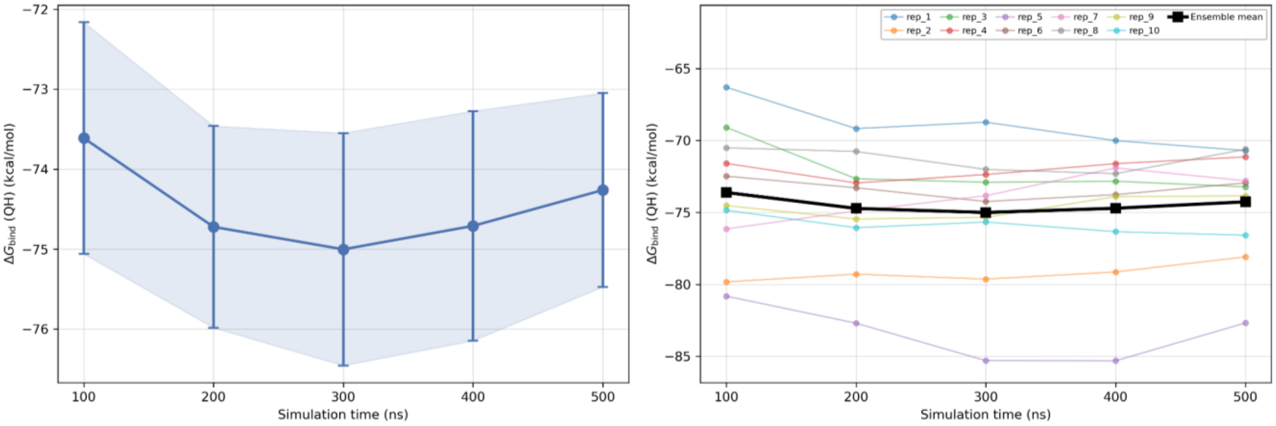


### j) PIK3CA (H2L)

Neoantigen:

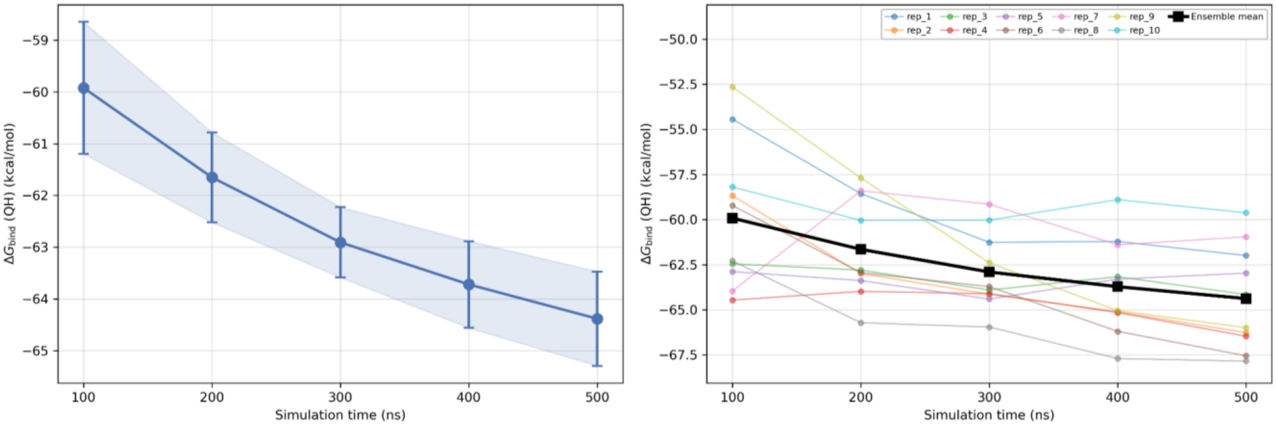


Wild-type:

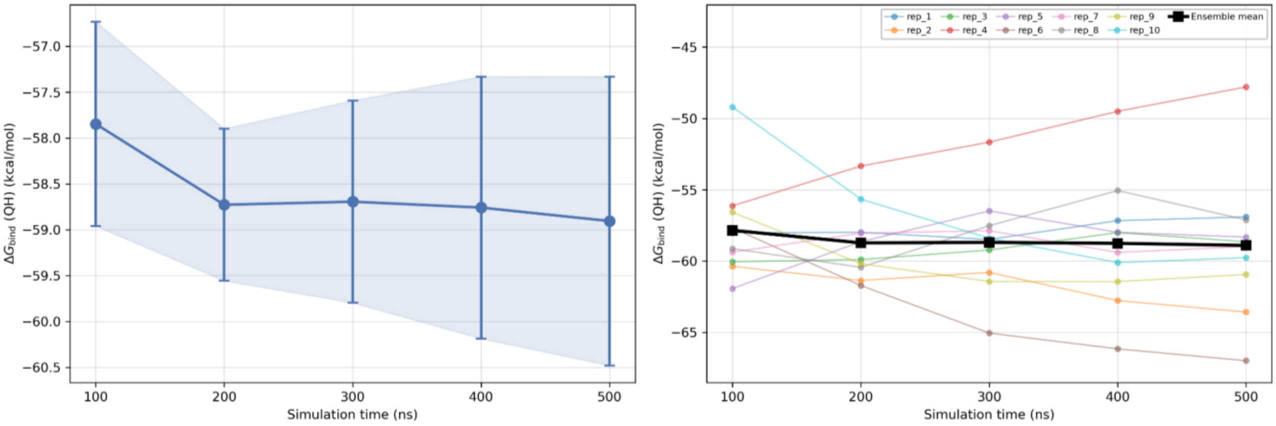

**Figure S2.** Peptide-terminal stability in the HLA binding groove, one multi-panel figure per complex. The upper ten panels correspond to the ten independent replicas of that complex. Each panel plots, against simulation time over the full 500 ns, the minimum heavy-atom distance between each terminal anchor region of the peptide and the HLA binding groove: blue for the N-terminal region and red for the C-terminal region, each region being defined as the terminal residue together with its adjacent neighbor. The horizontal dashed line marks the 5.0 Å distance threshold of the unbinding criterion. The bottom panel shows the corresponding Kaplan–Meier curve for the retention of both termini in the groove across the ten replicas (solid black line), with its 95% Greenwood confidence interval as a grey band; the curve therefore reads as the probability that a replica has not yet experienced a sustained unbinding event at a given time.

### a) FLT3 (D1Y)

Neoantigen:

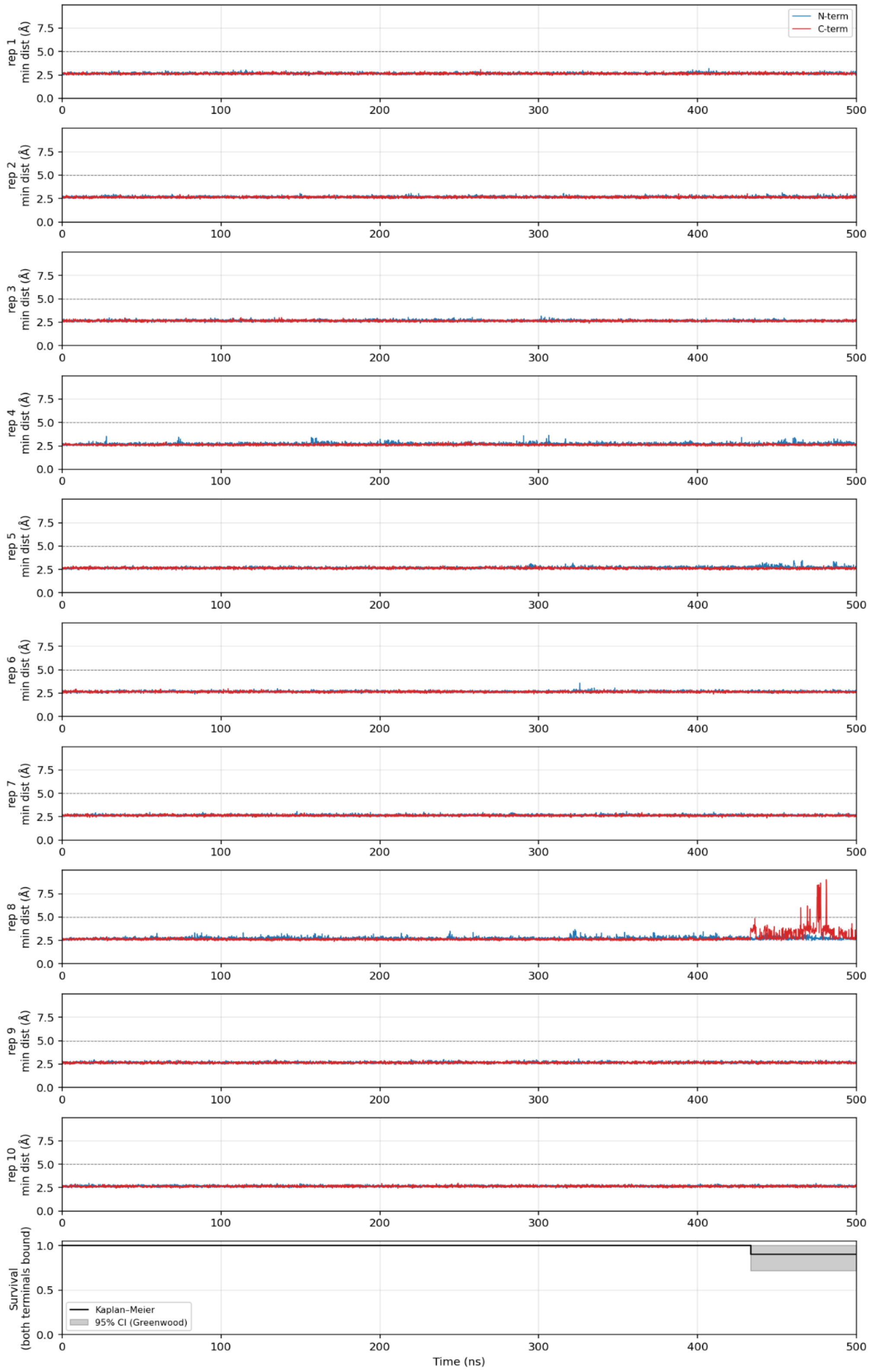

Wild-type:

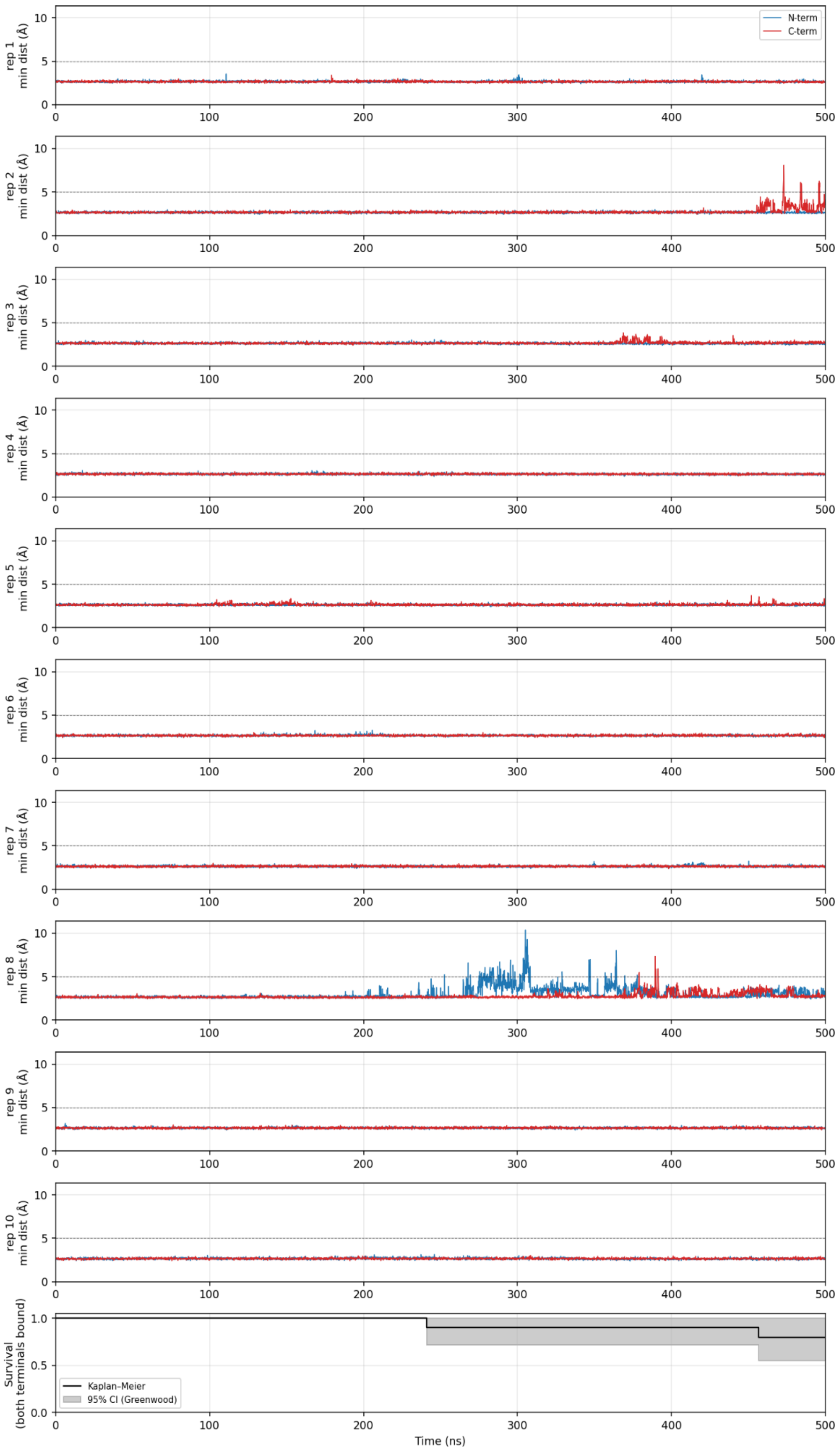

N-term
C-term
rep 1 min dist (Å)
rep 2 min dist (Å)
rep 3 min dist (Å)
rep 4 min dist (Å)
rep 5 min dist (Å)
rep 6 min dist (Å)
rep 7 min dist (Å)
rep 8 min dist (Å)
rep 9 min dist (Å)
rep 10 min dist (Å)
Survival (both terminals bound)
Kaplan–Meier
95% CI (Greenwood)
Time (ns)

**b) APC (fs)**

Neoantigen:

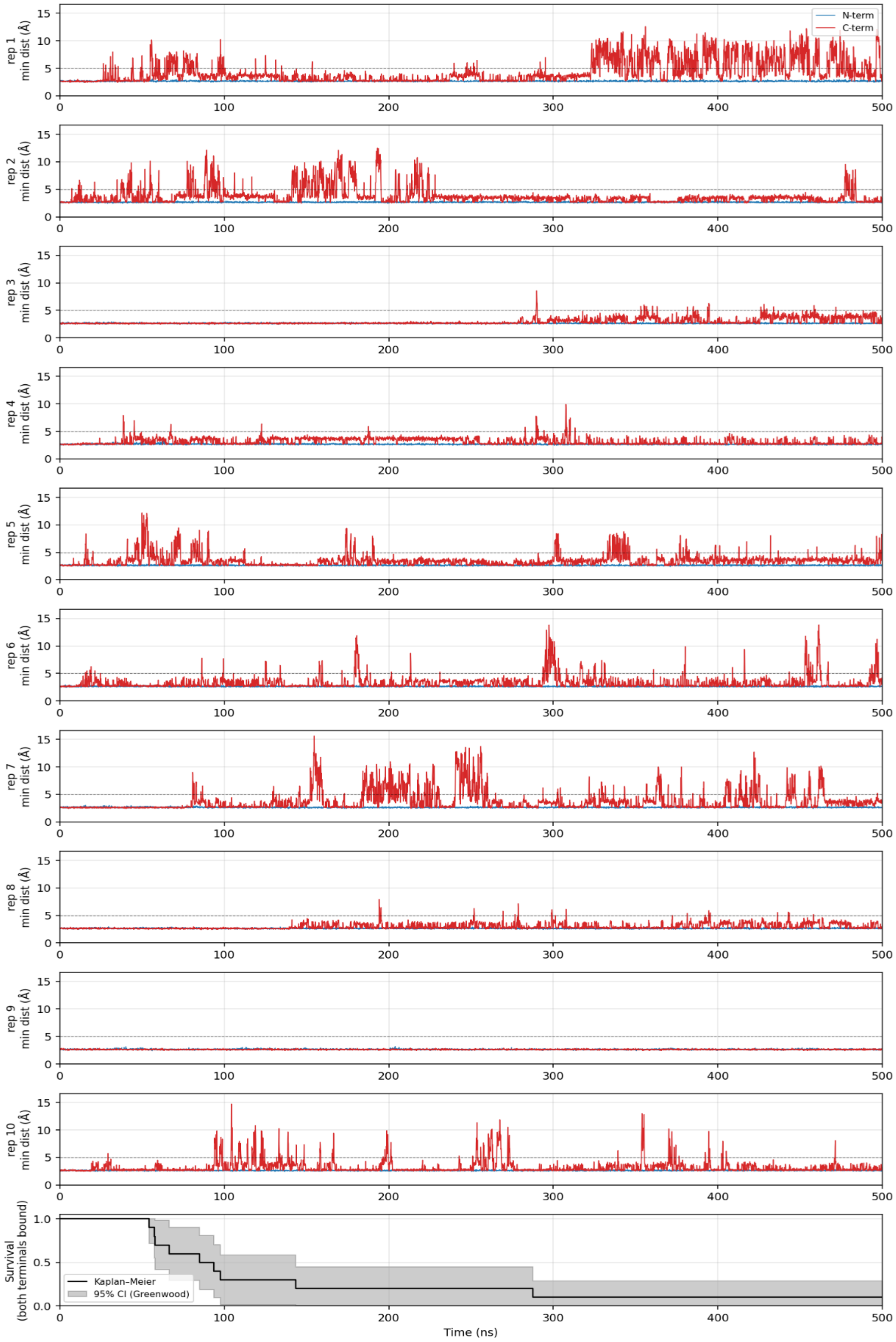

### c) PIK3CA (E11K)

Neoantigen:

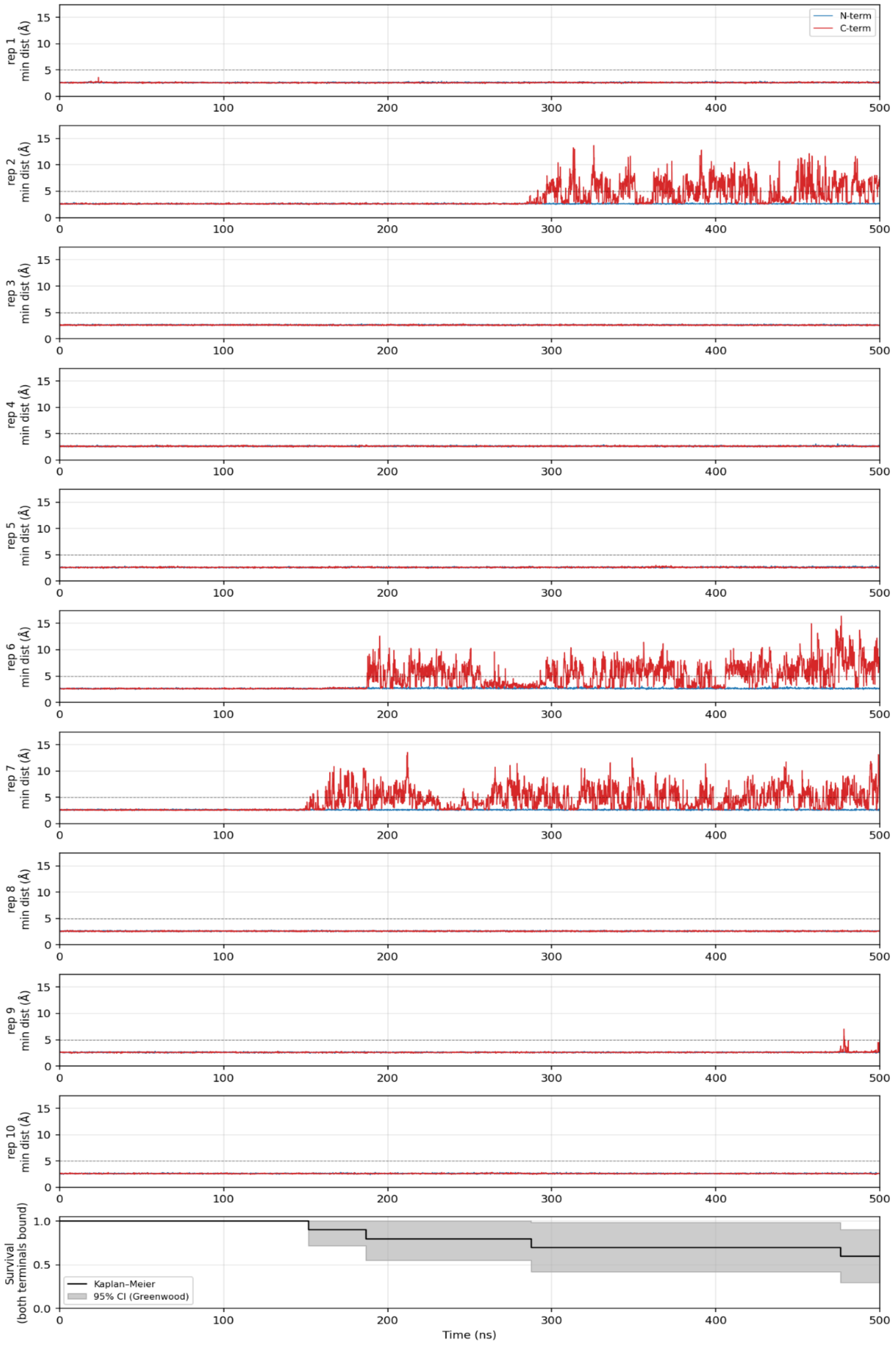

Wild-type:

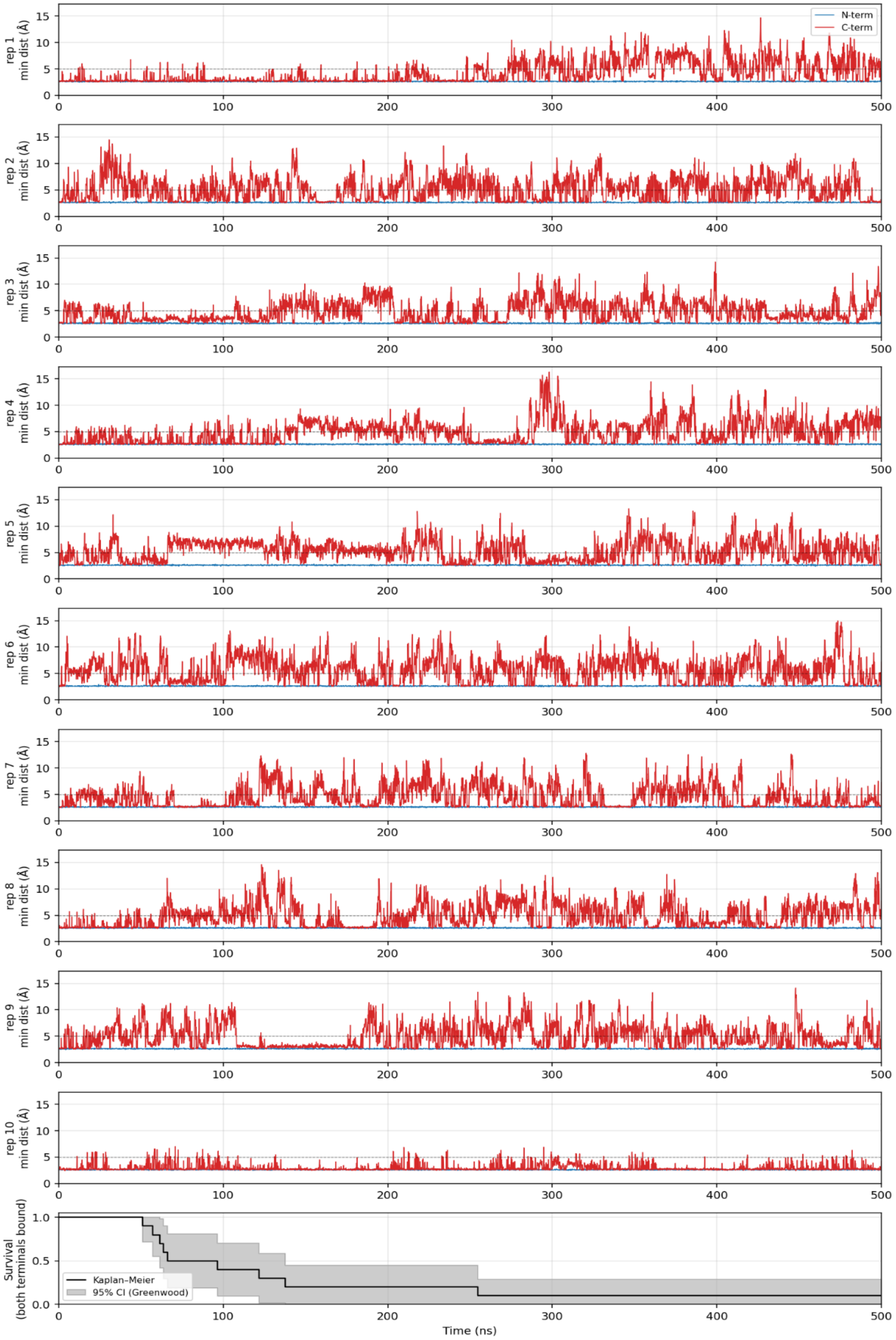

N-term
C-term
rep 1 min dist (Å)
rep 2 min dist (Å)
rep 3 min dist (Å)
rep 4 min dist (Å)
rep 5 min dist (Å)
rep 6 min dist (Å)
rep 7 min dist (Å)
rep 8 min dist (Å)
rep 9 min dist (Å)
rep 10 min dist (Å)
Survival (both terminals bound)
Kaplan–Meier
95% CI (Greenwood)
Time (ns)

### d) NPM1 (fs)

Neoantigen:

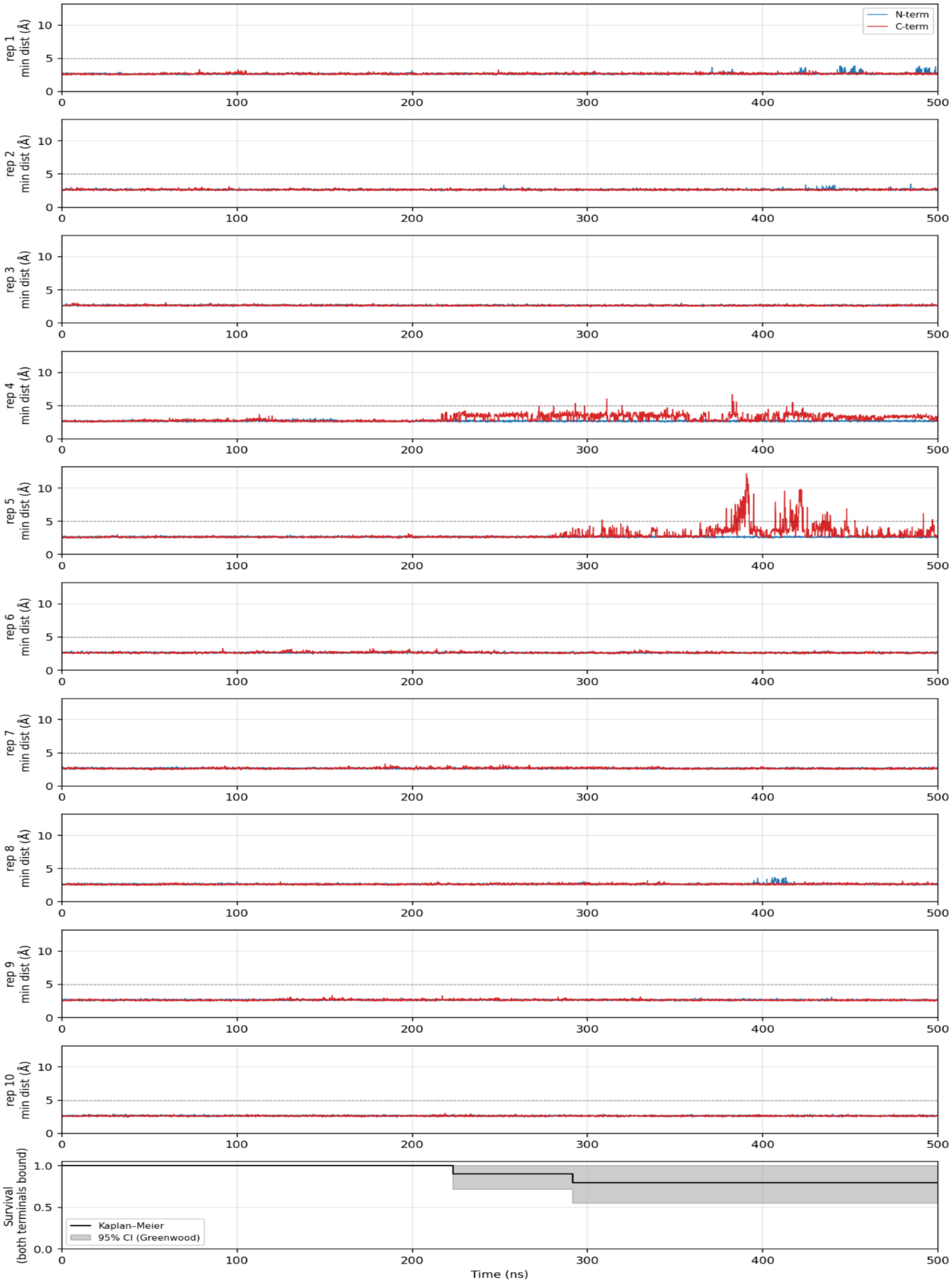

**e) KRAS (G6C)**

Neoantigen:

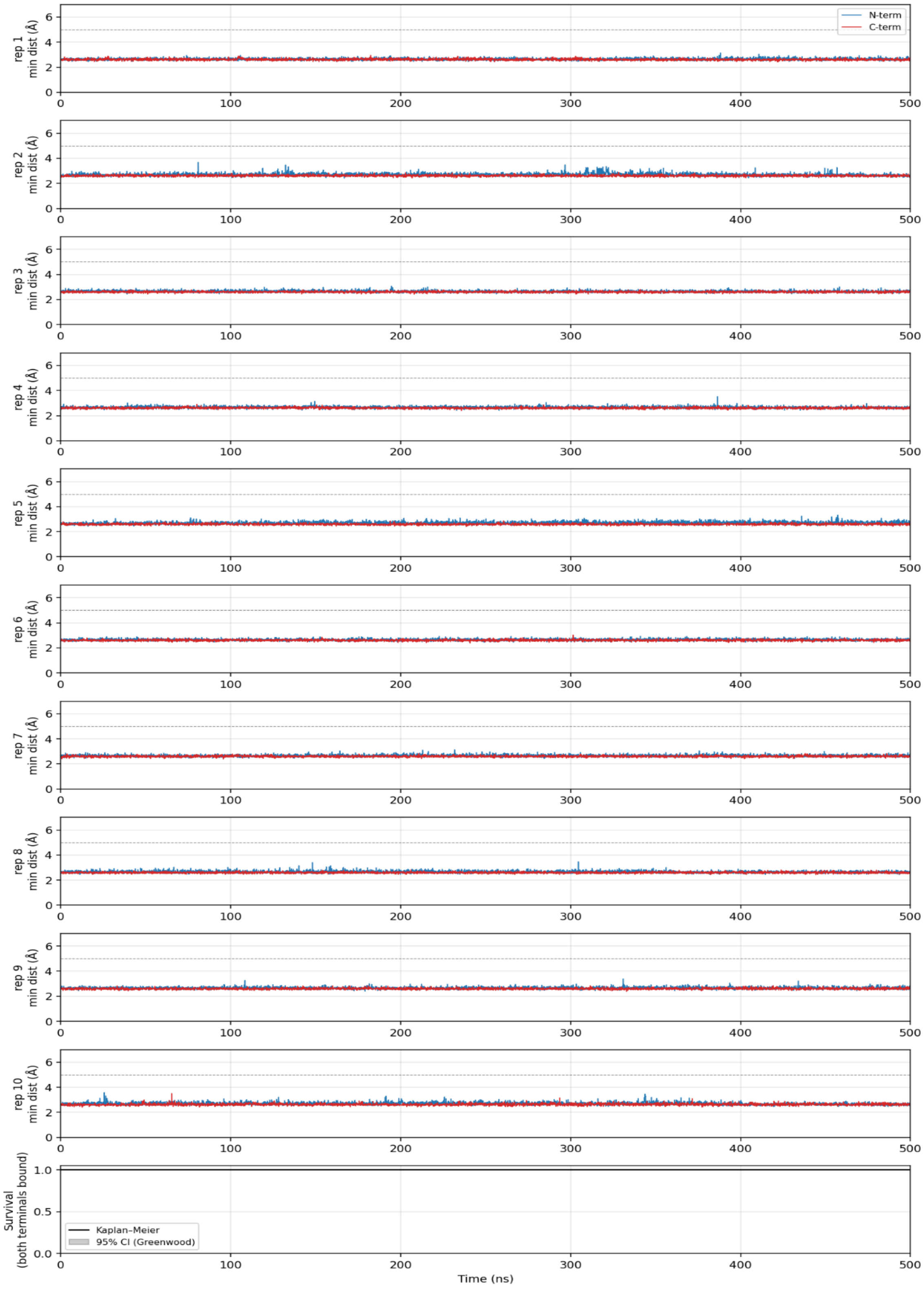

Wild-type:

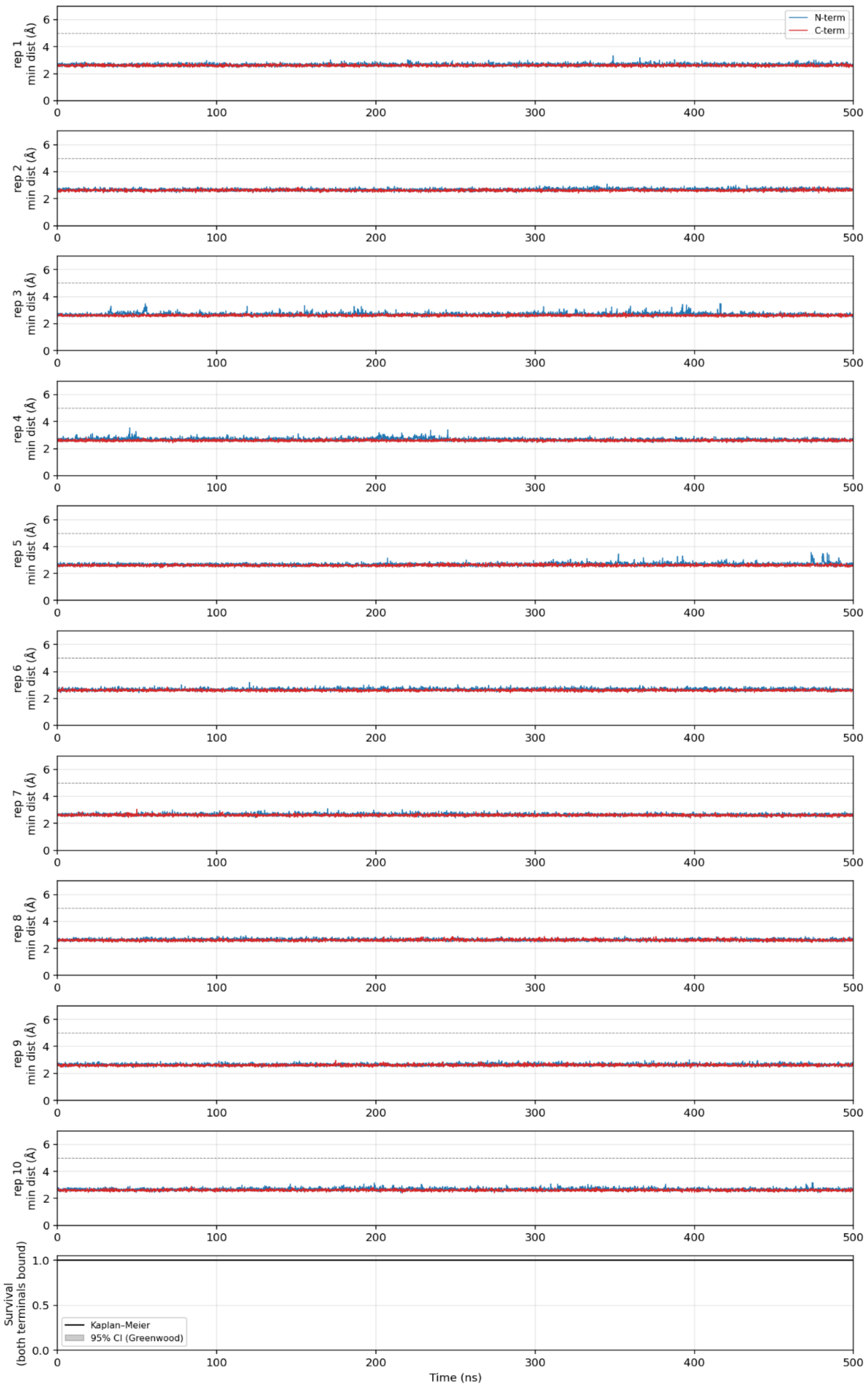
N-term
C-term
rep 1 min dist (Å)
rep 2 min dist (Å)
rep 3 min dist (Å)
rep 4 min dist (Å)
rep 5 min dist (Å)
rep 6 min dist (Å)
rep 7 min dist (Å)
rep 8 min dist (Å)
rep 9 min dist (Å)
rep 10 min dist (Å)
Survival (both terminals bound)
Kaplan–Meier
95% CI (Greenwood)
Time (ns)

### f) TGFBRII (fs)

Neoantigen:

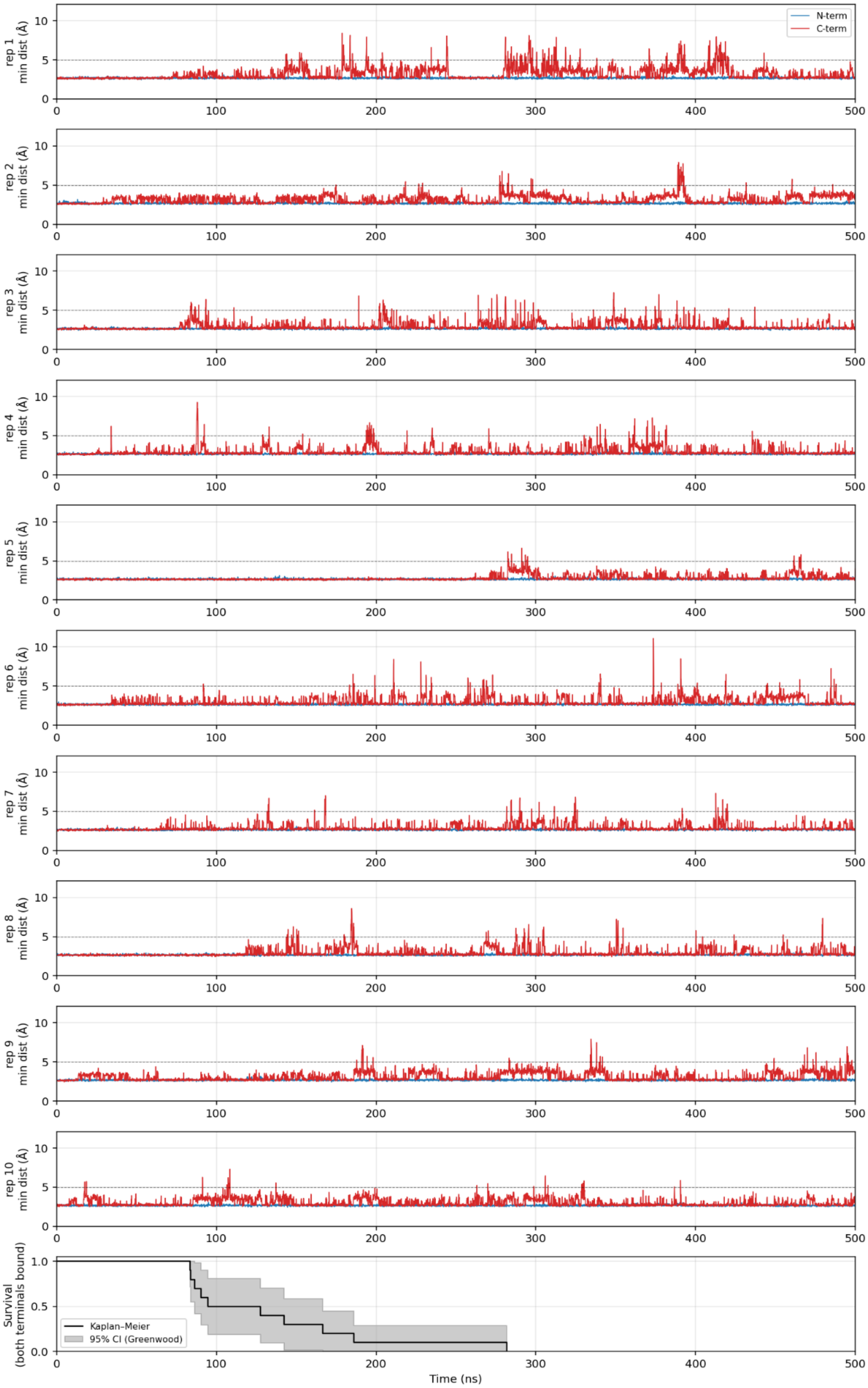

### g) KRAS (G3D)

Neoantigen:

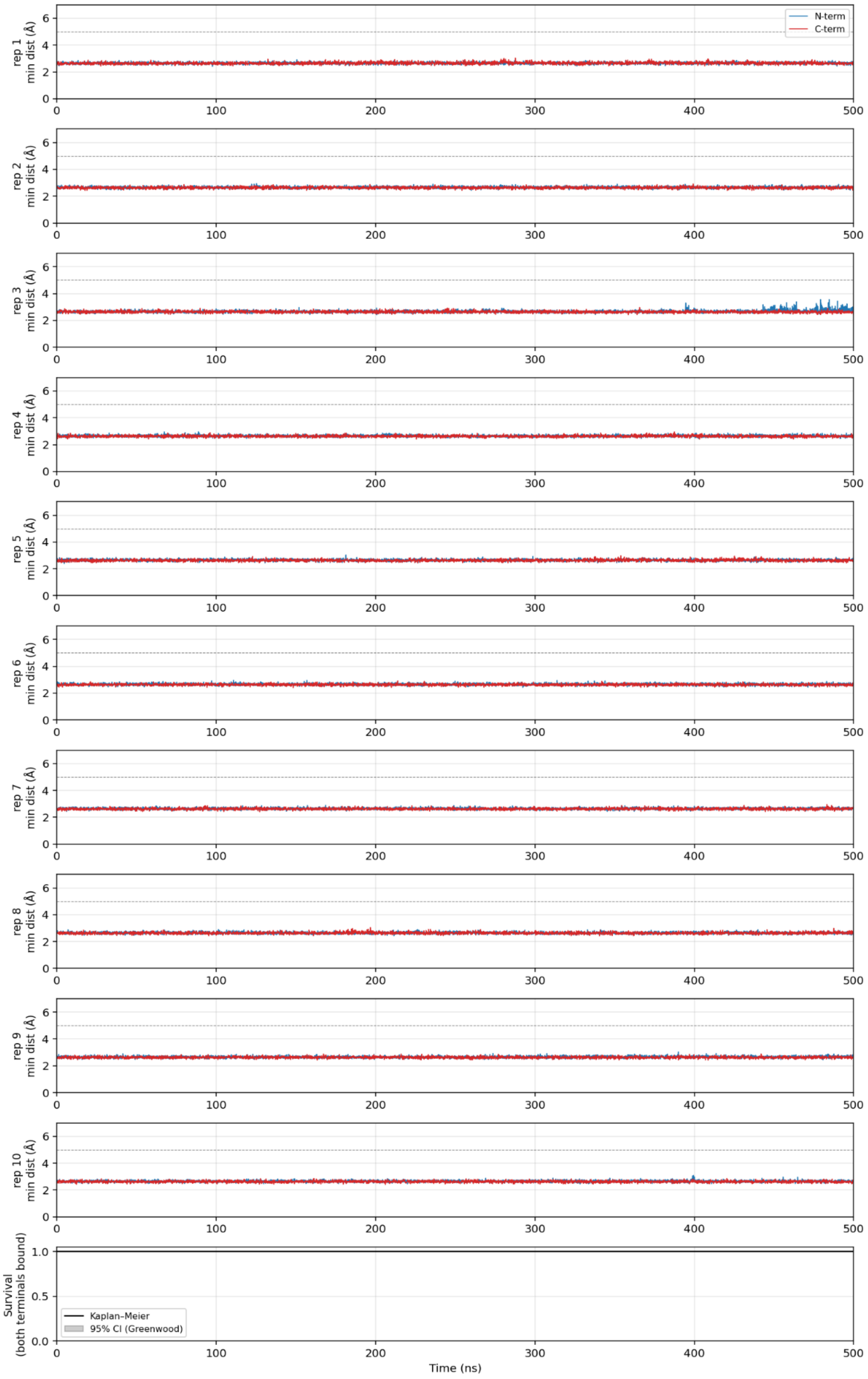

Wild-type:

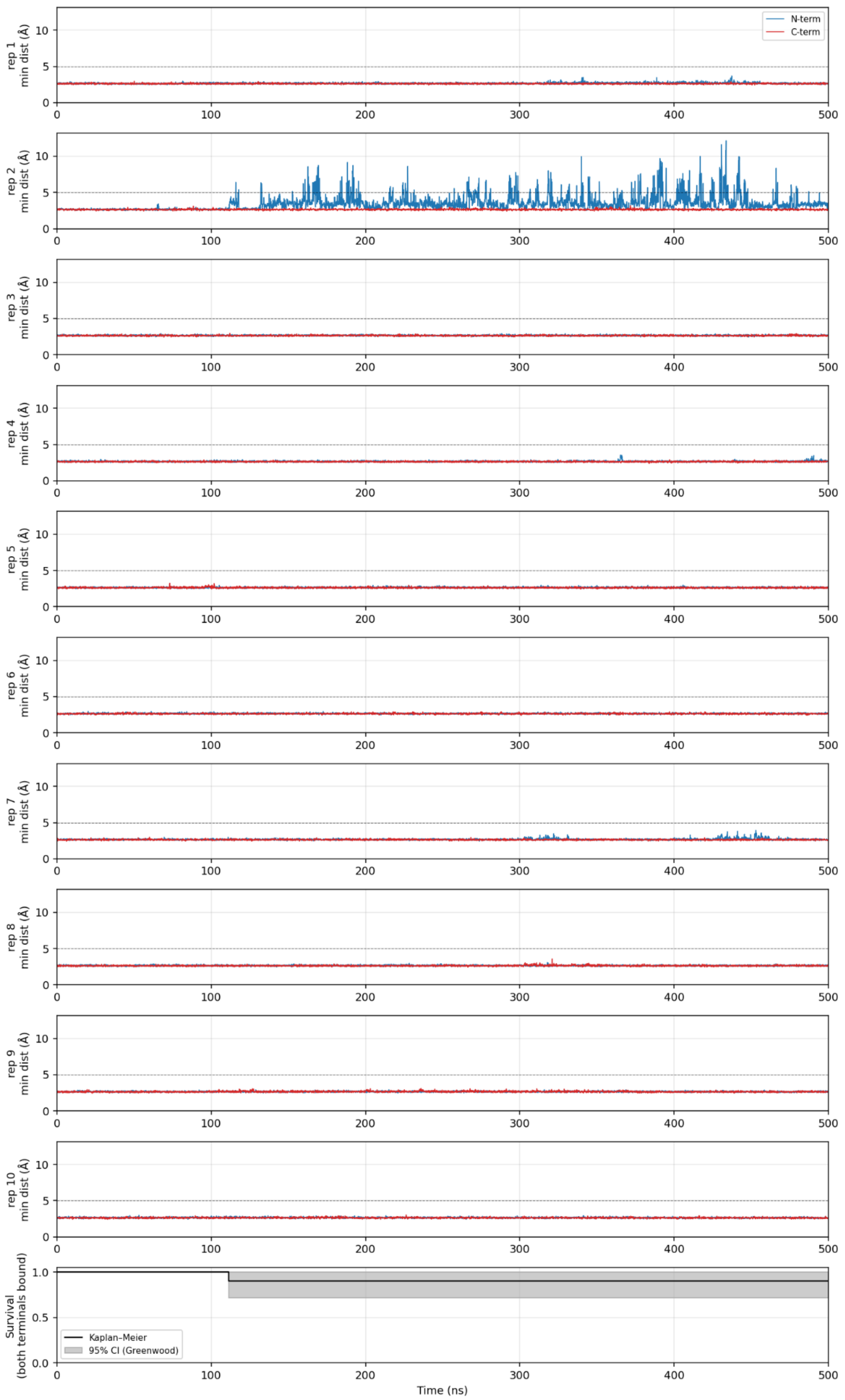
N-term
C-term
rep 1 min dist (Å)
rep 2 min dist (Å)
rep 3 min dist (Å)
rep 4 min dist (Å)
rep 5 min dist (Å)
rep 6 min dist (Å)
rep 7 min dist (Å)
rep 8 min dist (Å)
rep 9 min dist (Å)
rep 10 min dist (Å)
Survival (both terminals bound)
Kaplan-Meier
95% CI (Greenwood)
Time (ns)

### h) KRAS (G5V)

Neoantigen:

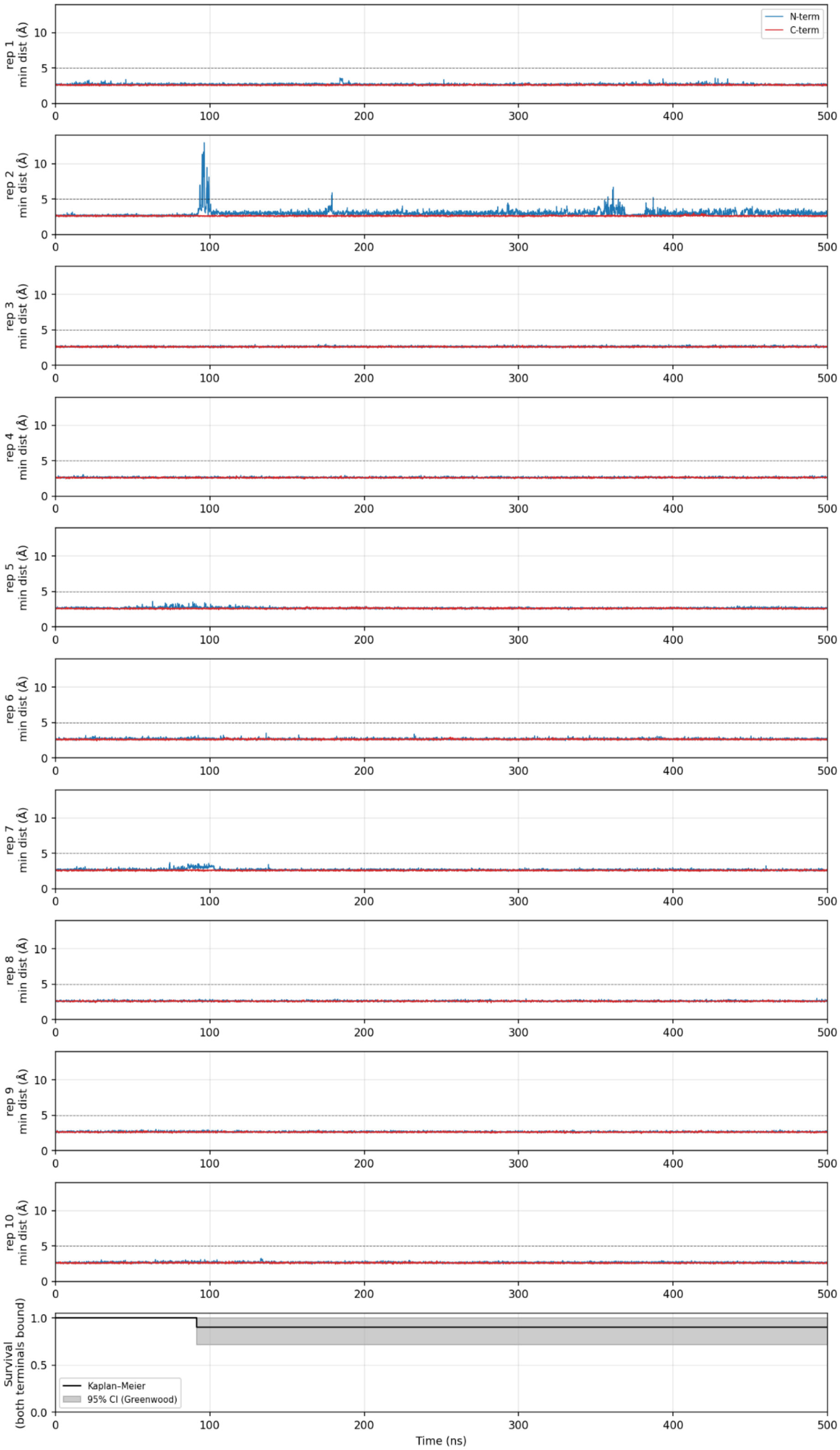

Wild-type:

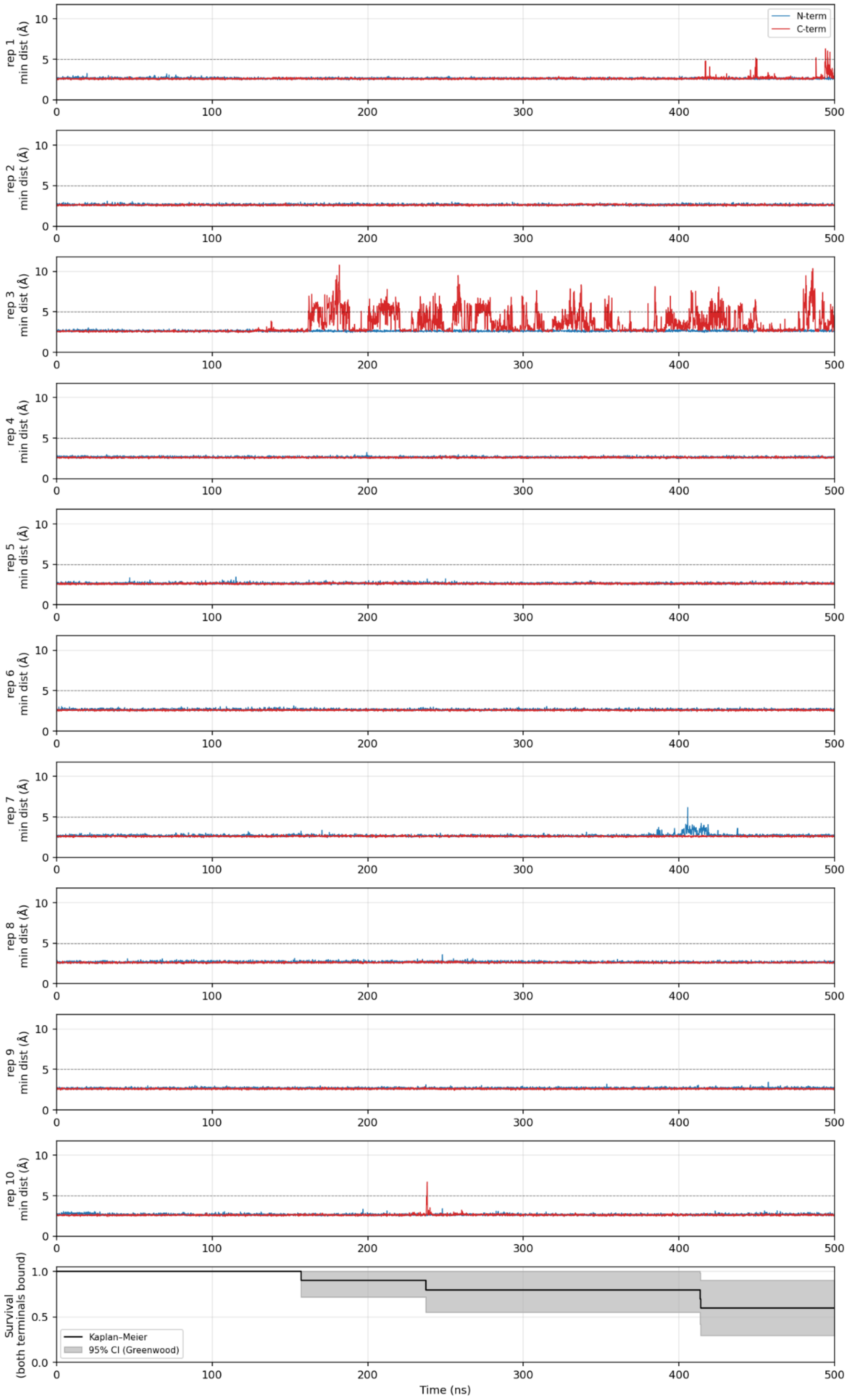
N-term
C-term
rep 1 min dist (Å)
rep 2 min dist (Å)
rep 3 min dist (Å)
rep 4 min dist (Å)
rep 5 min dist (Å)
rep 6 min dist (Å)
rep 7 min dist (Å)
rep 8 min dist (Å)
rep 9 min dist (Å)
rep 10 min dist (Å)
Survival (both terminals bound)
Kaplan–Meier
95% CI (Greenwood)
Time (ns)

## i) TP53 (R8H)

Neoantigen:

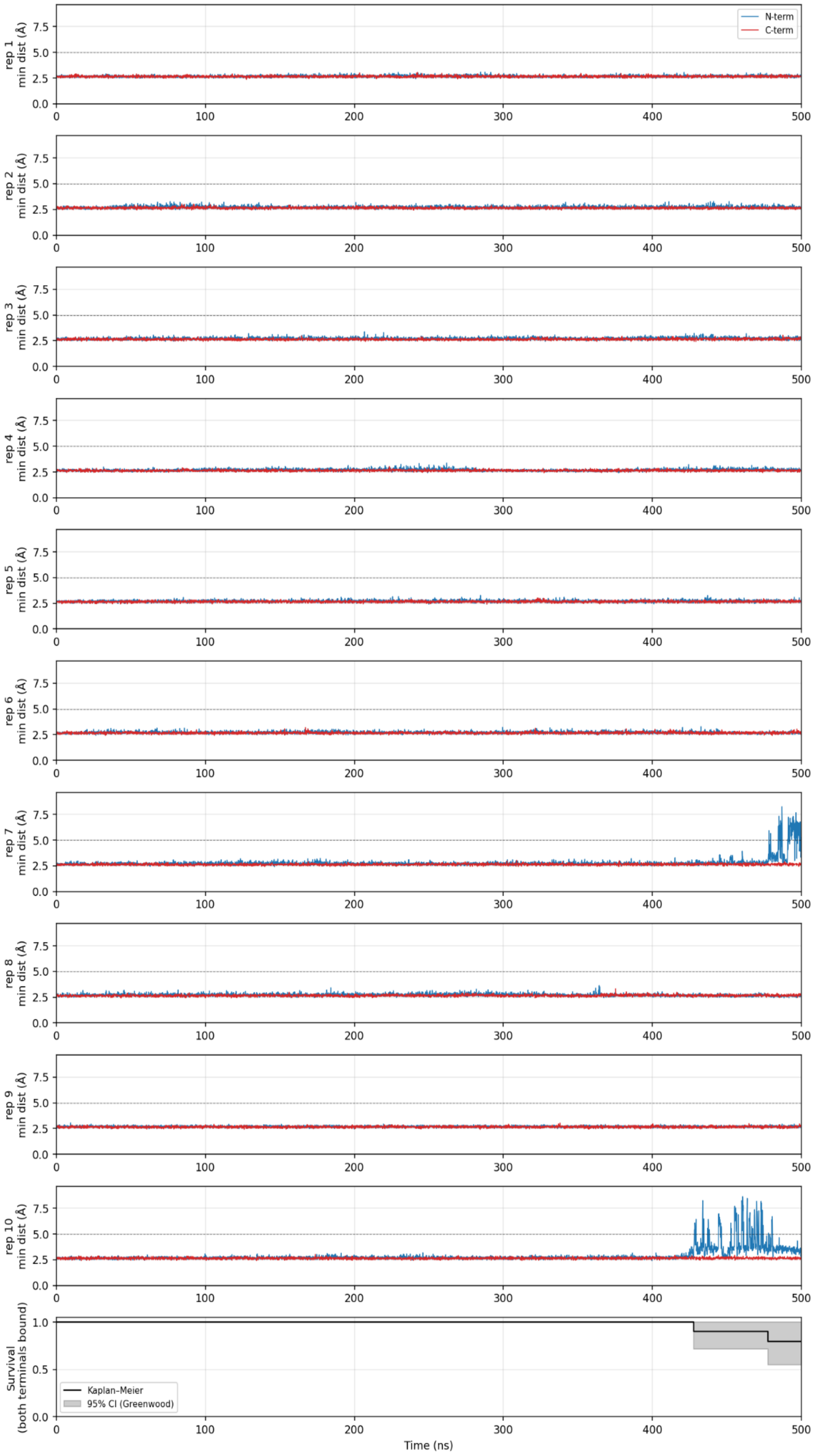

Wild-type:

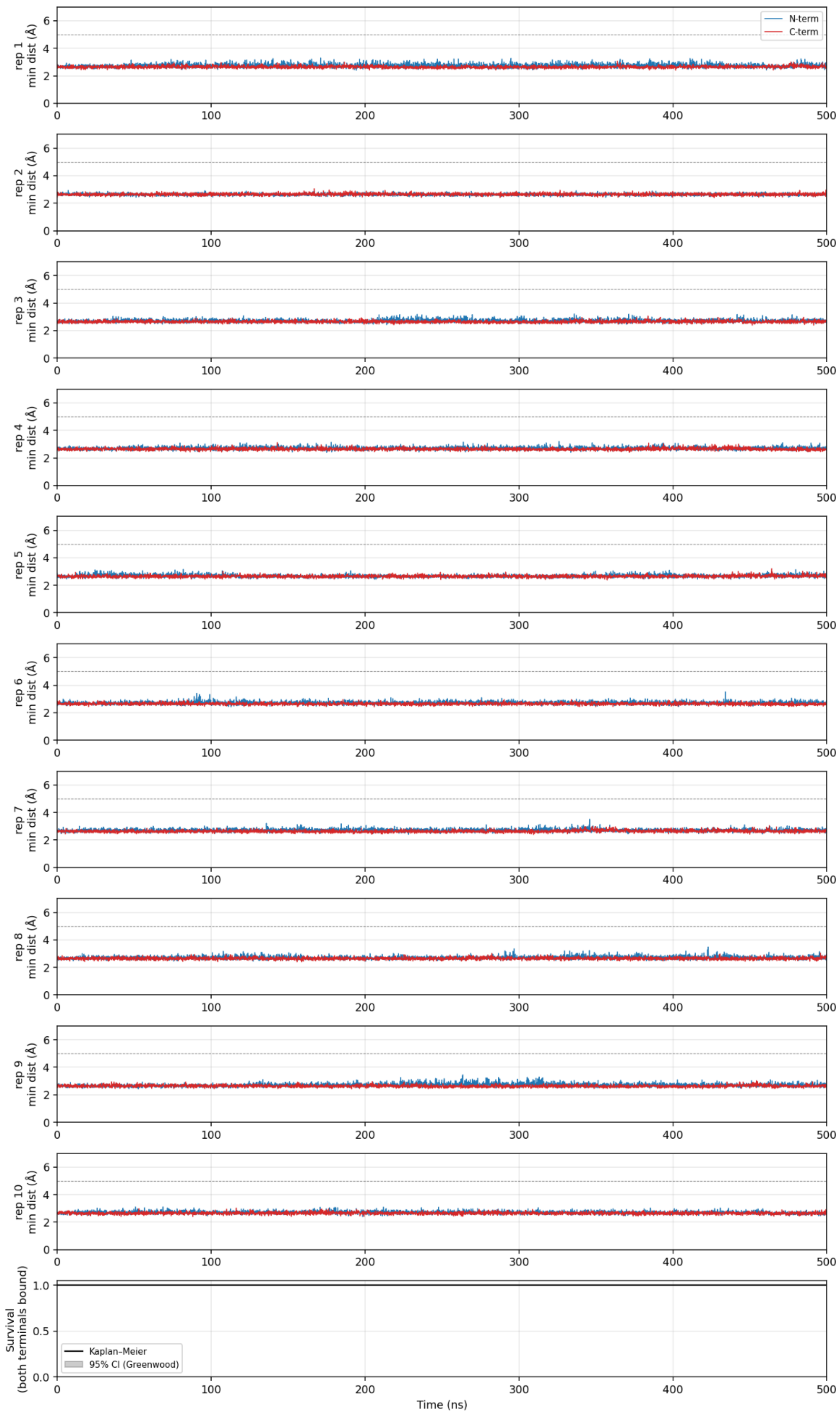
N-term
C-term
rep 1
min dist (Å)
rep 2
min dist (Å)
rep 3
min dist (Å)
rep 4
min dist (Å)
rep 5
min dist (Å)
rep 6
min dist (Å)
rep 7
min dist (Å)
rep 8
min dist (Å)
rep 9
min dist (Å)
rep 10
min dist (Å)
Survival
(both terminals bound)
Kaplan–Meier
95% CI (Greenwood)
Time (ns)

#### j) PIK3CA (H2L)

Neoantigen:

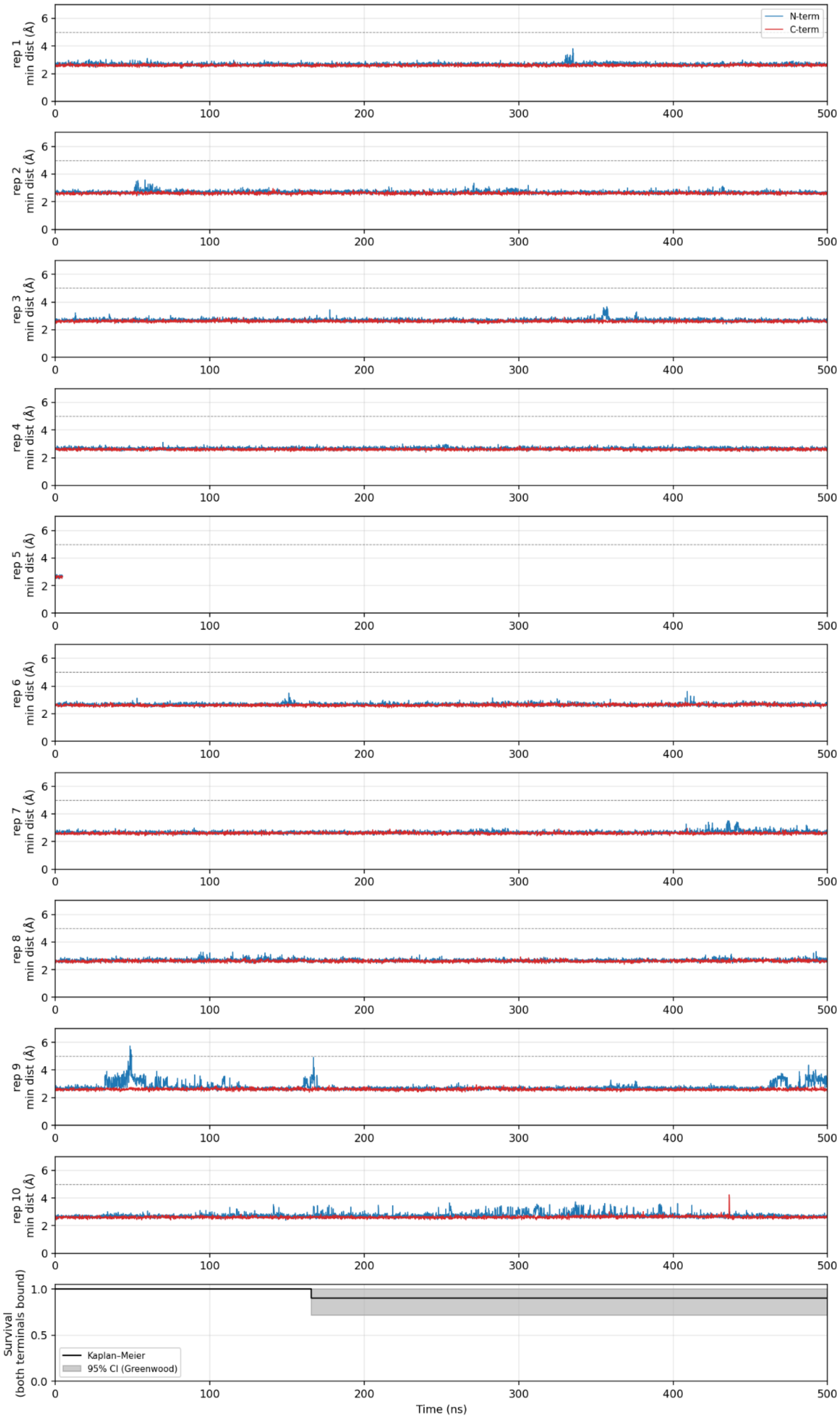

Wild-type:

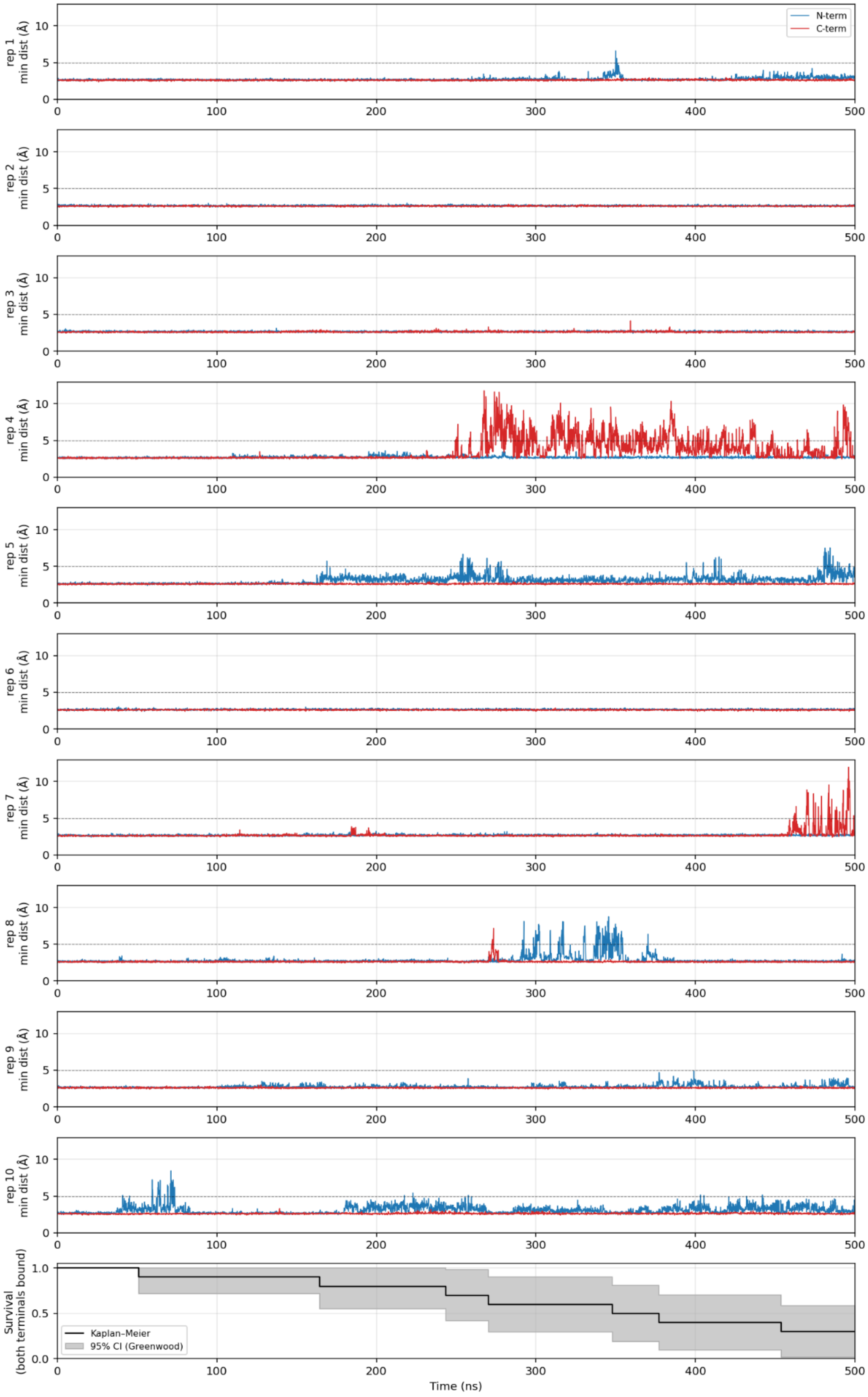

N-term
C-term
rep 1 min dist (Å)
rep 2 min dist (Å)
rep 3 min dist (Å)
rep 4 min dist (Å)
rep 5 min dist (Å)
rep 6 min dist (Å)
rep 7 min dist (Å)
rep 8 min dist (Å)
rep 9 min dist (Å)
rep 10 min dist (Å)
Survival (both terminals bound)
Kaplan–Meier
95% CI (Greenwood)
Time (ns)
